# Endogenous network modeling reveals mechanisms of repair Schwann cell decline and potential recovery targets

**DOI:** 10.64898/2026.05.09.723972

**Authors:** Zongyi Zhou, Ruiqi Xiong, Shunlian Fu, Yang Su, Qiang Ao, Yong-Cong Chen, Ping Ao

**Affiliations:** College of Biomedical Engineering, Sichuan University, Chengdu 610065, China; Chengdu University of Traditional Chinese Medicine, Chengdu 610072, Sichuan, China; Shanghai Center for Quantitative Life Sciences & Department of Physics, Shanghai University 200444, Shanghai, China; National Engineering Research Center for Biomaterials, Sichuan University, Chengdu, Sichuan 610064, China; Shenzhen International Quantum Academy (IQASZ), Shenzhen, Guangdong Province 518048, China

**Keywords:** landscape, peripheral nerve injury, nerve repair, network dynamics, Schwann cell, chronic denervation

## Abstract

Schwann cells, the principal glial cells of the peripheral nervous system, play a central role in nerve repair following injury. Upon injury, mature Schwann cells dedifferentiate into repair Schwann cells. These processes are governed by complex gene regulatory networks, yet the quantitative dynamics of these processes remain unclear. Here, using a bottom-up systems biology approach, we constructed an endogenous regulatory network model based on experimentally validated interactions, without relying on high-throughput data as input. The model captures Schwann cell dedifferentiation dynamics and reveals a potential landscape composed of stable states and intermediate transition states. Simulations recapitulate post-injury trajectories and confirm the role of c-Jun upregulation in maintaining repair capacity. Furthermore, the model predicts multiple potential therapeutic targets, including P53, JNK, and PTEN, for sustaining repair competence. We also identify intrinsic heterogeneity within repair Schwann cells and uncover key transition states that simultaneously connect repair-competent cells to both repair-deficient and apoptotic phenotypes, indicating that these intermediate states may represent critical regulatory bottlenecks and key cellular targets for improving the success of peripheral nerve regeneration. Overall, this work provides new insights into the precise regulation of Schwann cell fate and establishes a theoretical framework for regenerative medicine and clinical strategies in peripheral nerve repair.

## Introduction

Upon peripheral nerve injury, mature Schwann cells dedifferentiate into a transient repair phenotype that actively supports axonal regeneration (Napoli et al., 2012;Monk et al., 2015). The remarkable regenerative capacity of peripheral nerves is largely attributed to this plasticity of Schwann cells. Following injury, Schwann cells undergo a phenotypic transformation into a specialized reparative state, characterized by the downregulation of myelin-associated genes and the upregulation of pro-regenerative genes, including those encoding neurotrophic factors and cytokines involved in immune modulation. Concurrently, Schwann cells initiate myelin autophagy and recruit macrophages to facilitate the efficient clearance of myelin debris. They also reorganize into Büngner bands, which provide structural guidance for regenerating axons. Upon completion of regeneration, repair Schwann cells redifferentiate into either mature myelinating Schwann cells or non-myelinating Schwann cells, thereby restoring the functional architecture of the nerve (Jessen and Mirsky, 2016;2019).

Despite these reparative mechanisms, functional recovery following peripheral nerve injury remains suboptimal, a limitation largely driven by the inherently slow axonal regrowth and the transient reparative phenotype of Schwann cells. Over time, repair Schwann cells progressively lose their regenerative capacity, characterized by the downregulation of repair-associated genes and reduced expression of neurotrophic factors, which contribute to the failure of nerve regeneration. (Höke, 2006;Sulaiman and Gordon, 2009;Jessen and Mirsky, 2019). Chronic denervation of Schwann cells should be considered not only a therapeutic target but also a central objective of any effective strategy aimed at improving nerve regeneration. How to modulate these specialized Schwann cells to boost their reparative capacity and prevent functional decline over the protracted axonal regeneration period remains a critical research focus (Jessen and Mirsky, 2019;Nocera and Jacob, 2020;McMorrow et al., 2022).

In recent years, numerous studies have revealed the critical roles of various transcription factors and signaling pathways in Schwann cell responses to injury. Previous research has systematically delineated the major transcriptional regulatory networks involved in Schwann cell myelination, including the roles of key factors such as Sox10, Oct6/Brn2, and Krox20 in activating myelin-associated genes (Svaren and Meijer, 2008). In addition, studies have shown that sustained expression of Sox2 in mice can inhibit myelination while inducing a proliferative, undifferentiated state (Roberts et al., 2017). In Stat3 knockout experiments in mice, researchers observed a significant reduction of Schwann cells following denervation, indicating that this gene is essential for the long-term maintenance of Schwann cell phenotypes (Benito et al., 2017). Similarly, c-Jun knockout experiments demonstrated that loss of c-Jun led to substantial neuronal cell loss, impaired axonal regeneration, and reduced secretion of multiple neurotrophic factors. c-Jun has also been identified as a regulator of GDNF and BDNF (Fontana et al., 2012). Collectively, these studies highlight the presence of numerous key transcription factors involved in Schwann cell regulation. Such traditional approaches are largely based on a linear regulatory paradigm, focusing on the role of individual genes or single signaling pathways in specific biological processes. As a result, they primarily explain the function and importance of these factors in nerve regeneration in an isolated manner.

However, such locally focused approaches are insufficient to fully elucidate system-level regulatory mechanisms. As illustrated by the “radio” analogy proposed by Lazebnik, biologists often demonstrate that the loss of a specific factor causes the system to “fall silent,” thereby identifying it as a key target and assuming that modulating this single component may be sufficient to “repair” the system (Lazebnik, 2002). Similarly, Davidson pointed out that observations at the level of individual genes cannot reveal the overall organizational structure of biological systems or the mechanisms underlying organ development, but instead provide only a microscopic and inherently limited perspective (Davidson, 2010). In the context of chronic denervation–induced systemic functional decline, the loss of repair capacity in Schwann cells is not determined by any single factor alone, but rather arises from dysregulated coordination among key molecules and signaling pathways within the regulatory network. Therefore, it is necessary to analyze these biological phenomena from a systems-level perspective.

In recent years, with the rapid development of high-throughput sequencing technologies, researchers have begun to systematically analyze gene expression profiles of Schwann cells in different states using transcriptome sequencing and single-cell RNA sequencing, thereby constructing their potential gene regulatory networks (Arthur-Farraj et al., 2017;Brosius Lutz et al., 2022). Using high-resolution omics, investigators analyzed human Schwann cells, fibroblasts, and neural tissues at both the protein and transcript levels, revealing that cultured human Schwann cells retain their inherent reparative capacity (Weiss et al., 2016). Although these studies have characterized the molecular features of Schwann cells during nerve repair at a systems level and provided valuable references for further elucidation of their regulatory mechanisms, such high-throughput data mainly provide a descriptive characterization of system states and are limited in revealing the underlying mechanisms and causal relationships driving these changes.

The fate determination and functional remodeling of Schwann cells depend on complex interactions among multiple key molecules and signaling pathways. Correlation-based analyses alone are insufficient to elucidate their underlying regulatory logic and dynamic evolution. It is therefore necessary to introduce research frameworks grounded in cell biology and evolutionary biology that can capture causal relationships and system-level dynamics. Such approaches allow for understanding the interaction mechanisms among these critical regulatory factors at the network level and further reveal the intrinsic drivers of Schwann cell state transitions and changes in their repair capacity (Kitano, 2002;Maizels and Briscoe, 2026).

Indeed, Schwann cell dedifferentiation is governed by a highly interconnected regulatory network involving extensive cross-talk among transcription factors and signaling pathways, which cannot be captured by simplified models, highlighting the need for a system-level network framework (Nocera and Jacob, 2020). Conceptual foundations for such approaches trace back to the epigenetic and fitness landscapes proposed by Waddington and Wright, which provided topological representations of biological dynamics (Wright, 1932;Waddington, 2014). Kauffman later demonstrated that gene regulatory networks inherently converge to stable attractor states using stochastic Boolean models (Kauffman, 1969). Building on these ideas, the first quantitative landscape was constructed for the bacteriophage lambda gene switch (Zhu et al., 2004), leading to the endogenous network theory, which views molecular regulatory systems as nonlinear stochastic dynamical systems with multiple steady states corresponding to distinct cellular states, including disease states (Ao et al., 2008). This framework has since been successfully applied to multiple cancers and to developmental patterning of the telencephalon (Yuan et al., 2017;Sun et al., 2024;Yao et al., 2024).

Building on existing experimental evidence, we constructed an endogenous core regulatory network of Schwann cells in the context of peripheral nerve injury from a systems-level perspective. Dynamic modeling of this regulatory network revealed multiple steady states and transitional states, which are interconnected to form a dedifferentiation landscape that captures the topological structure of Schwann cell state transitions and elucidates the molecular processes underlying chronic denervation. Furthermore, we simulated targeted interventions within the network and successfully recapitulated experimentally validated targets required for maintaining the reparative activity of Schwann cells, while also predicting several additional candidate therapeutic targets.

## Materials and methods

### Construction of endogenous molecular-cellular network

In 2008, Ao and Hood *et al*. proposed an endogenous network model that employs nonlinear stochastic dynamical systems to describe the molecular regulatory mechanisms of complex biological systems within the framework of systems biology (Ao et al., 2008). This theory suggests that, over the course of biological evolution, molecules and cytokines within an organism, including tumor suppressor genes, oncogenes, and associated growth factors, collectively constitute a nonlinear and stochastic endogenous molecular-cellular network at the cellular level. This endogenous network can be quantitatively represented by the expression levels of key endogenous factors, forming a high-dimensional stochastic dynamical system. The nonlinear dynamic interactions among network components can generate multiple local steady states with either distinct or indistinct biological functions, which may correspond to specific cellular states observed experimentally. The endogenous network can autonomously form and sustain multiple steady states that persist over extended periods of time. These steady states collectively constitute the potential landscape, and the transitions of cellular states during differentiation can be interpreted as stochastic shifts between distinct steady states within this landscape. Compared with the prevalent linear additive thinking in traditional research, the global dynamic characteristics of the network provide a more comprehensive framework for understanding the mechanisms underlying disease onset and progression (Yuan et al., 2017).

To construct an endogenous network for a specific research object, the first step is to select modules and pathways according to biological phenomena and functions. These modules are both integral components of, and relatively independent from, the entire biological system. Interactions occur among key endogenous factors within each module, while higher-level functions, such as cell fate determination, can be achieved through crosstalk between modules. This hierarchical organization resembles the principle of modular organization (Hartwell et al., 1999). Based on the sequential responses of Schwann cells to peripheral nerve injury, we identified four key modules and three major pathways, including the Schwann cell differentiation, inflammation, proliferation, and apoptosis modules, as well as the JNK, Akt, and Ras/ERK pathways. Detailed information on all modules and their constituent factors of the endogenous network model is provided in Table S1 in Part A of the supplementary materials. The criteria for selecting modules and core factors in the endogenous network are described below.

### Selection criteria for modules and core factors in the endogenous network

Certain cellular functions, such as signal transduction, are carried out by functional modules within the cell. These modules consist of interacting molecules that regulate each other, and integrating insights from biological phenomena with molecular-level studies can facilitate a deeper understanding of their underlying mechanisms (Hartwell et al., 1999). The construction of endogenous networks is grounded in biological phenomena(Ao et al., 2008). First, we select modules based on observed cellular or tissue-level phenomena and their associated functions. Following peripheral nerve injury, Schwann cells undergo significant morphological and functional changes. According to current experimental evidence, they dedifferentiate from myelinating and non-myelinating Schwann cells into a flattened repair phenotype. Accordingly, a Schwann cell differentiation module was incorporated into the network (Jessen and Arthur-Farraj, 2019).

During Wallerian degeneration, Schwann cells release pro-inflammatory cytokines and recruit macrophages to cooperatively clear myelin debris, thereby creating a favorable environment for nerve regeneration. Based on this process, an inflammation module was included (Martini et al., 2008;Jessen and Mirsky, 2016).

In response to injury, Schwann cells exit the quiescent state and re-enter the cell cycle, leading to proliferation that fills the injured area and contributes to the formation of Bands of Büngner. Therefore, a cell proliferation module was added (Kim et al., 2000;Kitaura et al., 2000;Atanasoski et al., 2001).

Due to the slow rate of axonal growth, nerve repair often requires an extended period. Under long-term denervation, Schwann cells are unable to maintain the repair phenotype, accompanied by downregulation of repair-associated factors and even apoptosis. Accordingly, an apoptosis module was incorporated into the network (Scheib and Höke, 2013;Jessen and Mirsky, 2019).

Finally, during the transmission of dedifferentiation signals in Schwann cells, several key signaling pathways play critical roles, including the JNK, ERK, and Akt pathways. These pathways are involved in multiple processes of Schwann cell dedifferentiation and were therefore integrated into the network as essential signaling components (Nocera and Jacob, 2020;Wei et al., 2024). The selection criteria for core factors are described below.

The regulatory network constructed in this study does not attempt to encompass all molecular components at the genome-wide level. This reflects the current state of biological knowledge, as many regulatory factors involved in Schwann cell repair remain incompletely characterized, and even among known factors, most molecular interactions and intracellular kinetic parameters are not fully defined (Schellenberger and Palsson, 2009;Kirk et al., 2015;Xu et al., 2016). For example, YAP/TAZ have been reported to contribute to Schwann cell myelination (Grove et al., 2017); however, this functional role is largely captured in our network by the core myelination-associated regulators Krox20 and Sox10. Therefore, the effects of YAP/TAZ can be incorporated into these core factors. In addition, the role of YAP/TAZ in promoting Schwann cell proliferation after injury remains context-dependent and has been reported inconsistently across studies (Grove et al., 2020;Jeanette et al., 2021). Introducing YAP/TAZ as independent nodes may thus introduce redundancy and additional uncertainty, without substantially improving the functional representation of the myelination program. Similarly, TFEB/3 have been implicated in Schwann cell dedifferentiation and injury responses, but these functions are largely represented in the current model by key regulators such as c-Jun and Notch. Moreover, the currently available data on TFEB/3-mediated regulatory interactions remain relatively limited, making it difficult to define their precise roles within a mechanistic network framework. Collectively, the network was restricted to well-characterized core regulators with clearly established regulatory functions, ensuring both biological credibility and interpretability.

Biological systems are inherently organized in a hierarchical manner (Weinberg, 1995;Gehring, 1998;Noguchi et al., 2015). Consistently, single-gene knockout experiments have shown that most genes exert limited phenotypic effects when disrupted, supporting the existence of a core regulatory architecture (Hopkins, 2008). In this context, upstream master regulators such as c-Jun and Krox20 were selected to represent major functional programs. These factors were selected based on consistent experimental evidence supporting their central regulatory roles. Using Affymetrix whole-genome microarray analysis of injured mouse nerves, Arthur-Farraj et al. demonstrated that c-Jun is a key regulator of the Schwann cell injury response, controlling the expression of trophic factors and adhesion molecules, and contributing to regeneration track formation and myelin clearance (Arthur-Farraj et al., 2012b). Topilko et al. generated a Krox20 null allele (Krox20−/−) mouse model and demonstrated that loss of Krox20 arrests Schwann cells at an early differentiation stage, abolishes the expression of key myelin genes such as MBP and P0, and ultimately prevents peripheral nerve myelination (Topilko et al., 1994). In Runx2 knockout mouse models, researchers have observed impaired Schwann cell proliferation, migration, and axonal regeneration, indicating that Runx2 plays an important role in Schwann cell migration and peripheral nerve repair (Hu et al., 2024). As a master regulator of dedifferentiation, c-Jun can also activate downstream effectors to promote Schwann cell dedifferentiation, migration, and proliferation (Arthur-Farraj et al., 2012a;Fontana et al., 2012). Therefore, we integrated the functional role of Runx2 into that of c-Jun in our model.

In the present network, these core nodes are assumed to integrate the aggregate effects of their downstream effectors, effectively collapsing lower-level regulatory complexity into higher-level control modules. Explicit inclusion of numerous downstream factors would introduce redundancy, increase model complexity, and reduce both computational tractability and interpretability. Importantly, despite this abstraction, the simulation results remain in good agreement with existing experimental data and recapitulate key biological phenotypic transitions. Therefore, upstream core regulators were prioritized as key components of the network.

The regulatory factors included in the model also exhibit cell state specificity. Myelinating, non-myelinating (Remak), and repair Schwann cells are characterized by distinct molecular expression profiles. Accordingly, within the differentiation module, factors with minimal overlap across these cell states were preferentially selected, thereby enabling clear discrimination of cellular phenotypes based on their simulated expression signatures.

Finally, the selection of regulatory factors is problem-oriented. As this study focuses on Schwann cell dedifferentiation, priority was given to key regulators driving cellular reprogramming. In the following section, we describe the construction of the network in detail.

### Results of the construction of the endogenous regulatory network

The differentiation module is defined based on specific molecular markers associated with myelinating, non-myelinating, and reparative Schwann cells. In myelinating Schwann cells, the transcription factor Sox10 directly induces the key regulator Krox20, which is essential for peripheral myelin formation and thereby governs Schwann cell myelination (Reiprich et al., 2010;Jessen et al., 2015). Non-myelinating Schwann cells ensheath multiple small-caliber axons and express several molecular markers that are also present in immature Schwann cells, including NCAM and L1CAM (Jessen and Mirsky, 2002;Bolívar et al., 2020). After peripheral nerve injury, both myelinating and non-myelinating Schwann cells transdifferentiate into phenotypes that specifically promote nerve repair, providing metabolic support and guidance cues for neuronal survival, axonal regeneration, and target organ reinnervation. In the early stage, Schwann cells rapidly downregulate myelin-associated genes while activating repair-associated genes, such as c-Jun, Stat3, and Notch, to initiate dedifferentiation, and re-express cell adhesion molecules including NCAM and L1CAM. Subsequently, the expression of neurotrophic factors, such as brain-derived neurotrophic factor (BDNF), increases (Mirsky et al., 2008;Jessen and Mirsky, 2016). These molecules were collectively incorporated into the Schwann cell differentiation module.

During the transition toward a reparative phenotype, Schwann cells also participate in an inflammatory response, upregulating cytokines such as tumor necrosis factor-α (TNF-α) and interleukin-1β (IL-1β) (Jessen and Mirsky, 2016;Jessen and Arthur-Farraj, 2019). In the peripheral nerves of adult animals, Schwann cells remain in a quiescent state. Following injury, peripheral nerves undergo Wallerian degeneration, during which Schwann cells activate a proliferation program to facilitate nerve repair. At this stage, Schwann cell proliferation depends on cyclin D1. Notably, during embryonic development, D-type cyclins appear to be functionally redundant for Schwann cell growth. Therefore, we define the canonical cell cycle regulatory pathway, comprising Cyclin D/Cdk4,6, Rb, E2F, Myc, p53, and p21, as the proliferation module within the endogenous regulatory network of Schwann cells (Kim et al., 2000;Atanasoski et al., 2001). However, because of the slow pace of axonal growth during regeneration, the repair phenotype of Schwann cells is inherently unstable. Over prolonged periods of regeneration, Schwann cells undergo regression of the repair phenotype and may enter programmed cell death. Therefore, in the constructed endogenous network of Schwann cells, we incorporated the factors BAD, XIAP, CASP9, CASP3, Bcl2, and BAX to establish an apoptosis module (Scheib and Höke, 2013;Jessen and Mirsky, 2019;Wagstaff et al., 2021). Finally, following nerve injury, multiple signaling pathways in Schwann cells are activated in response to the insult, including the JNK, Ras/ERK, and PI3K/Akt pathways of the MAPK family (Jessen and Arthur-Farraj, 2019;Nocera and Jacob, 2020;Wei et al., 2024). The constructed endogenous regulatory network of Schwann cells comprises 30 nodes, 61 activating edges, and 43 inhibitory edges. In this network, nodes typically represent related genes, molecules, or signaling pathways. The network has no absolute upstream or downstream agents. To facilitate the representation of the network structure and enable subsequent dynamical modeling, the complex regulatory relationships were appropriately abstracted and simplified. Based on existing experimental evidence, the interactions between nodes were uniformly categorized into two types: activation and inhibition. Specifically, if one molecule promotes the production or activation of another, it is defined as an activation interaction. Conversely, if a molecule suppresses the expression of another or induces its inactivation, it is classified as an inhibitory interaction. Detailed regulatory relationships among all factors, along with the supporting references, are provided in Table S2 in Part A of the Supplementary Material. The interactions among these factors are shown in Figure 1.

**Figure 1.**
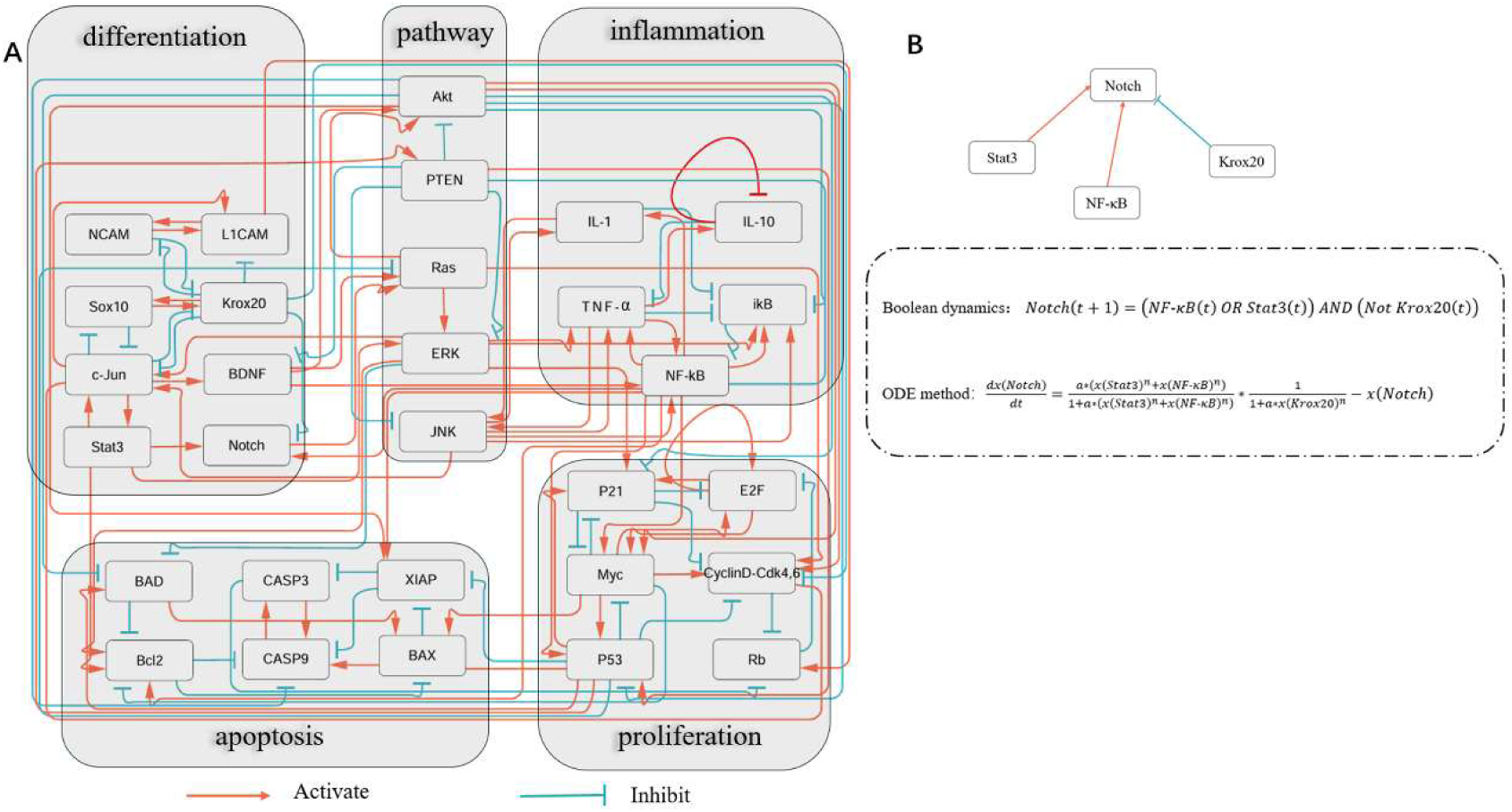
A) The core endogenous molecular-cellular network underlying Schwann cell dedifferentiation comprises 30 nodes, 61 activating edges, and 43 inhibitory edges. Specifically, orange edges denote activation, while blue edges represent inhibition. Nodes are further clustered into distinct groups based on modular classification. B) The endogenous network was modeled using both Boolean dynamics and differential equation–based approaches. As an illustrative example, the governing equation for the Notch node is presented. In this framework, x denotes the molecular concentration of the node, n is the Hill coefficient, and a represents the reciprocal of the apparent dissociation constant. The degradation rate is fixed at 1 in all models, corresponding to a linear decay term of −x. The complete gene interaction relationships are provided in Table S2 in Part A of the Supplementary Materials.

### Mathematical methods for quantifying endogenous networks

#### Boolean dynamics

Boolean dynamics provides a computational framework that can capture the structural information of a network without focusing too much on its details (Albert et al., 2008;Bornholdt, 2008a;Saadatpour and Albert, 2013). In this framework, Notch nodes can be represented as:

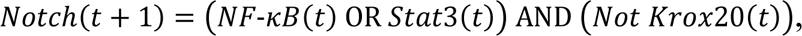

The equation forms of other factors are provided in Supplementary Section B. Boolean dynamics provides an effective method to capture the global structural characteristics of networks, as shown in Fig 1B. We adopted this model when constructing the endogenous network and compared it with the stochastic differential equation (SDE) method, which can identify saddle points that Boolean dynamics cannot recognize. It is worth noting that the modeling results of the core endogenous network are basically consistent between Boolean and differential equation representations, emphasizing that the dynamic behavior of the system is mainly determined by its intrinsic topology.

#### Stochastic differential equation

Compared with simplified Boolean dynamics and computationally demanding master equations, stochastic differential equations (SDEs) preserve model details while maintaining controllable computational complexity (Van Kampen, 1992;Bornholdt, 2008b). In endogenous networks, the expression levels or activities of n factors are represented as ***x*** = (x₁, x₂, …, xₙ)^T^. We employ SDEs to describe the temporal evolution of these factors, and the general form of the stochastic differential equation can be expressed as follows:

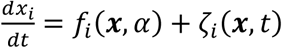

Among them, 𝑓_i_(𝒙, α) is a nonlinear function parameterized by α, and 𝜁 represents a multiplicative Gaussian white noise with a mean of zero. The correlation of the noise term is given by (ζ_i_(𝒙, 𝑡)ζ_j_(𝒙, 𝑡^’^)) = 2ɛ𝐷_ij_(𝒙)δ(𝑡 − 𝑡^’^). Here ⟨… ⟩ is the average distribution of noise, 𝜖 denotes the intensity of noise, 𝐷_ij_(𝒙) is the diffusion coefficient matrix, and 𝛿(𝑡 − 𝑡^’^) is the Dirac delta function. We have developed a framework for analyzing stochastic differential equations (Ao, 2004;2005;Yuan and Ao, 2012), in which the original equation can be decomposed into three components: a dissipative matrix 𝑆(𝒙), an antisymmetric matrix 𝐴(𝒙), and a potential function ф(𝒙, α),

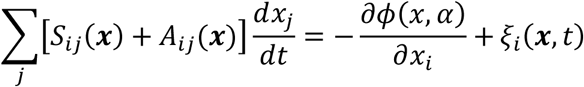

Where 𝜉 is the zero mean white noise with covariance satisfying (𝜉_i_(𝒙, 𝑡)𝜉_j_(𝒙, 𝑡^’^)) = 2𝜖𝑆_ij_(𝒙)𝛿(𝑡 − 𝑡^’^). By setting the stochastic and deterministic terms equal respectively, two relationships were obtained:

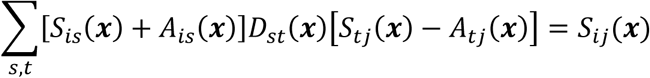

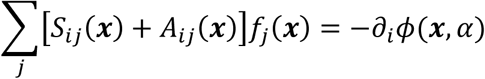

The potential function can be derived using ф(𝒙) = −∫ [𝑆(𝑥) + 𝐴(𝒙)]𝑓(𝒙)𝑑𝑥. This connects the potential function ф(𝒙, α) to the Boltzmann–Gibbs distribution 𝜌(𝒙) of the stochastic process in state space:

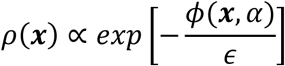

Because the system’s steady-state distribution follows the Boltzmann–Gibbs distribution, the potential function is decoupled from the noise intensity. Consequently, in our method, the potential function remains stable and invariant under any noise level. The existence of this potential function underpins our subsequent analysis.

In subsequent analyses, we adopt the method introduced above for evaluating endogenous networks, which effectively eliminates the dependence of Equation on noise. Focusing on the deterministic term of the equation 𝑓_i_(𝑥, α), we use the Hill function to describe how other factors exert activating or inhibitory influences on the expression level of the i-th agent(Ao et al., 2010):

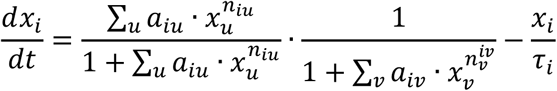

Here, n denotes the Hill coefficient, a is the reciprocal of the apparent dissociation constant, 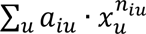 and 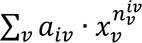 represent the cumulative activating and inhibitory influences on factor i, respectively, and τ_i_ denotes the degradation time of factor i, the equation for the Notch node, provided as an example, is shown in Figure 1B. Because precise estimation of these parameters across several orders of magnitude is challenging, we normalized the expression or activation levels of all factors to a range between 0 and 1.

Biological systems exhibit robust stability, and moderate changes in kinetic parameters generally do not substantially affect system behavior. In our endogenous network, the inherent topology of the network dictates the stability and adaptive behavior of the biological system (Barkai and Leibler, 1997;Kitano, 2002). In biochemical reactions, the Hill coefficient is typically in the range of 2–5 (Ahrends et al., 2014). Accordingly, we chose a value of 3 for all activation and inhibition interactions in our model. Parameter variations within reasonable bounds do not qualitatively alter the model predictions, reflecting the inherent robustness of the system (Yuan et al., 2017;Yao et al., 2024).In subsequent calculations, all τᵢ values were set to 1 for simplicity, assuming that factor degradation is primarily driven by cell division. Positioning the steep ‘S’ region of the sigmoid function near the center ensures that the network is most sensitive to input variations around the baseline, facilitates controllable steady states, and better captures threshold-like decision behaviors characteristic of biological systems. Accordingly, the parameters *a* and *n* satisfy the relationship 𝑎 ≈ 2^n^. Here, we set *a* = 3 and *n* = 10. Each factor in the network can be described as such differential equations, and all equations are provided in Part C of the supplementary materials.

### Solution to ODE

We employ the Newton and Euler methods to solve nonlinear differential equations, using uniformly distributed random vectors 𝑿_0_ as initial conditions:

The ODE system was simulated using an explicit Euler scheme with a fixed step size, and iterations continued until both the change between successive states and the magnitude of the vector field fell below predefined convergence thresholds, indicating convergence to a steady state. In parallel, fixed points of the Hill equations were directly identified using MATLAB’s *fsolve*, which applies the Newton method. Solutions of 𝑓(𝑋) = 0 were classified by evaluating the Jacobian matrix: points with all eigenvalues having negative real parts were designated as stable fixed points, whereas those with one or more positive eigenvalues were identified as saddle points, with the number of unstable directions recorded; solutions with multiple positive eigenvalues were further defined as hypertransition states. To ensure robustness and avoid numerical bias, we applied both the Euler method and the Newton method concurrently, and only steady states consistently detected by both approaches were retained.

### Perturbations on endogenous network

In general, disturbances to dynamical systems can be categorized into two major types: structural and positional perturbations. Structural perturbations manifest as modifications to the parameters or functional form of the governing equations, whereas positional perturbations arise from fluctuations in the system’s internal noise or changes in the external environment, such as drug-induced stimuli. Accumulated intrinsic noise can drive spontaneous transitions from one steady state to another via intermediate transition states, while external environmental shifts may directly displace the system from one basin of attraction to another.

In this study, we kept the governing equations unchanged and introduced small stochastic fluctuations to mimic noise-driven crossings of local energy barriers, enabling reconstruction of the transition pathways linking intermediate states to their adjacent steady states. This procedure allowed us to identify both the steady states and the corresponding transition trajectories shaped by the endogenous network structure. In addition, we implemented positional perturbations that emulate external interventions by modifying the initial expression levels of selected factors to mimic gene upregulation or downregulation, thereby simulating drug-induced environmental changes.

## Results

### Endogenous network modeling results of Schwann cells

Schwann cells undergo dedifferentiation to acquire a repair phenotype following peripheral nerve injury. To elucidate the underlying mechanisms, we constructed the core endogenous regulatory network of Schwann cells. The interactions among related modules and key signaling pathways are illustrated in Figure 1. As described previously, this network is closed, with each node both regulating and being regulated by others. There is no absolute upstream or downstream hierarchy within the system. The network operates autonomously and eventually converges to biologically meaningful steady states. The computational framework is detailed in the Materials and Methods section. The system was modeled using a set of high-dimensional nonlinear differential equations, we increased the number of random initial vectors in the nonlinear differential equation simulations from 10^4^ to 2 × 10^6^. The Euler method rapidly converged to 12 steady states from the outset. The Newton method initially identified 7 steady states and reached convergence to 12 steady states after 10^5^ iterations, as shown in Fig 2C. Similarly, the number of transition states stabilized at 47 after 10^5^ random initializations, as depicted in Figure 2B. These results indicate that the network’s steady states and transition states have been successfully identified. In the Boolean dynamic simulations, we tested one million distinct initial conditions and identified 37 attractors, comprising 7 point attractors and 30 linear attractors, as illustrated in Figure 2A. Comparison with the ODE results showed that each Boolean attractors had a corresponding ODE solution, with the linear attractors representing transition states, as shown in Figure 2D. As described in the Materials and Methods section, the agreement between the two modeling approaches confirms the reliability of our results. Detailed computational results for Boolean dynamics and differential equations are presented in Tables S3–S5 in Part D of the supplementary materials.

**Figure 2.**
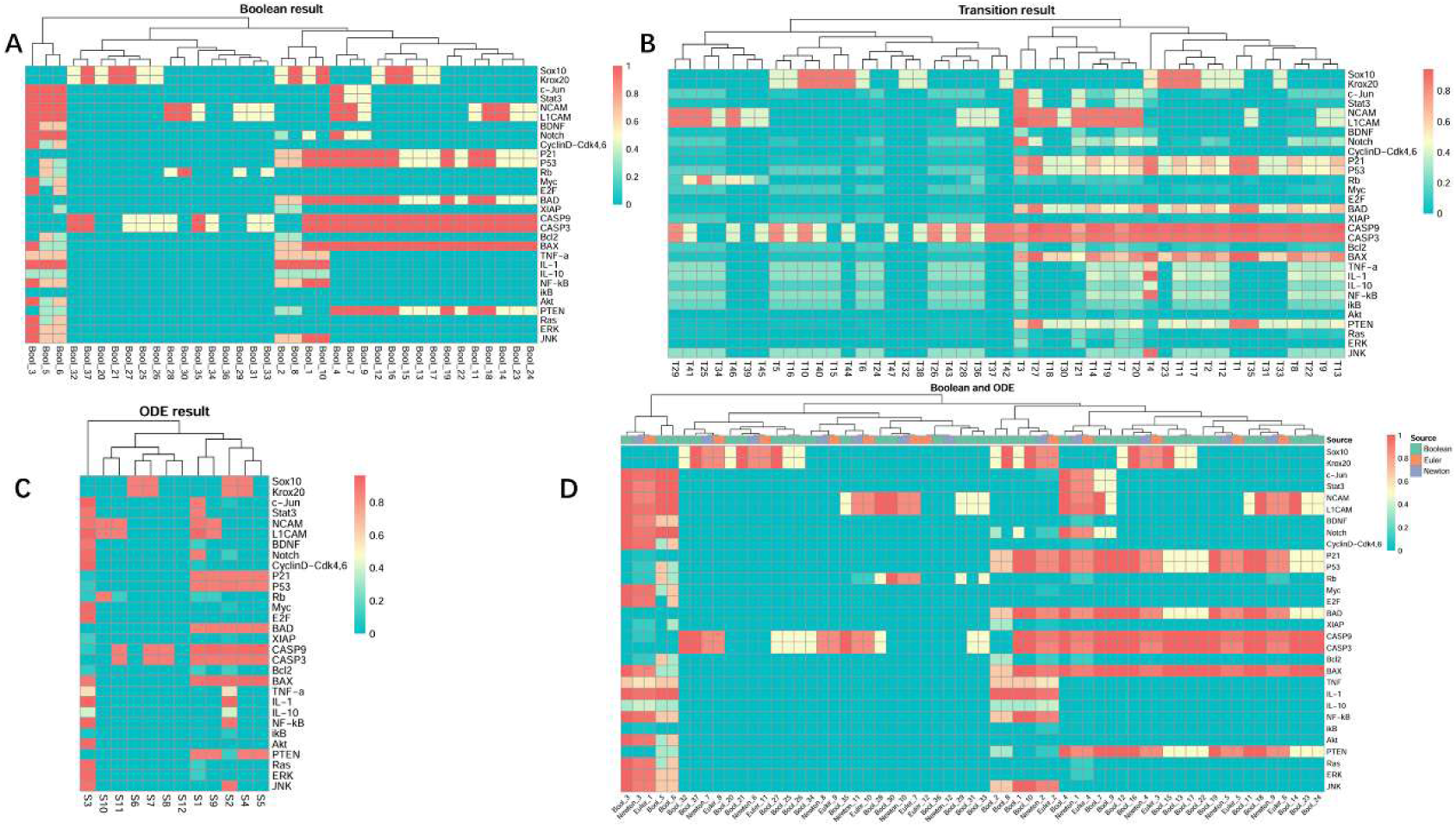
Endogenous networks of Schwann cells were quantified via differential equations and Boolean algebra. A) A total of 37 attractors were computed using Boolean algebra with one million random initial values. B) Forty-seven transition states were derived via Newton’s method following one million random vector iterations. C) Twelve steady states were solved using ordinary differential equations (ODEs). D) Comparative analysis of attractors from Boolean algebra versus steady states and transition states obtained via ODEs. The specific numerical values for each steady state are provided in the Part D of the Supplementary Materials.

### Steady states correspond to different cell types

To further investigate the potential biological significance of the steady states generated by the endogenous regulatory network, we first analyzed the biological implications of each steady state at the module level. Specifically, the status of each module was determined by examining the expression levels of factors within the corresponding functional modules in each steady state. In this study, a threshold of 0.5 was first set, with factor expression levels above 0.5 considered active (corresponding to 1) and those below 0.5 considered inactive (corresponding to 0). Accordingly, the expression levels of all factors were categorized into active and inactive states. Taking the cell cycle module as an example, we further illustrate how the functional state of a module under a steady state is determined based on the expression levels of its key factors. For each steady state, the status of each module was defined as either ON or OFF.

After thresholding, the expression levels of factors in the cell cycle module under steady state S3 are as follows:

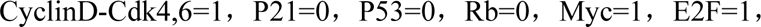

In steady state S3, the expression pattern of the cell cycle module exhibits a pro-proliferative signature, indicating its active state, as the activation of Cyclin D–Cdk4/6 promotes G1-to-S phase progression, Myc and E2F enhance cell cycle gene expression, and low levels of P21, P53, and Rb reduce inhibitory regulation. Similarly, the expression states of other modules were defined using the same approach. In particular, lineage-specific factors within the differentiation module were used to preliminarily define cell types, and the detailed expression values of all factors in each steady state are provided in Supplementary Material C. Based on the expression patterns of factors in the differentiation module, the 12 steady states were first classified into four categories, representing different Schwann cell types and functional states, as summarized in Table 1.Based on existing experimental evidence, the expression characteristics of myelinating, non-myelinating, and repair Schwann cell states across different functional modules are summarized in Table 2. Accordingly, we further determined which specific cell type each steady state represents.

**Table 1.**
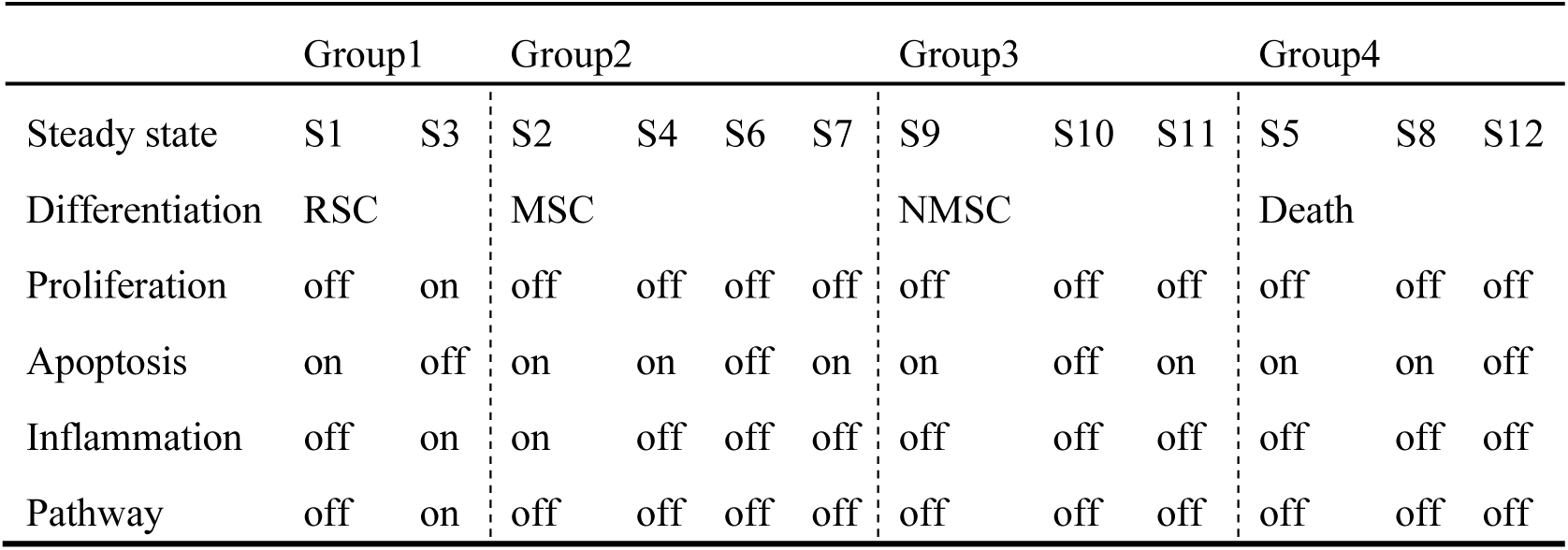
Preliminary categorization of steady states derived from the endogenous regulatory network of Schwann cells. Group1, RSC, repair Schwann cells; Group2, MSC, myelinating Schwann cells; Group3 NMSC, non-myelinating Schwann cells; Group4, Death, cells with apoptotic propensity or undergoing cell death.

**Table 2.** The left panel summarizes, based on experimental evidence, the distinct expression states of myelinating, non-myelinating, and repair Schwann cells across key functional modules. The right panel presents three representative steady states (S6, S10, and S3) derived from the endogenous network model, which are preliminarily classified as myelinating, non-myelinating, and repair Schwann cells, respectively, based on differentiation module markers.

| Experimental knowledge |  |  |  | Model steady state |  |  |
| --- | --- | --- | --- | --- | --- | --- |
| Cell type | MSC | NMSC | RSC | S6 | S10 | S3 |
| Proliferation | off | off | on | off | off | on |
| Apoptosis | off | off | off | off | off | off |
| Inflammation | off | off | on | off | off | on |
| Pathway | off | off | on | off | off | on |

In steady state S6, the myelin-associated transcription factors Krox20 and Sox10 are highly expressed. Within the proliferation module, CyclinD–Cdk4/6 expression is suppressed, indicating that cells are in a quiescent cell cycle state. In the apoptosis module, Casp3 and Casp9 are not expressed, suggesting that the apoptotic program is not activated. In addition, inflammatory factors are absent, indicating a lack of inflammatory response, and major signaling pathways remain inactive. In a study comparing the functions of mutant and wild-type Krox20, the latter was shown to promote Schwann cell exit from the cell cycle and suppress apoptosis (Arthur-Farraj et al., 2006). This is consistent with the characteristics of mature myelinating Schwann cells in intact nerves, and therefore we define S6 as representing the myelinating Schwann cell. Using the same analytical framework, we next examined other steady states. In steady state S10, NCAM and L1CAM are highly expressed, whereas myelin-associated and dedifferentiation-related factors are suppressed. The cell cycle remains in a quiescent state, with no expression of Casp3 and Casp9, indicating that apoptosis is not occurring. Inflammatory factors and signaling pathways are also suppressed. These features are consistent with the known characteristics of non-myelinating Schwann cells, particularly their association with NCAM and L1CAM expression (Yu et al., 2009;Jessen and Mirsky, 2019). Therefore, S10 is defined as the non-myelinating Schwann cell. In steady state S3,myelin-associated factors are suppressed, while dedifferentiation-related factors, including c-Jun, Stat3, and BDNF, are markedly upregulated. The increased expression of CyclinD–Cdk4/6 suggests that cells have re-entered the cell cycle and initiated proliferation. Meanwhile, Casp3 and Casp9 remain unexpressed, indicating that apoptosis is not activated. In addition, inflammatory factors such as TNF-α and IL-1 are activated, suggesting that the cells are undergoing an inflammatory response. Key signaling pathways, including JNK, Akt, and ERK, are also activated. These molecular features are consistent with experimentally reported characteristics of reparative Schwann cells, which exhibit downregulation of myelin-associated genes, upregulation of dedifferentiation-related factors, activation of inflammatory responses, and increased proliferative activity. Importantly, as no apoptotic program is engaged in this state, S3 corresponds to a normal reparative Schwann cell phenotype (Glenn and Talbot, 2013;Brosius Lutz and Barres, 2014;Jessen and Mirsky, 2016). In the following sections, steady state S6 will be referred to as the “M-steady state” (myelinating), S10 as the “NM-steady state” (non-myelinating), and S3 as the “RSC-steady state” (reparative Schwann cells).

Based on the previous analytical approach, we further examined the biological significance of other steady states. As shown in Figure 2C, in steady state S1, although factors associated with the reparative Schwann cell phenotype, such as c-Jun, Stat3, and Notch, are highly expressed, the neurotrophic factor BDNF is expressed at very low levels. Additionally, the cell cycle, inflammatory modules, and signaling pathways are suppressed, while the apoptosis module is activated, indicating the emergence of pro-apoptotic signaling. These features correspond to Schwann cells with reduced reparative capacity following prolonged denervation. This interpretation is supported by experimental evidence from a rat lumbar ventral root avulsion model, in which neurotrophic factors (e.g., BDNF, GDNF) exhibit a transient increase followed by a progressive decline over time in the absence of sustained regenerative support, correlating with diminished neuronal survival and limited axonal regeneration, indicating a gradual loss of the repair-supportive microenvironment. In addition, studies of chronic denervation show that Schwann cells undergo senescence characterized by cell cycle arrest and reduced proliferative capacity, further impairing regenerative support (Eggers et al., 2010;Ronchi et al., 2017;Fuentes-Flores et al., 2023). Steady state S2 is characterized by low expression of both neurotrophic and dedifferentiation-associated factors, accompanied by elevated inflammatory and pro-apoptotic signals and widespread pathway inactivation. Although this state exhibits a certain level of myelin-associated factor expression, insufficient reparative signaling and persistent inflammatory interference may prevent successful remyelination following injury, thereby leading to a progressive decline in reparative capacity. Previous studies have shown that chronic inflammation can disrupt Schwann cell repair functions and impair their ability to support nerve regeneration (Zhang et al., 2026). This profile is consistent with an inflammation-associated dysfunctional repair Schwann cell state observed under chronic denervation conditions (Fuentes-Flores et al., 2023). Therefore, this steady state is defined as a Schwann cell state with reduced reparative capacity induced by chronic inflammation. In the following text, steady states S1 and S2 are referred to as the “DRSC-steady state” (declining repair Schwann cell), reflecting their diminished reparative capacity. It should be noted that the term “DRSC-steady state” is used here as a functional grouping to indicate that both states are associated with impaired repair capacity, rather than implying complete equivalence between them.

Finally, in steady states S4, S5, S7, S8, S9, and S11, the apoptosis module is activated, accompanied by high expression levels of Casp3 and Casp9, while other modules and signaling pathways remain at low expression levels, indicating that the cells are predominantly executing the apoptotic program. In steady state S12, all factors are expressed at negligible levels. Therefore, these steady states were classified as apoptotic cells, and hereafter are collectively referred to as the “ASC-steady state” (apoptotic Schwann cell). This classification is based on shared apoptotic features, despite minor differences in molecular expression patterns among these steady states.

### Experimental verification

To further validate the biological relevance of the steady states derived from the mathematical model, we integrated multi-dimensional experimental evidence by systematically comparing the predicted expression patterns of key factors with both low-throughput experimental data and high-throughput transcriptomic datasets. Notably, validation primarily focused on reparative Schwann cells, based on two considerations. Firstly, reparative Schwann cells exhibit well-defined molecular characteristics in nerve injury models, with relatively abundant data available, ensuring the reliability of the validation benchmark. Second, mature myelinating and non-myelinating Schwann cells often exhibit overlapping transcriptional features and remain challenging to distinguish or purify in existing transcriptomic datasets, as reported in previous studies (Brosius Lutz et al., 2022). Therefore, the experimental validation was mainly conducted on reparative Schwann cells, evaluating the model predictions at the level of key molecular expression, thereby achieving a balance between data availability and analytical reliability.

We further validated the correspondence between the RSC-steady state (S3) and repair Schwann cells using low-throughput experimental data. Common low-throughput approaches, such as PCR, Western blotting, immunohistochemistry (IHC), and others, were used as the basis of the collected data(Parkinson et al., 2008). For experimental data, if a target factor is reported as significantly upregulated, strongly positive, or highly expressed in the literature or experimental observations, it is assigned as an active state with a value of 1. Conversely, if it is observed to be lowly expressed or undetectable, it is assigned as an inactive state with a value of 0. Factors lacking publicly available supporting data, as well as those not reported in a given study, were assigned a value of “na”. For the steady states derived from the model, since factor expression levels were normalized to the range of 0–1, a threshold of 0.4 was used for discretization. Specifically, values greater than 0.4 were assigned to 1, whereas values below 0.4 were assigned to 0, thereby establishing a basis for direct comparison between model predictions and experimental data. Except for several factors for which data were unavailable, the RSC-steady state (S3) representing repair Schwann cells showed a 92% consistency with the experimental results, supporting the reliability of the model prediction, as shown in Figure 3A. The references for the low-throughput data can be found in Part E of the Supplementary Material.

**Figure 3.**
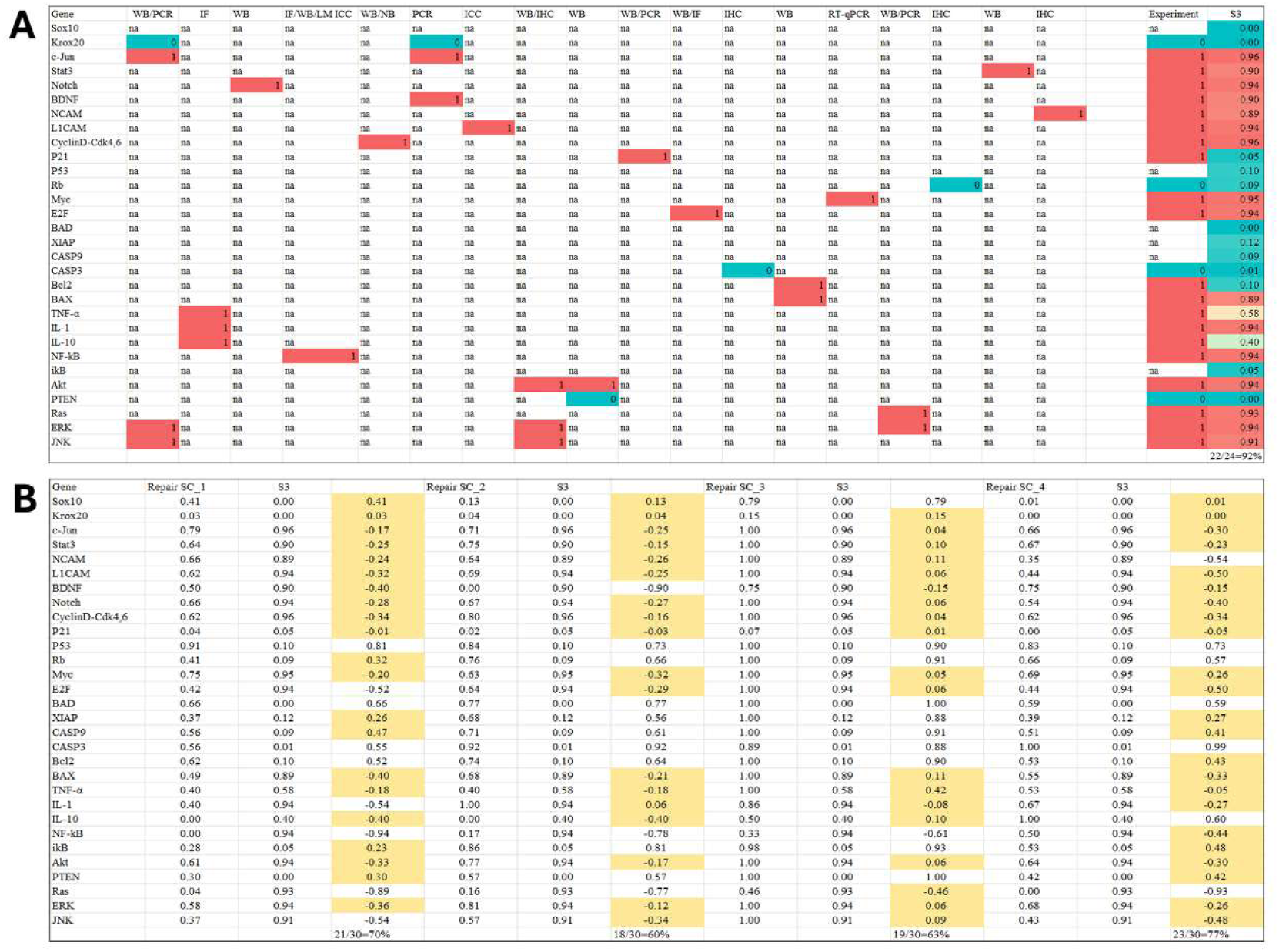
Experimental validation of the endogenous network computational results. Based on existing knowledge from low-throughput experimental data, the expression profiles of multiple genes in Schwann cells following differentiation were systematically collated. Each column of the table denotes one independent study or experimental dataset. B) Transcriptomic data of peripheral nerve injury models were retrieved from the public Gene Expression Omnibus (GEO) database (accession number: GSE109075). After preprocessing data from injured samples and control group samples, the transcriptomic data were normalized to the range of 0–1. Collected references for low-throughput experiments and normalization methods for high-throughput data are provided in the Part E of Supplementary Materials.

Although low-throughput experiments can accurately characterize the expression of specific factors, it is difficult for a single experiment to encompass all relevant molecules. Therefore, we further validated the reliability of the model predictions using high-throughput transcriptomic data. The dataset GSE109075 was selected, which profiles the transcriptomes of normal adult sciatic nerves and injured nerves. The data were normalized to a range of 0–1, where 0 represents the lowest expression level of a factor and 1 represents the highest. A threshold of 0.5 was set: if the absolute difference between the model-predicted value and the normalized experimental value was ≤ 0.5, the factor was considered consistent; otherwise, it was considered inconsistent. As illustrated in Figure 3B, the overall consistency between the calculated steady states and the high-throughput data ranged from 60% to 77%. Considering the inherent variability and experimental error associated with high-throughput measurements, this level of agreement is acceptable.

To validate the reliability of our model predictions, we further compared the computed steady states, namely the M-steady state, NM-steady state, and RSC-steady state, with single cell RNA sequencing (scRNA seq) data. We utilized a rat sciatic nerve injury scRNA seq dataset from GSE216665, which includes samples from uninjured (naive) nerves as well as 3, 12, and 60 days post injury. Data analysis was performed using the Seurat (v4) R package, and detailed procedures are provided in the Part F of the Supplementary Methods. As shown in Figure 4, the colored contour lines represent the density distribution of real single cells, while the black circles denote independently computed theoretical steady states. The temporal evolution exhibits a highly coherent and self-consistent trajectory. In the uninjured condition, the dark blue (myelinating) and yellow (Remak) cell populations are tightly clustered on the left side of the PC1 axis, closely enveloping and corresponding to the theoretically predicted M steady state and NM steady state, indicating that the model accurately captures the physiological states of healthy nerves. Following injury, a large number of red cells representing the repair phenotype emerge, and the centroid of their density contours shifts markedly toward the right along the PC1 axis. Notably, at day 12, the evolutionary trajectory of the cell population precisely converges toward the theoretically predicted RSC steady state. Within our computational framework, the RSC steady state represents an idealized steady state in a mathematical sense, corresponding to the theoretical limit of the repair phenotype. In contrast, cells in the day 12 post-injury samples remain in a dynamic transitional process toward this state. Although the cell population at this stage has not fully overlapped with the theoretical steady state in transcriptomic space, its trajectory shows a clear directional tendency consistent with attraction toward the RSC attractor. By day 60 post-injury, with the progression of chronic denervation, a subset of cells begins to undergo apoptosis or initiate remyelination, leading to a leftward shift in the distribution and a gradual return toward mature steady states. The strong agreement between the observed dynamic progression and the theoretically predicted trajectories provides compelling dynamical evidence for the model’s predictive power in determining cell fate transitions. It should be noted that, as our model does not rely on any previously published high-throughput datasets as input and the computed steady states correspond to idealized cellular states, this concordance provides robust support for the reliability of our model predictions.

**Figure 4.**
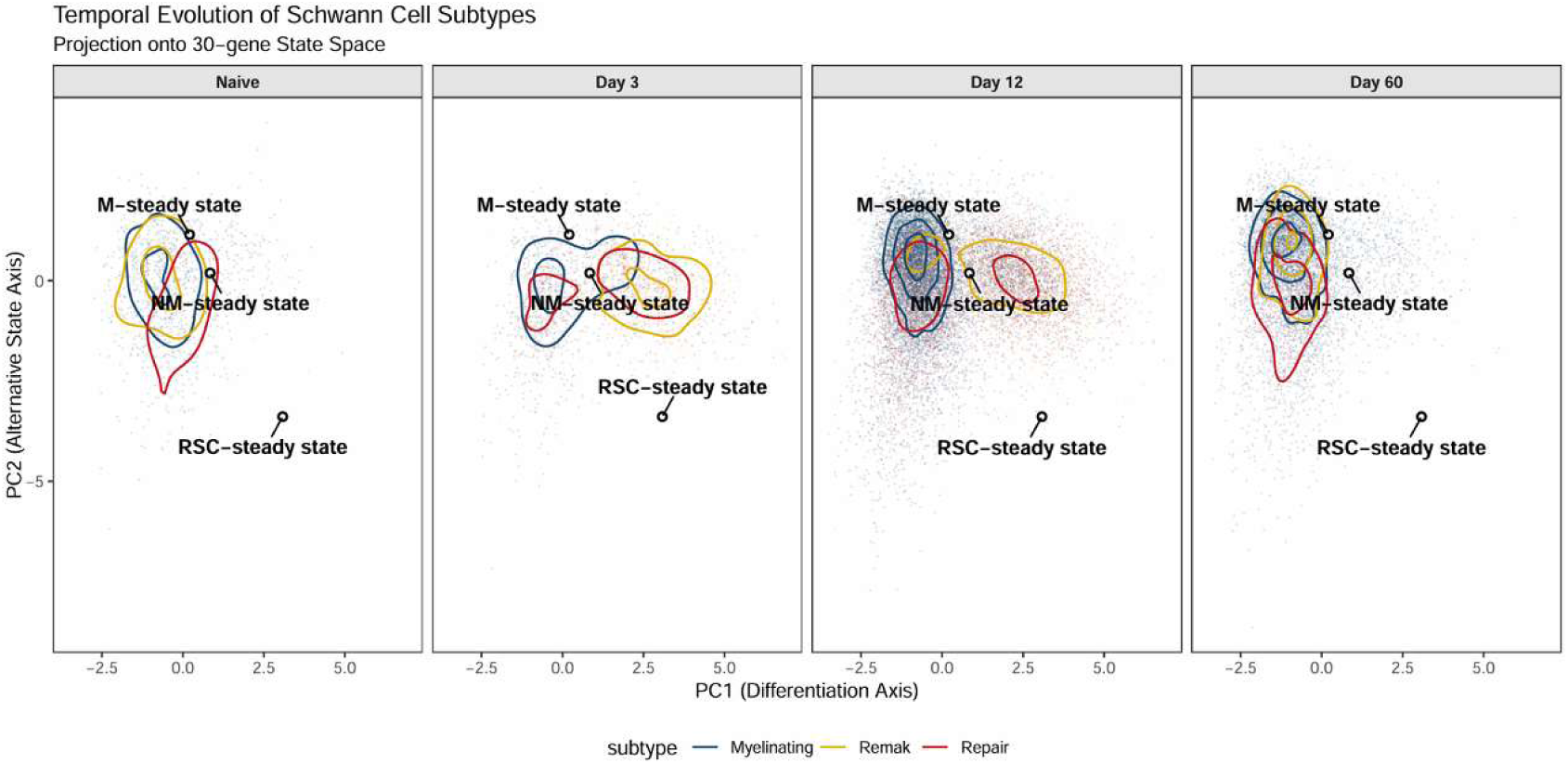
Temporal trajectory of Schwann cell phenotypic evolution projected onto the 30-gene regulatory state space. A-D, Principal Component Analysis (PCA) projection of single-cell transcriptomic profiles (GSE216665) from rat sciatic nerves at four stages: Naive (A), Day 3 (B), Day 12 (C), and Day 60 (D) post-injury. Each dot represents an individual cell, color-coded by its assigned subtype: Myelinating (blue), Remak (yellow), and Repair (red) Schwann cells. Density contours illustrate the spatial distribution and concentration of each subtype across the regenerative timeline. Theoretical Landmarks: The black open circles represent the three theoretical steady states derived independently from our mathematical regulatory model: Myelinating steady state (M-steady state), Non-myelinating steady state (NM-steady state), and Repair Schwann cell steady state (RSC-steady state). The detailed analytical procedures are provided in Part F of the Supplementary Information.

In summary, the consistencies observed between the model-predicted steady states and both low- and high-throughput experimental data fall within a reasonable range, further supporting the reliability of our computational results.

### Landscape of Schwann cell dedifferentiation

To characterize the dedifferentiation behavior of Schwann cells following peripheral nerve injury, we further introduced perturbations into the network to derive the transition paths between stable and transition states. By connecting all stable and transition states, we reconstructed the overall network topology, as shown in the Figure 5A.

**Figure 5.**
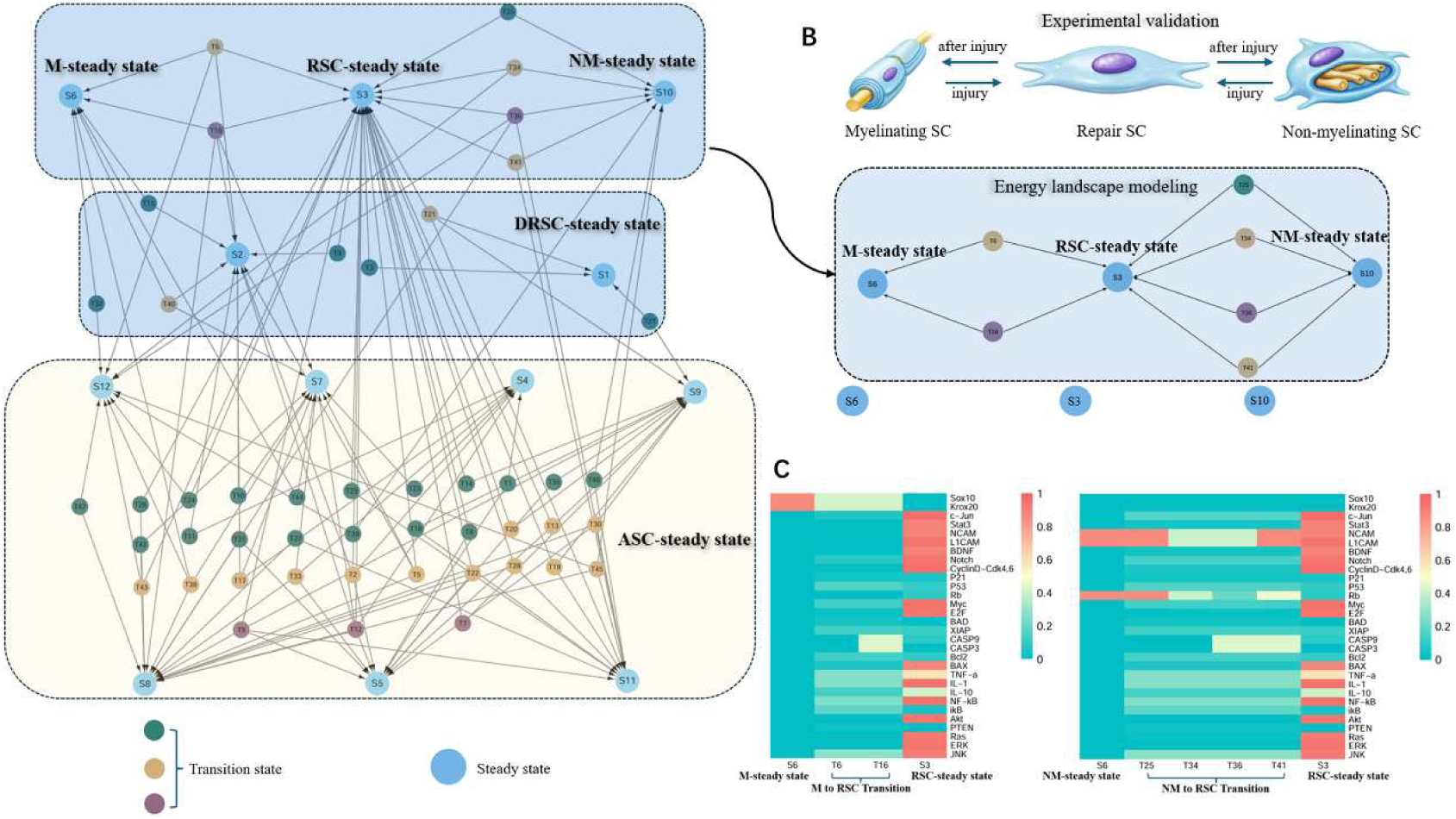
A) In the first blue box, steady state S6 corresponds to myelinating Schwann cells, S3 represents repair Schwann cells, and S10 denotes non-myelinating Schwann cells. These phenotypic states are interconnected through multiple transition states (T6, T16, T25, T34, T36, and T41), forming a structured dynamical landscape. The landscape derived from the endogenous network captures the dedifferentiation dynamics of Schwann cells following peripheral nerve injury. In the second blue box, steady states S2 and S1 represent Schwann cell states with reduced repair capacity. The steady state highlighted in the yellow box corresponds to Schwann cells exhibiting a propensity toward apoptosis. B) The upper panel summarizes experimentally validated dedifferentiation pathways of Schwann cells. The lower panel shows the cell state transition landscape obtained from endogenous network modeling (delineated by the blue box). The model-derived transitions are consistent with established experimental observations. C) Mechanisms underlying changes in factor expression profiles during the transition of myelinating and non- myelinating Schwann cells to reparative Schwann cells.

Transition states were classified according to the number of eigenvalues of the Jacobian matrix with non-negative real parts, these transition states interconnect distinct steady states. After overcoming the potential barriers between them, different steady states can achieve mutual conversion. The resulting landscape map reveals the dedifferentiation pathways of Schwann cells, as highlighted by the first blue box in Figure 5A. Within this framework, the M-steady state (S6), NM-steady state (S10), and RSC-steady state (S3) are interconnected by multiple transition states. In the computed landscape, both the M-steady state (S6) and NM-steady state (S10) can transition to the RSC-steady state (S3), Figure 5B. This finding is particularly noteworthy, as reported by Jessen, both myelinating and non-myelinating Schwann cells can dedifferentiate into repair Schwann cells following peripheral nerve injury. Our computational results successfully recapitulate this phenotypic transition (Jessen et al., 2015;Jessen and Mirsky, 2016;Zhang et al., 2023).

We further analyzed the transition states connecting the M-steady state (S6), NM-steady state (S10), RSC-steady state (S3). In the following sections, the transitional states connecting the M-steady state (S6) and the RSC-steady state (S3) are referred to as “M to RSC Transition”, while those connecting the NM-steady state (S10) and the RSC-steady state (S3) are referred to as “NM to RSC Transition”. Among them, the M to RSC Transition (T6, T16) directly link the M-steady state (S6) to the RSC-steady state (S3). As shown in Figure 5C, we visualized the expression profiles of the M to RSC Transition (T6, T16). Compared with the M-steady state (S6), both transitional states exhibit a prominent downregulation of myelin-associated factors, accompanied by an overall upregulation of inflammatory modules. In addition, the JNK signaling pathway becomes activated, while dedifferentiation-associated regulators, including c-Jun and Notch, show moderate increases. Proliferation-related factors are slightly upregulated but do not exhibit substantial changes overall, whereas pro-survival factors such as Bcl2 and XIAP display mild increases.

These molecular features are consistent with the well-established injury response program of Schwann cells described by Jessen and colleagues, in which myelin-associated genes are rapidly downregulated following nerve injury, accompanied by the re-expression of factors that are normally inactive in intact nerves, such as TNF-α, IL-1, and BDNF, as well as the induction of proliferative activity. Therefore, our results recapitulate key aspects of experimentally observed Schwann cell dedifferentiation (Jessen and Mirsky, 2016).

Previous studies have shown that following delayed nerve repair, the number of Schwann cells that can be isolated from the distal nerve stump gradually decreases (Saito et al., 2009;Jonsson et al., 2013). The observation of staged apoptotic events suggests that distinct Schwann cell subpopulations, characterized by different levels of stability and sensitivity to apoptosis, may undergo selective cell death at different time points. Notably, the apoptotic factors Casp3 and Casp9 exhibit higher expression levels in the M to RSC Transition (T16) compared with M to RSC Transition (T6). However, current studies on Schwann cell dedifferentiation have primarily focused on phenotypic reprogramming and regeneration-associated functions, with comparatively less emphasis on the apoptotic dynamics during intermediate transitional states. Based on this observation, we propose that repair Schwann cells derived from dedifferentiation may exhibit intrinsic heterogeneity. Specifically, transitional states such as M to RSC Transition (T16) may represent a subpopulation of Schwann cells that are more prone to apoptosis and therefore have a shorter lifespan compared with typical repair Schwann cells. The identification and targeted modulation of such vulnerable subpopulations may provide new opportunities for therapeutic intervention in peripheral nerve injury.

In non-myelinating Schwann cells, NM to RSC Transition (T25, T34, T36 and T41) connect to the RSC-steady state (S3). We further analyzed these transitional states based on their expression profiles in Figure 5C. It can be observed that the four transitional states share several common features. Specifically, they exhibit a moderate upregulation of c-Jun and Notch, a marked increase in inflammation-related factors, and a gradual elevation in the expression of cell cycle–associated genes. In addition, pro-survival factors such as Bcl2 and XIAP are upregulated, accompanied by progressive activation of the JNK signaling pathway. Notably, NCAM and L1CAM remain highly expressed across all four transitional states. In myelinating Schwann cells, the upregulation of NCAM and L1CAM during dedifferentiation is likely attributable to the fact that these adhesion molecules are suppressed under normal myelinating conditions and are reactivated upon injury. In contrast, non-myelinating Schwann cells intrinsically maintain the expression of these molecules, making their sustained high levels in transitional states biologically plausible. This observation is consistent with the characteristic molecular profile of non-myelinating Schwann cells, which express adhesion molecules such as NCAM and L1CAM while lacking myelin-associated proteins. Previous immunohistochemical analyses of human peripheral nerve sections have shown that non-myelinating Schwann cells express NCAM but do not express myelin-forming proteins, further supporting the biological plausibility of our results (Jessen and Mirsky, 2016;Jessen and Arthur-Farraj, 2019;Pestronk et al., 2023). Similar to the M to RSC transition (T16), elevated expression levels of casp9 and casp3 are also observed in the NM to RSC transitions (T36 and T41). This suggests that during the transition from non-myelinating Schwann cells to repair Schwann cells, intermediate states may also exhibit a certain degree of heterogeneity, with some transitional states potentially displaying a higher propensity for apoptosis.

In the previous section, we identified the steady states representing impaired repair capacity, namely the DRSC steady states (S1 and S2), and systematically characterized their expression profiles. Building upon this, we further investigated their relationship with the repair Schwann cell steady state (RSC steady state, S3).Within the endogenous regulatory network landscape constructed in this study, the transition states T4, T5, T12, and T16 connect the RSC steady state (S3) to the DRSC steady state (S2), describing a trajectory of progressive loss of repair capacity, see Figure 6A. For clarity, these transition states are hereafter collectively referred to as the R to DR Transition. The expression levels of all regulatory factors across these transition states are visualized in Figure 6B.From the expression patterns, dedifferentiation-associated factors are consistently downregulated during the R to DR Transition (T4, T5, T12, and T16), while myelination-related factors are progressively upregulated. Notably, apoptosis-related factors, including casp3 and casp9, show a gradual increase in expression. In parallel, the cell cycle is suppressed, key signaling pathways become progressively inhibited, and inflammatory factors such as TNF-α and IL-1 remain persistently expressed. Biologically, these results indicate that the cells are not following a remyelination trajectory, but instead exhibit sustained inflammatory activation. Ultimately, in the DRSC steady state (S2), casp3 and casp9 reach markedly elevated levels, indicating activation of the apoptotic program. This prediction is consistent with previous experimental observations and reviews indicating that chronic inflammation can lead to a decline in repair capacity and apoptosis of Schwann cells (Zhang et al., 2026). For example, a spatio-temporal analysis of motoneuron survival, axonal regeneration, and neurotrophic factor expression following ventral root avulsion and reimplantation has shown that, under conditions of chronic denervation, the expression of neurotrophic factors fails to be sustained over time (Eggers et al., 2010).This finding supports our model prediction that Schwann cells with impaired repair capacity are unable to maintain a pro-regenerative molecular program. Importantly, recent reviews have indicated that chronic inflammation can lead to downregulation of BDNF. In our computational results, the transition from the RSC-steady state (S3) to the DRSC-steady state (S2) is accompanied by sustained inflammatory activation and suppressed BDNF expression, indicating that our network successfully captures this process (Zhang et al., 2026).

**Figure 6.**
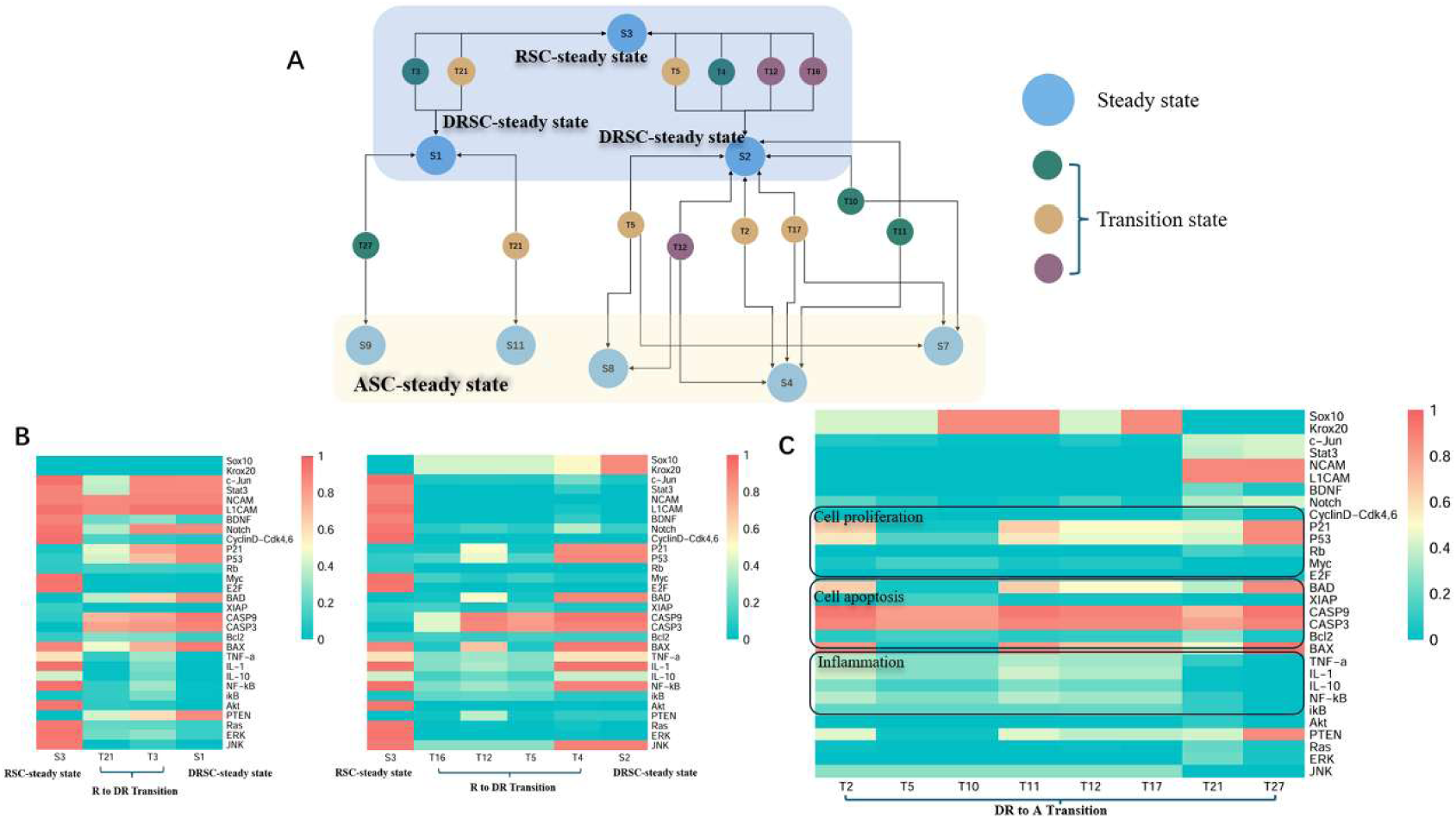
A) The endogenous network predicts the trajectory by which repair Schwann cells (RSC) undergo a decline in reparative phenotype and gradually exhibit an apoptotic tendency. The blue box encompasses repair Schwann cells and those with a dysfunctional phenotype (indicated by a downward arrow), whereas the yellow box denotes Schwann cells with an apoptotic propensity. B) The transition process from repair Schwann cells to a state of dysfunctional phenotype, along with the underlying mechanisms governing changes in factor expression. C) Factor expression profiles of transition states between dysfunctional Schwann cells and those with an apoptotic propensity.

In addition, as shown in Figure 6A, we analyzed the transition states T3 and T21 connecting the RSC-steady state (S3) and DRSC-steady state (S1), which we hereafter refer to as R to DR Transition (T3, T21). As shown in Figure 6B, these two transition states exhibit downregulation of BDNF, inhibition of the cell cycle, gradual upregulation of the apoptotic factors casp3 and casp9, very low expression of the anti-apoptotic factors Bcl2 and XIAP, and suppression of inflammation-related factors and signaling pathways, without signs of chronic inflammation. Notably, c-Jun is further downregulated in R to DR Transition (T21). Both transition states correspond to features of Schwann cells with declining repair capacity, resembling the long-term denervated Schwann cells. In a recent study using a mouse sciatic nerve transection model, SCs from aged animals and chronically denervated nerves were analyzed by immunofluorescence for markers of senescence. The analyses demonstrated that chronically denervated Schwann cells exhibit canonical features of repair capacity decline, such as irreversible cell cycle arrest. In a rat sciatic nerve transection model, the researchers observed a marked reduction in Schwann cell numbers in the distal stump after a 6-month delayed repair, indicating that prolonged denervation leads to a decline in Schwann cell repair capacity, potentially progressing to apoptosis (Boyd and Gordon, 2003;Höke and Brushart, 2010;Jonsson et al., 2013;Fuentes-Flores et al., 2023).

Moreover, the landscape generated from quantifying the endogenous network predicts the transition by which repairing Schwann cells progressively lose their phenotype and shift toward an apoptotic state. The degeneration of reparative Schwann cells is a multistage process that begins with a decline in the reparative phenotype, characterized by the progressive downregulation of repair-associated factors, and ultimately culminates in widespread cell death (Jessen and Mirsky, 2019). The modeling results derived from the endogenous network recapitulate this progression, as shown in Figure 6A. The transitional states connecting the DRSC-steady state and the ASC-steady state are hereafter referred to as the DR to A Transition. The expression profiles of these DR to A Transitions (T2, T5, T10, T11, T12, T17, T21, and T27) are visualized in Figure 6C. Overall, these transitions exhibit low expression of dedifferentiation-related factors, with the cell cycle module completely inactive, and generally low levels of inflammatory and signaling pathways. Importantly, the apoptotic factors caspase-3 (Casp3) and caspase-9 (Casp9) are markedly elevated, indicating activation of the apoptotic pathway. This suggests that the corresponding cellular states have begun to undergo programmed cell death, which explains why they are connected to the ASC-steady state. This systems-level analysis extends transcriptome-based observations by showing that loss of repair identity is closely linked to the activation of apoptosis. Together, these results suggest that the DR to A transition follows a structured, rather than stochastic, cell fate trajectory, providing a dynamical framework for understanding how repair-deficient Schwann cells progress toward apoptosis.

Notably, several transitional states (T5, T12, and T21) simultaneously connect the RSC-steady state with the DRSC-steady state, as well as the DRSC-steady state with the ASC-steady state. This topological feature suggests that these states occupy a central position within the regulatory landscape, acting as critical transition hubs rather than simple intermediate states along a linear trajectory. The dual connectivity of these states indicates that repair-deficient Schwann cells are not irreversibly committed to degeneration, but instead reside within a dynamically unstable region from which cells may either revert to a reparative phenotype or progress toward apoptosis. This further implies the existence of a potential decision window during which targeted interventions could redirect cell fate, highlighting these transitional states as key control points in the regulation of Schwann cell repair capacity. Together, these findings reveal that the transition from repair to degeneration is not a linear process, but is instead governed by a branched and dynamically regulated state-space architecture.

### Prediction of potential therapeutic targets

Based on the above analyses, we further investigated potential targets for restoring the repair capacity of Schwann cells undergoing functional decline. R to DR transition (T3,T21) exhibit reduced expression of inflammatory modules and attenuation of key signaling pathways. Consistently, R to DR transition (T4,T5,T12,T16) display decreased inflammatory activity, suppression of key pathways, downregulation of dedifferentiation-associated factors—including c-Jun, BDNF, and Stat3—and increased activation of apoptotic modules. Collectively, these findings suggest that enhancing inflammatory signaling, reactivating key pathways, upregulating core dedifferentiation factors, and inhibiting apoptosis may represent effective strategies for maintaining the reparative phenotype of Schwann cells. We next leverage the previously constructed network to test these predictions.

### Simulated target therapy

In endogenous networks, modulating the expression of specific factors can drive transitions between distinct cellular states. We applied this strategy to validate and simulate targeted interventions in Schwann cells. Our simulated interventions support existing findings while further extending mechanistic insights into Schwann cell (SC) repair capacity. We summarized the effective intervention strategies in Table 3. We first examined the role of c-Jun in Schwann cells. It is well established that c-Jun is a key regulator of Schwann cell dedifferentiation. During chronic denervation, Schwann cells exhibit a downregulation of c-Jun. Researchers established a chronic denervation model in mice by transecting the sciatic nerve. Ten weeks after injury, c-Jun expression was markedly downregulated, leading to impaired Schwann cell repair function and failed nerve regeneration. Maintenance of c-Jun expression was sufficient to rescue this deficit (Wagstaff et al., 2021). Similarly, another study established a chronic denervation model in rats and demonstrated that administration of Neurotrophin-3 maintained high levels of c-Jun expression, thereby promoting nerve regeneration (Xu et al., 2023). Importantly, loss-of-function experiments further confirmed the central role of c-Jun in Schwann cell-mediated nerve repair: mice with selective inactivation of c-Jun in Schwann cells exhibited severely impaired regenerative capacity, demonstrating that c-Jun acts as a global regulator of the Schwann cell injury response and is essential for the specification of the denervated repair Schwann cell state, including its characteristic gene expression program, cellular structure, and regenerative function (Arthur-Farraj et al., 2012a).

**Table 3.** For the two Schwann cell states with dysfunctional repair capacity (S1 and S2), we systematically upregulated or downregulated each regulatory factor within these steady states to assess whether they could be reprogrammed into an active, pro-repair phenotype. The resulting transitions are summarized in the table, which highlights the specific intervention strategies capable of driving these dysfunctional states toward a repair Schwann cell (RSC) identity.

| Steady state | Interference target | Interference measures | Result | Cell type |
| --- | --- | --- | --- | --- |
| S1<br>DRSC-steady state | BDNF | Downregulation | S1 | Dysfunctional RSC |
|  | BDNF | Upregulation | RSC-steady state | RSC |
|  | JNK | Downregulation | S1 | Dysfunctional RSC |
|  | JNK | Upregulation | RSC-steady state | RSC |
| | NF-κB | Downregulation | S1 | Dysfunctional RSC |
| | NF-κB | Upregulation | RSC-steady state | RSC |
|  | TNF | Downregulation | S1 | Dysfunctional RSC |
|  | TNF | Upregulation | RSC-steady state | RSC |
|  | P53 | Downregulation | RSC-steady state | RSC |
|  | P53 | Upregulation | S1 | Dysfunctional RSC |
|  | PTEN | Downregulation | RSC-steady state | RSC |
|  | PTEN | Upregulation | S1 | Dysfunctional RSC |
| S2<br>DRSC-steady state | c-Jun | Downregulation | S2 | Dysfunctional RSC |
|  | c-Jun | Upregulation | RSC-steady state | RSC |
|  | Krox20 | Downregulation | RSC-steady state | RSC |
|  | Krox20 | Upregulation | S2 | Dysfunctional RSC |
|  | Sox10 | Downregulation | RSC-steady state | RSC |
|  | Sox10 | Upregulation | S2 | Dysfunctional RSC |

As described above, the DRSC steady state (S2) represents repair-deficient Schwann cells, characterized by reduced c-Jun expression, whereas the RSC steady state (S3) corresponds to normal repair-competent Schwann cells. By simulating upregulation of c-Jun, we were able to drive the DRSC steady state (S2) back to the RSC steady state (S3), see Figure 7. This result indicates that our model recapitulates experimental observations showing that restoring or maintaining c-Jun expression can rescue the regenerative capacity of Schwann cells with impaired repair potential. Together with the genetic loss-of-function evidence, these findings further demonstrate the reliability of our model in capturing the causal role of c-Jun and in predicting effective molecular interventions (Jessen and Mirsky, 2021).

**Figure 7.**
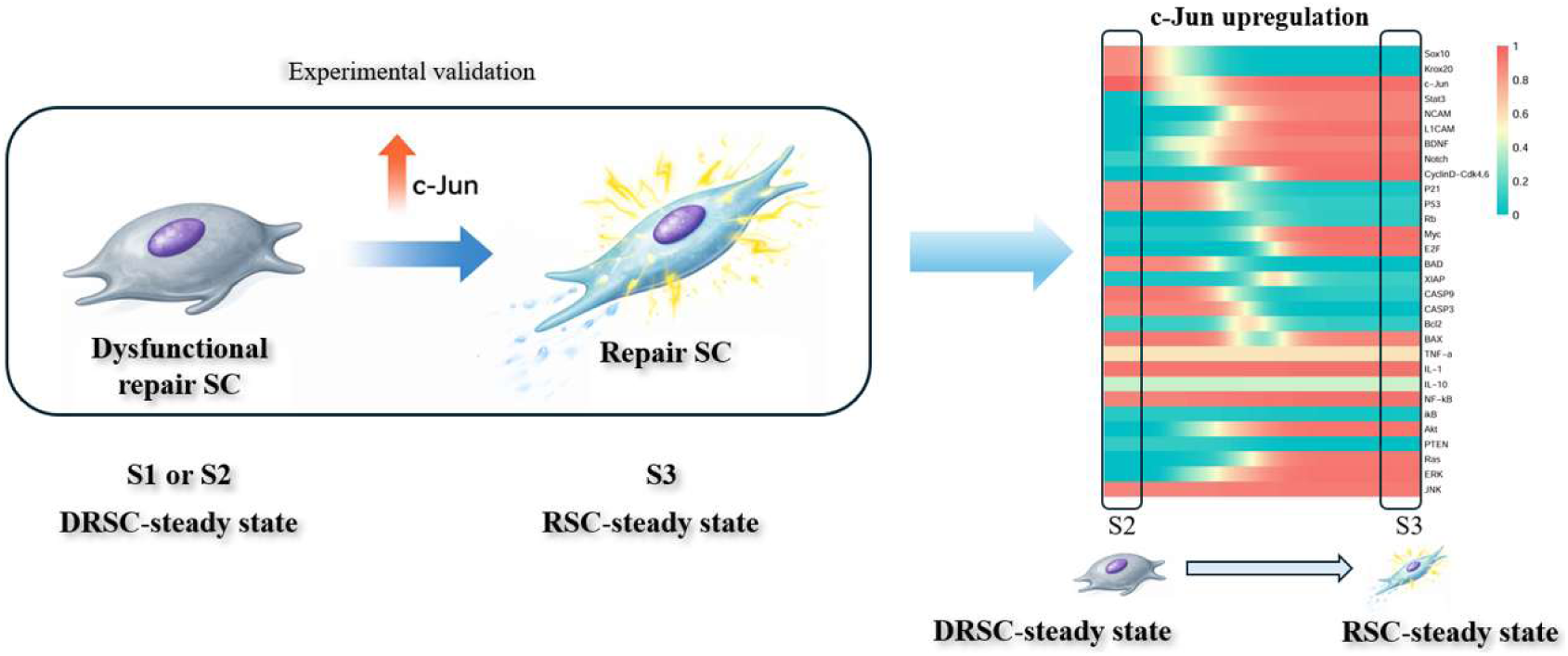
Target predicted by endogenous network modeling for maintaining the reparative capacity of Schwann cells. Upregulation of c-Jun (highlighted in the box) has been experimentally demonstrated to be essential for sustaining the reparative phenotype of Schwann cells (Wagstaff et al., 2021). In our model, DRSC-steady state (S2) correspond to dysfunctional repair Schwann cells, whereas RSC-steady state (S3) represents a repair Schwann cell state with robust repair ability. Consistent with experimental observations, enforced upregulation of c-Jun drives the transition from the DRSC-steady state (S2) to the RSC-steady state (S3), thereby restoring the reparative phenotype.

Researchers investigated the effects of BDNF on the JAK/STAT pathway using four different cell types. The results demonstrated that BDNF can activate the JAK/STAT pathway in both rat and human Schwann cells, and promote the release of the cytokine IL-6 (Lin et al., 2016). While the previous study did not specifically examine repair-deficient Schwann cells, these findings nonetheless suggest that BDNF upregulation may serve as a promising intervention strategy. In our simulated intervention experiments, upregulation of BDNF was able to reverse the decline in Schwann cell repair capacity, furthermore, IL-6 was also upregulated in the model (Fig. 8). This supplements existing knowledge by suggesting that BDNF is not only a physiological signaling molecule but also a viable therapeutic target capable of restoring impaired regenerative functions in severe or chronic injury contexts.

**Figure 8.**
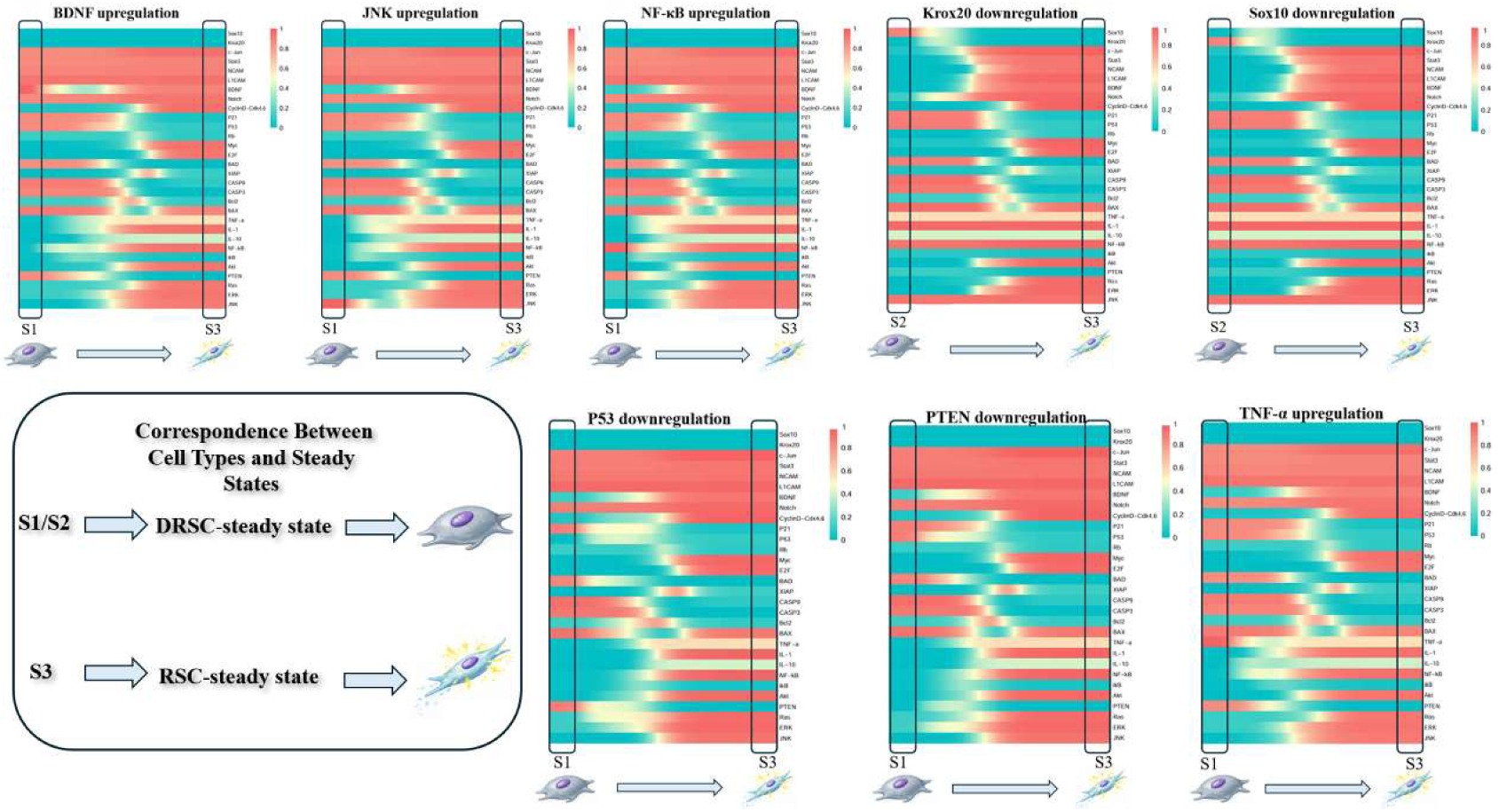
Effects of single-target perturbations on steady state transitions of repair-deficient Schwann cells. Heatmaps show the dynamic changes in gene expression profiles following individual target perturbations, including BDNF upregulation, JNK upregulation, NF-κB upregulation, TNF-α upregulation, as well as downregulation of Krox20, Sox10, P53, and PTEN. Each row represents a gene in the regulatory network, and the color scale (0–1) indicates normalized expression levels (teal: low; red: high). For each perturbation, the system trajectory is recorded from the initial steady state (DRSC-steady state) to the final steady state (RSC-steady state), with intermediate states shown along the horizontal axis. Black boxes highlight the regions corresponding to the initial and final steady states. Upregulation of BDNF, JNK, NF-κB, and TNF-α, as well as downregulation of P53 and PTEN, drives the system from the DRSC-steady state (S1) toward the RSC-steady state (S3). Downregulation of Krox20 and Sox10 induces the transition from the DRSC-steady state (S2) to the RSC-steady state (S3), indicating that appropriate interventions in repair-deficient Schwann cells can maintain their repair capacity. The schematic summary (bottom left) illustrates the correspondence between modeled steady states and biological cell types: S1/S2 correspond to the DRSC-steady state, representing repair-deficient Schwann cells, whereas S3 corresponds to the RSC-steady state, representing normal repair Schwann cells.

To our knowledge, there is no direct experimental evidence demonstrating that upregulation of JNK can reverse the decline in Schwann cell repair capacity. Regarding the signaling role of JNK, existing literature presents seemingly contradictory findings. Previous studies using cultured primary Schwann cells have shown that activation of the MKK7–JNK signaling pathway can induce c-Jun expression (Shin et al., 2013). However, another study showed that inhibition of JNK in transected nerve explants failed to affect injury-induced c-Jun upregulation, suggesting that c-Jun activation in Schwann cells may, at least in part, occur through JNK-independent mechanisms (Blom et al., 2014). Our simulations refine this understanding by demonstrating that while JNK may not be the sole or indispensable initiator of the injury response, its upregulation is sufficient to reverse the decline in SC repair capacity. In our model, upregulation of JNK drives the transition from the DRSC-steady (S1) state to the RSC-steady state (S3). This suggests that JNK may function as a context-dependent modulatory factor, contributing to the restoration, maintenance, and stability of the repair state, rather than acting as an essential upstream driver of c-Jun activation. By distinguishing between these two kinetic phases, the model supplements the existing knowledge, providing a plausible explanation for why JNK appears redundant in acute transection models yet remains a potent target for restoring long-term regenerative potential. This prediction suggests that JNK may contribute to restoring Schwann cell repair capacity when it is impaired, providing a testable hypothesis for future studies.

In addition, our simulated interventions indicate that upregulation of NF-κB and TNF-α is sufficient to maintain Schwann cell repair capacity. While prolonged or excessive inflammation is generally considered detrimental, controlled inflammatory signaling has been shown to support nerve repair (Zhang et al., 2026). For example, in a rat sciatic nerve injury model, TNF-α was shown to promote Schwann cell activation and macrophage recruitment, thereby facilitating myelin clearance and creating a permissive environment for regeneration (Iwatsuki et al., 2013). Similarly, in a zebrafish spinal cord injury model, pharmacological suppression of the inflammatory response impaired axonal regeneration, whereas enhancement of inflammation accelerated regrowth, demonstrating that an appropriate level of inflammation is required for effective repair (Tsarouchas et al., 2018).

Moreover, the predicted preservation of repair ability through the downregulation of P53 and PTEN confirms the roles of these molecules as critical molecular brakes during chronic denervation. We speculate that reducing P53 expression may decrease Schwann cells’ sensitivity to chronic denervation, thereby preventing apoptosis or entry into permanent senescence under such conditions, whereas PTEN downregulation likely alleviates inhibition of key pathways such as Akt, promoting Schwann cell survival. While previous studies focused on their roles in axonal regeneration (Ohtake et al., 2015), our model extends this understanding by suggesting that their suppression is critical for preventing the decline in Schwann cell repair capacity.

In our simulations, downregulation of Krox20 and Sox10 reversed the phenotype associated with reduced Schwann cell repair capacity. The antagonistic relationship between myelination and repair programs has been well documented (Topilko et al., 1994;Jessen and Arthur-Farraj, 2019). Our model builds on this by suggesting that relieving pro-myelinating transcriptional pressure alone may be sufficient to reactivate the repair program. This raises the possibility that the decline in repair capacity during chronic injury is partly driven by the persistent activity of myelination-associated inhibitory signals, providing a testable hypothesis for future studies.

In summary, our simulations indicate that modulating key regulatory nodes—including c-Jun, BDNF, JNK, NF-κB, TNF-α, P53, PTEN, Krox20, and Sox10—can restore or maintain Schwann cell repair capacity, and some of these interventions have been experimentally validated. These results confirm the predictive power of our model and provide a framework for identifying testable molecular targets to preserve or rescue Schwann cell regenerative potential. All intervention strategies are summarized in Figure 8.

## Discussion

In this study, we employed a bottom-up strategy to construct a core endogenous regulatory network of Schwann cells. By formulating dynamical equations, we derived the corresponding potential landscape and systematically characterized the global dynamics of Schwann cell fate regulation. Because no experimental data were used as direct inputs, the resulting landscape is not constrained by data quality or sample size, representing a fundamental distinction from data-driven approaches. This knowledge-based topological foundation successfully identified the major stable and transitional states of Schwann cells with high predictive validity against existing experimental datasets, allowing us to elucidate the intrinsic mechanisms of cellular plasticity. Beyond mechanistic insights, our model provides a powerful *in silico* platform to screen therapeutic targets for peripheral nerve injury. By simulating genetic perturbations, our model successfully recapitulated experimental observations—such as the rescue of declining repair capacity in chronic denervation via c-Jun upregulation (Wagstaff et al., 2021). Furthermore, we identified a broader spectrum of potent interventional targets, including BDNF, JNK, NF-κB, TNF-α, p53, PTEN, Krox20, and Sox10, which can similarly drive the transition of functionally declining Schwann cells back into a normal reparative state. In future studies, we will systematically explore combinations of these genetic perturbations to devise optimal therapeutic strategies.

Overall, our model provides new biological insights that go beyond the known transcriptional networks. First, we established a quantitative framework for Schwann cell fate regulation based on complex systems theory. This study overcomes the limitations of traditional approaches that focus on single molecules and linear signaling pathways by integrating complex systems theory with nerve repair research, thereby characterizing the dynamic process of Schwann cell fate transitions at the systems level. By introducing nonlinear dynamical modeling, we construct for the first time a multi-layered network—from functional modules to signaling pathways to key factors—that captures the interactions among regulatory elements and their collective behavior, providing a novel theoretical framework for understanding Schwann cell phenotypic transitions. Besides, our model improves the understanding of repair Schwann cell decline during chronic injury by revealing both the dynamic transition trajectories leading to repair failure and the underlying mechanisms, including the emergence of apoptosis-prone subpopulations and distinct pathways driving the progression toward apoptosis.

Importantly, for the first time, this study computationally derived the topological potential landscape of Schwann cell dedifferentiation, systematically recapitulating the dynamic trajectories from the normal state to the reparative state and further revealing multiple potential paths leading from declining repair capacity to apoptosis. These results not only overcome the limitations of experimental studies, which cannot continuously track cellular state transitions, but also provide a novel perspective for understanding the intrinsic dynamical mechanisms underlying Schwann cell fate differentiation.

Furthermore, we constructed a dynamical model that does not rely on high-throughput data yet achieves effective correspondence with experimental observations. Unlike studies that depend on large-scale omics data for correlation-based analyses, our model does not require external high-throughput data as input. Instead, it is built upon existing biological knowledge to establish the regulatory network, and steady states are obtained through nonlinear dynamical computations. The expression features of these predicted steady states correspond well with both low-throughput experimental results and transcriptomic data, demonstrating the potential of this approach for mechanistic interpretation and theoretical prediction even in the absence of large-scale datasets.

Finally, we predicted that dedifferentiated reparative Schwann cells may exhibit intrinsic heterogeneity based on differences in molecular expression across transitional states. In our simulations, the transitional state T16 connecting the myelinating and reparative steady states showed higher expression of apoptotic factors, potentially representing a Schwann cell subpopulation that is more prone to apoptosis; thus, compared with typical reparative Schwann cells, these cells are expected to have shorter lifespans. Experimental studies indicate that, following delayed nerve repair, the number of Schwann cells isolated from distal nerve stumps gradually decreases, suggesting that reparative Schwann cells are intrinsically heterogeneous: subpopulations with distinct stability and apoptosis sensitivity may undergo selective cell death at different stages, leading to the observed decline in cell numbers (Saito et al., 2009;Jonsson et al., 2013). However, to date, no studies have specifically targeted these apoptosis-prone subpopulations. Identification and targeted modulation of these apoptosis-prone subpopulations may provide new opportunities for therapeutic intervention in peripheral nerve injury.

In general, building on the currently established core endogenous network of Schwann cell dedifferentiation, we constructed a topological landscape to describe the dedifferentiation process. However, beyond capturing key processes such as inflammation, differentiation, proliferation, and apoptosis in adult Schwann cells following injury, we have not yet addressed fate determination during embryonic development and organismal aging, such as the differentiation of Schwann cell precursors and immature Schwann cells (Monk et al., 2015). Therefore, our computational results are currently limited to describing Schwann cell dedifferentiation after nerve injury. Schwann cell phenotypic plasticity is not only critical for nerve repair but also potentially contributes to regeneration in other tissues (Carr and Johnston, 2017). Future work could incorporate key modules and factors related to embryonic Schwann cell development and aging to expand the endogenous network, thereby linking developmental processes, post-injury cellular behaviors, and age-associated functional decline. This approach would enable the analysis of differences in their underlying dynamical mechanisms, providing new insights for precise modulation of Schwann cell fate and offering a theoretical foundation for regenerative medicine and clinical research.

## Statements

### Author Contributions

PA: Conceptualization, Methodology, Supervision, Writing-review & editing, Resources. ZY: Conceptualization, Investigation, Project administration, Validation, Visualization, Formal analysis, Software, Data curation, Writing-original draft. RX: Software, Methodology, Investigation, Resources, Writing-review & editing. SF: Investigation, Resources, Project administration, Writing-review & editing. YS: Project administration, Investigation, Writing-review & editing. QA: Supervision, Conceptualization, Writing-review & editing. Y-CC: Methodology, Conceptualization, Supervision, Writing-review & editing, Resources, Funding acquisition.

### Ethical Statement

This study did not involve human participants, animal experiments, or any procedures requiring ethical approval. All analyses were performed using mathematical modeling and publicly available datasets; therefore, no informed consent or ethical approval was required.

### Data Availability Statement

The datasets analyzed for this study can be found in the Gene Expression Omnibus (GEO) repository under the accession number GSE109075 and GSE216665.

## Supporting information

all supplemental materials

## References

Ahrends, R., Ota, A., Kovary, K.M., Kudo, T., Park, B.O., and Teruel, M.N. (2014). Controlling low rates of cell differentiation through noise and ultrahigh feedback. Science 344, 1384–1389.

Albert, I., Thakar, J., Li, S., Zhang, R., and Albert, R. (2008). Boolean network simulations for life scientists. Source Code Biol Med 3, 16.

Ao, P. (2004). Potential in stochastic differential equations: novel construction. Journal of physics A: mathematical and general 37, L25.

Ao, P. (2005). Laws in Darwinian evolutionary theory. Physics of life Reviews 2, 117–156.

Ao, P., Galas, D., Hood, L., Yin, L., and Zhu, X. (2010). Towards predictive stochastic dynamical modeling of cancer genesis and progression. Interdisciplinary Sciences: Computational Life Sciences 2, 140–144.

Ao, P., Galas, D., Hood, L., and Zhu, X. (2008). Cancer as robust intrinsic state of endogenous molecular-cellular network shaped by evolution. Med Hypotheses 70, 678–684.

Arthur-Farraj, P., Mirsky, R., Parkinson, D.B., and Jessen, K.R. (2006). A double point mutation in the DNA-binding region of Egr2 switches its function from inhibition to induction of proliferation: A potential contribution to the development of congenital hypomyelinating neuropathy. Neurobiology of disease 24, 159–169.

Arthur-Farraj, P.J., Latouche, M., Wilton, D.K., Quintes, S., Chabrol, E., Banerjee, A., Woodhoo, A., Jenkins, B., Rahman, M., Turmaine, M., Wicher, G.K., Mitter, R., Greensmith, L., Behrens, A., Raivich, G., Mirsky, R., and Jessen, K.R. (2012a). c-Jun reprograms Schwann cells of injured nerves to generate a repair cell essential for regeneration. Neuron 75, 633–647.

Arthur-Farraj, P.J., Latouche, M., Wilton, D.K., Quintes, S., Chabrol, E., Banerjee, A., Woodhoo, A., Jenkins, B., Rahman, M., Turmaine, M., Wicher, G.K., Mitter, R., Greensmith, L., Behrens, A., Raivich, G., Mirsky, R., and Jessen, K.R. (2012b). c-Jun Reprograms Schwann Cells of Injured Nerves to Generate a Repair Cell Essential for Regeneration. Neuron 75, 633–647.

Arthur-Farraj, P.J., Morgan, C.C., Adamowicz, M., Gomez-Sanchez, J.A., Fazal, S.V., Beucher, A., Razzaghi, B., Mirsky, R., Jessen, K.R., and Aitman, T.J. (2017). Changes in the Coding and Non-coding Transcriptome and DNA Methylome that Define the Schwann Cell Repair Phenotype after Nerve Injury. Cell Rep 20, 2719–2734.

Atanasoski, S., Shumas, S., Dickson, C., Scherer, S.S., and Suter, U. (2001). Differential cyclin D1 requirements of proliferating Schwann cells during development and after injury. Mol Cell Neurosci 18, 581–592.

Barkai, N., and Leibler, S. (1997). Robustness in simple biochemical networks. Nature 387, 913–917.

Benito, C., Davis, C.M., Gomez-Sanchez, J.A., Turmaine, M., Meijer, D., Poli, V., Mirsky, R., and Jessen, K.R. (2017). STAT3 Controls the Long-Term Survival and Phenotype of Repair Schwann Cells during Nerve Regeneration. J Neurosci 37, 4255–4269.

Blom, C.L., Mårtensson, L.B., and Dahlin, L.B. (2014). Nerve injury-induced c-Jun activation in Schwann cells is JNK independent. Biomed Res Int 2014, 392971.

Bolívar, S., Navarro, X., and Udina, E. (2020). Schwann Cell Role in Selectivity of Nerve Regeneration. Cells 9.

Bornholdt, S. (2008a). Boolean network models of cellular regulation: prospects and limitations. Journal of the Royal Society interface 5, S85–S94.

Bornholdt, S. (2008b). Boolean network models of cellular regulation: prospects and limitations. J R Soc Interface 5 Suppl 1, S85–94.

Boyd, J.G., and Gordon, T. (2003). Neurotrophic factors and their receptors in axonal regeneration and functional recovery after peripheral nerve injury. Mol Neurobiol 27, 277–324.

Brosius Lutz, A., and Barres, B.A. (2014). Contrasting the glial response to axon injury in the central and peripheral nervous systems. Dev Cell 28, 7–17.

Brosius Lutz, A., Lucas, T.A., Carson, G.A., Caneda, C., Zhou, L., Barres, B.A., Buckwalter, M.S., and Sloan, S.A. (2022). An RNA-sequencing transcriptome of the rodent Schwann cell response to peripheral nerve injury. J Neuroinflammation 19, 105.

Carr, M.J., and Johnston, A.P. (2017). Schwann cells as drivers of tissue repair and regeneration. Curr Opin Neurobiol 47, 52–57.

Davidson, E.H. (2010). The regulatory genome: gene regulatory networks in development and evolution. Elsevier.

Eggers, R., Tannemaat, M., Ehlert, E., and Verhaagen, J. (2010). A spatio-temporal analysis of motoneuron survival, axonal regeneration and neurotrophic factor expression after lumbar ventral root avulsion and implantation. Experimental neurology 223, 207–220.

Fontana, X., Hristova, M., Da Costa, C., Patodia, S., Thei, L., Makwana, M., Spencer-Dene, B., Latouche, M., Mirsky, R., Jessen, K.R., Klein, R., Raivich, G., and Behrens, A. (2012). c-Jun in Schwann cells promotes axonal regeneration and motoneuron survival via paracrine signaling. J Cell Biol 198, 127–141.

Fuentes-Flores, A., Geronimo-Olvera, C., Girardi, K., Necuñir-Ibarra, D., Patel, S.K., Bons, J., Wright, M.C., Geschwind, D., Hoke, A., Gomez-Sanchez, J.A., Schilling, B., Rebolledo, D.L., Campisi, J., and Court, F.A. (2023). Senescent Schwann cells induced by aging and chronic denervation impair axonal regeneration following peripheral nerve injury. EMBO Mol Med 15, e17907.

Gehring, W.J. (1998). Master control genes in development and evolution: The homeobox story. Yale University Press.

Glenn, T.D., and Talbot, W.S. (2013). Signals regulating myelination in peripheral nerves and the Schwann cell response to injury. Curr Opin Neurobiol 23, 1041–1048.

Grove, M., Kim, H., Santerre, M., Krupka, A.J., Han, S.B., Zhai, J., Cho, J.Y., Park, R., Harris, M., and Kim, S. (2017). YAP/TAZ initiate and maintain Schwann cell myelination. Elife 6, e20982.

Grove, M., Lee, H., Zhao, H., and Son, Y.-J. (2020). Axon-dependent expression of YAP/TAZ mediates Schwann cell remyelination but not proliferation after nerve injury. Elife 9, e50138.

Hartwell, L.H., Hopfield, J.J., Leibler, S., and Murray, A.W. (1999). From molecular to modular cell biology. Nature 402, C47–52.

Höke, A. (2006). Neuroprotection in the peripheral nervous system -: Rationale for more effective therapies. Archives of Neurology 63, 1681–1685.

Höke, A., and Brushart, T. (2010). Introduction to special issue: Challenges and opportunities for regeneration in the peripheral nervous system. Exp Neurol 223, 1–4.

Hopkins, A.L. (2008). Network pharmacology: the next paradigm in drug discovery. Nature chemical biology 4, 682–690.

Hu, R., Dun, X., Singh, L., and Banton, M.C. (2024). Runx2 regulates peripheral nerve regeneration to promote Schwann cell migration and re-myelination. Neural Regeneration Research 19, 1575–1583.

Iwatsuki, K., Arai, T., Ota, H., Kato, S., Natsume, T., Kurimoto, S., Yamamoto, M., and Hirata, H. (2013). Targeting anti-inflammatory treatment can ameliorate injury-induced neuropathic pain. PLoS One 8, e57721.

Jeanette, H., Marziali, L.N., Bhatia, U., Hellman, A., Herron, J., Kopec, A.M., Feltri, M.L., Poitelon, Y., and Belin, S. (2021). YAP and TAZ regulate Schwann cell proliferation and differentiation during peripheral nerve regeneration. Glia 69, 1061–1074.

Jessen, K.R., and Arthur-Farraj, P. (2019). Repair Schwann cell update: Adaptive reprogramming, EMT, and stemness in regenerating nerves. Glia 67, 421–437.

Jessen, K.R., and Mirsky, R. (2002). Signals that determine Schwann cell identity. J Anat 200, 367–376.

Jessen, K.R., and Mirsky, R. (2016). The repair Schwann cell and its function in regenerating nerves. Journal of Physiology-London 594, 3521–3531.

Jessen, K.R., and Mirsky, R. (2019). The Success and Failure of the Schwann Cell Response to Nerve Injury. Frontiers in Cellular Neuroscience 13, 14.

Jessen, K.R., and Mirsky, R. (2021). The Role of c-Jun and Autocrine Signaling Loops in the Control of Repair Schwann Cells and Regeneration. Front Cell Neurosci 15, 820216.

Jessen, K.R., Mirsky, R., and Lloyd, A.C. (2015). Schwann cells: development and role in nerve repair. Cold Spring Harb Perspect Biol 7, a020487.

Jonsson, S., Wiberg, R., Mcgrath, A.M., Novikov, L.N., Wiberg, M., Novikova, L.N., and Kingham, P.J. (2013). Effect of delayed peripheral nerve repair on nerve regeneration, Schwann cell function and target muscle recovery. PLoS One 8, e56484.

Kauffman, S.A. (1969). Metabolic stability and epigenesis in randomly constructed genetic nets. J Theor Biol 22, 437–467.

Kim, H.A., Pomeroy, S.L., Whoriskey, W., Pawlitzky, I., Benowitz, L.I., Sicinski, P., Stiles, C.D., and Roberts, T.M. (2000). A developmentally regulated switch directs regenerative growth of Schwann cells through cyclin D1. Neuron 26, 405–416.

Kirk, P.D., Babtie, A.C., and Stumpf, M.P. (2015). SYSTEMS BIOLOGY. Systems biology (un)certainties. Science 350, 386–388.

Kitano, H. (2002). Systems biology: a brief overview. Science 295, 1662–1664.

Kitaura, H., Shinshi, M., Uchikoshi, Y., Ono, T., Iguchi-Ariga, S.M., and Ariga, H. (2000). Reciprocal regulation via protein-protein interaction between c-Myc and p21(cip1/waf1/sdi1) in DNA replication and transcription. J Biol Chem 275, 10477–10483.

Lazebnik, Y. (2002). Can a biologist fix a radio?—Or, what I learned while studying apoptosis. Cancer cell 2, 179–182.

Lin, G., Zhang, H., Sun, F., Lu, Z., Reed-Maldonado, A., Lee, Y.C., Wang, G., Banie, L., and Lue, T.F. (2016). Brain-derived neurotrophic factor promotes nerve regeneration by activating the JAK/STAT pathway in Schwann cells. Transl Androl Urol 5, 167–175.

Maizels, R.J., and Briscoe, J. (2026). Gene regulatory networks: from correlative models to causal explanations. Nat Rev Genet.

Martini, R., Fischer, S., López-Vales, R., and David, S. (2008). Interactions between Schwann cells and macrophages in injury and inherited demyelinating disease. Glia 56, 1566–1577.

Mcmorrow, L.A., Kosalko, A., Robinson, D., Saiani, A., and Reid, A.J. (2022). Advancing Our Understanding of the Chronically Denervated Schwann Cell: A Potential Therapeutic Target? Biomolecules 12.

Mirsky, R., Woodhoo, A., Parkinson, D.B., Arthur-Farraj, P., Bhaskaran, A., and Jessen, K.R. (2008). Novel signals controlling embryonic Schwann cell development, myelination and dedifferentiation. J Peripher Nerv Syst 13, 122–135.

Monk, K.R., Feltri, M.L., and Taveggia, C. (2015). New insights on schwann cell development. Glia 63, 1376–1393.

Napoli, I., Noon, L.A., Ribeiro, S., Kerai, A.P., Parrinello, S., Rosenberg, L.H., Collins, M.J., Harrisingh, M.C., White, I.J., Woodhoo, A., and Lloyd, A.C. (2012). A Central Role for the ERK-Signaling Pathway in Controlling Schwann Cell Plasticity and Peripheral Nerve Regeneration In Vivo. Neuron 73, 729–742.

Nocera, G., and Jacob, C. (2020). Mechanisms of Schwann cell plasticity involved in peripheral nerve repair after injury. Cell Mol Life Sci 77, 3977–3989.

Noguchi, T.-a.K., Ninomiya, N., Sekine, M., Komazaki, S., Wang, P.-C., Asashima, M., and Kurisaki, A. (2015). Generation of stomach tissue from mouse embryonic stem cells. Nature cell biology 17, 984–993.

Ohtake, Y., Hayat, U., and Li, S. (2015). PTEN inhibition and axon regeneration and neural repair. Neural Regen Res 10, 1363–1368.

Parkinson, D.B., Bhaskaran, A., Arthur-Farraj, P., Noon, L.A., Woodhoo, A., Lloyd, A.C., Feltri, M.L., Wrabetz, L., Behrens, A., Mirsky, R., and Jessen, K.R. (2008). c-Jun is a negative regulator of myelination. J Cell Biol 181, 625–637.

Pestronk, A., Schmidt, R.E., Bucelli, R., and Sim, J. (2023). Schwann cells and myelin in human peripheral nerve: Major protein components vary with age, axon size and pathology. Neuropathol Appl Neurobiol 49, e12898.

Reiprich, S., Kriesch, J., Schreiner, S., and Wegner, M. (2010). Activation of Krox20 gene expression by Sox10 in myelinating Schwann cells. J Neurochem 112, 744–754.

Roberts, S.L., Dun, X.P., Doddrell, R.D.S., Mindos, T., Drake, L.K., Onaitis, M.W., Florio, F., Quattrini, A., Lloyd, A.C., D’antonio, M., and Parkinson, D.B. (2017). Sox2 expression in Schwann cells inhibits myelination in vivo and induces influx of macrophages to the nerve. Development 144, 3114–3125.

Ronchi, G., Cillino, M., Gambarotta, G., Fornasari, B.E., Raimondo, S., Pugliese, P., Tos, P., Cordova, A., Moschella, F., and Geuna, S. (2017). Irreversible changes occurring in long-term denervated Schwann cells affect delayed nerve repair. J Neurosurg 127, 843–856.

Saadatpour, A., and Albert, R. (2013). Boolean modeling of biological regulatory networks: a methodology tutorial. Methods 62, 3–12.

Saito, H., Kanje, M., and Dahlin, L.B. (2009). Delayed nerve repair increases number of caspase 3 stained Schwann cells. Neurosci Lett 456, 30–33.

Scheib, J., and Höke, A. (2013). Advances in peripheral nerve regeneration. Nature Reviews Neurology 9, 668–676.

Schellenberger, J., and Palsson, B. (2009). Use of randomized sampling for analysis of metabolic networks. J Biol Chem 284, 5457–5461.

Shin, Y.K., Jang, S.Y., Park, J.Y., Park, S.Y., Lee, H.J., Suh, D.J., and Park, H.T. (2013). The Neuregulin-Rac-MKK7 pathway regulates antagonistic c-jun/Krox20 expression in Schwann cell dedifferentiation. Glia 61, 892–904.

Sulaiman, O.a.R., and Gordon, T. (2009). Role of chronic Schwann cell denervation in poor functional recovery after nerve injuries and experimental strategies to combat it. Neurosurgery 65, A105–A114.

Sun, C., Yao, M., Xiong, R., Su, Y., Zhu, B., Chen, Y.C., and Ao, P. (2024). Evolution of Telencephalon Anterior-Posterior Patterning through Core Endogenous Network Bifurcation. Entropy (Basel) 26.

Svaren, J., and Meijer, D. (2008). The molecular machinery of myelin gene transcription in Schwann cells. Glia 56, 1541–1551.

Topilko, P., Schneider-Maunoury, S., Levi, G., Baron-Van Evercooren, A., Chennoufi, A.B., Seitanidou, T., Babinet, C., and Charnay, P. (1994). Krox-20 controls myelination in the peripheral nervous system. Nature 371, 796–799.

Tsarouchas, T.M., Wehner, D., Cavone, L., Munir, T., Keatinge, M., Lambertus, M., Underhill, A., Barrett, T., Kassapis, E., Ogryzko, N., Feng, Y., Van Ham, T.J., Becker, T., and Becker, C.G. (2018). Dynamic control of proinflammatory cytokines Il-1β and Tnf-α by macrophages in zebrafish spinal cord regeneration. Nat Commun 9, 4670.

Van Kampen, N.G. (1992). Stochastic processes in physics and chemistry. Elsevier.

Waddington, C.H. (2014). The strategy of the genes. Routledge.

Wagstaff, L.J., Gomez-Sanchez, J.A., Fazal, S.V., Otto, G.W., Kilpatrick, A.M., Michael, K., Wong, L.Y.N., Ma, K.H., Turmaine, M., Svaren, J., Gordon, T., Arthur-Farraj, P., Velasco-Aviles, S., Cabedo, H., Benito, C., Mirsky, R., and Jessen, K.R. (2021). Failures of nerve regeneration caused by aging or chronic denervation are rescued by restoring Schwann cell c-Jun. Elife 10.

Wei, C., Guo, Y., Ci, Z., Li, M., Zhang, Y., and Zhou, Y. (2024). Advances of Schwann cells in peripheral nerve regeneration: From mechanism to cell therapy. Biomed Pharmacother 175, 116645.

Weinberg, R.A. (1995). The retinoblastoma protein and cell cycle control. cell 81, 323–330.

Weiss, T., Taschner-Mandl, S., Bileck, A., Slany, A., Kromp, F., Rifatbegovic, F., Frech, C., Windhager, R., Kitzinger, H., Tzou, C.H., Ambros, P.F., Gerner, C., and Ambros, I.M. (2016). Proteomics and transcriptomics of peripheral nerve tissue and cells unravel new aspects of the human Schwann cell repair phenotype. Glia 64, 2133–2153.

Wright, S. (1932). The roles of mutation, inbreeding, crossbreeding, and selection in evolution.

Xu, M.J., Chen, Y.C., Xu, J., Ao, P., and Zhu, X.M. (2016). Kinetic model of metabolic network for xiamenmycin biosynthetic optimisation. IET Syst Biol 10, 17–22.

Xu, X., Song, L., Li, Y., Guo, J., Huang, S., Du, S., Li, W., Cao, R., and Cui, S. (2023). Neurotrophin-3 promotes peripheral nerve regeneration by maintaining a repair state of Schwann cells after chronic denervation via the TrkC/ERK/c-Jun pathway. Journal of Translational Medicine 21, 733.

Yao, M., Su, Y., Xiong, R., Zhang, X., Zhu, X., Chen, Y.C., and Ao, P. (2024). Deciphering the topological landscape of glioma using a network theory framework. Sci Rep 14, 26724.

Yu, W.M., Yu, H., Chen, Z.L., and Strickland, S. (2009). Disruption of laminin in the peripheral nervous system impedes nonmyelinating Schwann cell development and impairs nociceptive sensory function. Glia 57, 850–859.

Yuan, R., and Ao, P. (2012). Beyond itô versus stratonovich. Journal of Statistical Mechanics: Theory and Experiment 2012, P07010.

Yuan, R., Zhu, X., Wang, G., Li, S., and Ao, P. (2017). Cancer as robust intrinsic state shaped by evolution: a key issues review. Rep Prog Phys 80, 042701.

Zhang, H., Zhang, Z., and Lin, H. (2023). Research progress on the reduced neural repair ability of aging Schwann cells. Front Cell Neurosci 17, 1228282.

Zhang, Y., Zhang, H., Su, Y., Nuo, M., Wu, W., Jiang, H., and Meng, X. (2026). The role of immune regulation in peripheral nerve regeneration: functions of inflammatory cells and cytokines. Front Pharmacol 17, 1735833.

Zhu, X.M., Yin, L., Hood, L., and Ao, P. (2004). Robustness, stability and efficiency of phage lambda genetic switch: dynamical structure analysis. J Bioinform Comput Biol 2, 785–817.

