## Supplementary material for "Endogenous network modeling reveals mechanisms of repair Schwann cell decline and potential recovery targets": all supplemental materials

**Supplemental Material**

**Abstract:** This Supplementary Information is organized into six parts. Part A details all regulatory relationships among the factors in the endogenous network of Schwann cells, together with the literature supporting these interactions. Part B lists all Boolean equations of the endogenous network. Part C describes the differential equations used to simulate the network dynamics. Part D presents all numerical solutions derived from differential equation–based calculations. Part E summarizes the literature used for low-throughput validation and describes the procedures for processing GEO transcriptomic datasets. Part F outlines the analytical procedures for single-cell data validation.

A． **The factors and interactions of the endogenous network**

Table S1 summarizes all modules of the network model and their constituent factors, comprising 30 nodes and 105 functional relationships. This comprehensive representation of the network facilitates a deeper understanding of Schwann cell dedifferentiation.

**Table S1.** Modules and factors of endogenous network in Schwann cells

| **Modules** | **Factors** |
| --- | --- |
| Differentiation | Krox20; Sox10; NCAM; L1CAM; c-Jun; BDNF; Stat3; Notch |
| Proliferation | P21; E2F; Myc; P53; Rb; CyclinD-Cdk4,6 |
| Inflammation | IL-1; IL-10; TNF-α; iκB; NF-κB |
| Apoptosis | BAD; Bcl2; XIAP; BAX; Caspase3; Caspase9 |
| Pathway | JNK; Akt; Ras/ERK; PTEN |

The core endogenous network of Schwann cells was constructed based on experimental evidence, with regulatory relationships among factors curated from the literature. The following section details the interactions among all factors included in the endogenous network, together with the supporting references.

**Table S2.** Regulatory relationships among factors in the network

|  | **Factors** | **Activated by** | **Inhibit by** | **Reference** |
| --- | --- | --- | --- | --- |
| x(1) | Sox10 | Krox20 | c-Jun | [53; 58] |
| x(2) | Krox20 | Sox10 | c-Jun;NCAM | [58] |
| x(3) | c-Jun | Stat3;ERK;JNK | Krox20;Sox10 | [82; 13; 52; 44; 79] |
| x(4) | Stat3 | c-Jun |  | [82] |
| x(5) | NCAM | L1CAM | Krox20 | [33] |
| x(6) | L1CAM | NCAM;c-Jun | Krox20 | [71] |
| x(7) | BDNF | c-Jun | PTEN | [68; 71] |
| x(8) | Notch | NF-κB;Stat3 | Krox20 | [80; 78; 17] |
| x(9) | CyclinD-Cdk4,6 | Akt;Myc;Ras;c-Jun | P21;Krox20;P53 | [48; 26; 49; 2; 52; 50] |
| x(10) | P21 | E2F;ERK;P53 | Akt;Myc | [29; 66; 64; 56; 25; 1] |
| x(11) | P53 | Myc;PTEN;NF-κB | Akt | [18; 23; 39; 11; 51; 69] |
| x(12) | Rb | L1CAM | CyclinD-Cdk4,6;Caspase3 | [76; 38; 35] |
| x(13) | Myc | NF-κB;E2F;Akt | P21;P53 | [11; 51; 41; 84; 1] |
| x(14) | E2F | E2F;Myc | Rb;P21 | [48; 15; 66; 28] |
| x(15) | BAD | P53 | Akt;ERK | [12; 65; 34] |
| x(16) | XIAP | NF-kB;Akt | BAX;P53 | [51; 32; 37; 8; 10] |
| x(17) | Caspase9 | BAX;Caspase3 | Akt;Bcl2;XIAP | [7; 24; 67; 46; 47] |
| x(18) | Caspase3 | Caspase9 | XIAP | [14; 24; 62] |
| x(19) | Bcl2 | NF-κB;ERK;Stat3 | Myc;BAD | [5; 9; 32; 3; 30] |
| x(20) | BAX | BAD;P53;Myc | Bcl2 | [42; 73; 83; 30] |
| x(21) | TNF-α | NF-κB;ERK;JNK | IL-10 | [51; 61; 57] |
| x(22) | IL-1 | NF-κB;JNK |  | [75; 43] |
| x(23) | IL-10 | TNF-α | IL-10 | [22; 54] |
| x(24) | NF-κB | JNK;BDNF;TNF-α | iκB | [20; 51; 81; 4] |
| x(25) | iκB | NF-κB;ERK;JNK | TNF-α;IL-1;Akt | [70; 51; 59; 81; 6; 45; 27; 77; 75] |
| x(26) | Akt | Ras;BDNF | PTEN | [74; 55; 40] |
| x(27) | PTEN | P53 | NF-κB | [69; 72] |
| x(28) | Ras | BDNF;Notch | P53 | [16; 21; 60] |
| x(29) | ERK | Stat3;Ras | PTEN | [74; 63; 55] |
| x(30) | JNK | IL-1;TNF-α | PTEN | [19; 31; 74; 36] |

**References for Supplementary Material**

[1]Abbas T, Dutta A (2009) p21 in cancer: intricate networks and multiple activities. Nat Rev Cancer 9:400-414.

[2]Bakiri L, Lallemand D, Bossy-Wetzel E, Yaniv M (2000) Cell cycle-dependent variations in c-Jun and JunB phosphorylation: a role in the control of cyclin D1 expression. Embo j 19:2056-2068.

[3]Bhattacharya S, Ray RM, Johnson LR (2005) STAT3-mediated transcription of Bcl-2, Mcl-1 and c-IAP2 prevents apoptosis in polyamine-depleted cells. Biochem J 392:335-344.

[4]Boyle K, Azari MF, Cheema SS, Petratos S (2005) TNFalpha mediates Schwann cell death by upregulating p75NTR expression without sustained activation of NFkappaB. Neurobiol Dis 20:412-427.

[5]Breitschopf K, Haendeler J, Malchow P, Zeiher AM, Dimmeler S (2000) Posttranslational modification of Bcl-2 facilitates its proteasome-dependent degradation: molecular characterization of the involved signaling pathway. Mol Cell Biol 20:1886-1896.

[6]Burow ME, Weldon CB, Melnik LI, Duong BN, Collins-Burow BM, Beckman BS, McLachlan JA (2000) PI3-K/AKT regulation of NF-kappaB signaling events in suppression of TNF-induced apoptosis. Biochem Biophys Res Commun 271:342-345.

[7]Cardone MH, Roy N, Stennicke HR, Salvesen GS, Franke TF, Stanbridge E, Frisch S, Reed JC (1998) Regulation of cell death protease caspase-9 by phosphorylation. Science 282:1318-1321.

[8]Carter BZ, Mak DH, Schober WD, Koller E, Pinilla C, Vassilev LT, Reed JC, Andreeff M (2010) Simultaneous activation of p53 and inhibition of XIAP enhance the activation of apoptosis signaling pathways in AML. Blood 115:306-314.

[9]Catz SD, Johnson JL (2001) Transcriptional regulation of bcl-2 by nuclear factor kappa B and its significance in prostate cancer. Oncogene 20:7342-7351.

[10]Dan HC, Sun M, Kaneko S, Feldman RI, Nicosia SV, Wang HG, Tsang BK, Cheng JQ (2016) Akt phosphorylation and stabilization of X-linked inhibitor of apoptosis protein (XIAP). J Biol Chem 291:22846.

[11]Dang CV (1999) c-Myc target genes involved in cell growth, apoptosis, and metabolism. Mol Cell Biol 19:1-11.

[12]Datta SR, Dudek H, Tao X, Masters S, Fu H, Gotoh Y, Greenberg ME (1997) Akt phosphorylation of BAD couples survival signals to the cell-intrinsic death machinery. Cell 91:231-241.

[13]Davis RJ (2000) Signal transduction by the JNK group of MAP kinases. Cell 103:239-252.

[14]Deveraux QL, Reed JC (1999) IAP family proteins--suppressors of apoptosis. Genes Dev 13:239-252.

[15]Dyson N (1998) The regulation of E2F by pRB-family proteins. Genes Dev 12:2245-2262.

[16]Eastman-Reks SB, Vedeckis WV (1986) Glucocorticoid inhibition of c-myc, c-myb, and c-Ki-ras expression in a mouse lymphoma cell line. Cancer Res 46:2457-2462.

[17]Espinosa L, Cathelin S, D'Altri T, Trimarchi T, Statnikov A, Guiu J, Rodilla V, Inglés-Esteve J, Nomdedeu J, Bellosillo B, Besses C, Abdel-Wahab O, Kucine N, Sun SC, Song G, Mullighan CC, Levine RL, Rajewsky K, Aifantis I, Bigas A (2010) The Notch/Hes1 pathway sustains NF-κB activation through CYLD repression in T cell leukemia. Cancer Cell 18:268-281.

[18]Evan GI, Wyllie AH, Gilbert CS, Littlewood TD, Land H, Brooks M, Waters CM, Penn LZ, Hancock DC (1992) Induction of apoptosis in fibroblasts by c-myc protein. Cell 69:119-128.

[19]Finch A, Holland P, Cooper J, Saklatvala J, Kracht M (1997) Selective activation of JNK/SAPK by interleukin-1 in rabbit liver is mediated by MKK7. FEBS Lett 418:144-148.

[20]Finco TS, Westwick JK, Norris JL, Beg AA, Der CJ, Baldwin AS, Jr. (1997) Oncogenic Ha-Ras-induced signaling activates NF-kappaB transcriptional activity, which is required for cellular transformation. J Biol Chem 272:24113-24116.

[21]Fitzgerald K, Harrington A, Leder P (2000) Ras pathway signals are required for notch-mediated oncogenesis. Oncogene 19:4191-4198.

[22]Foey AD, Parry SL, Williams LM, Feldmann M, Foxwell BM, Brennan FM (1998) Regulation of monocyte IL-10 synthesis by endogenous IL-1 and TNF-alpha: role of the p38 and p42/44 mitogen-activated protein kinases. J Immunol 160:920-928.

[23]Frisch SM, Francis H (1994) Disruption of epithelial cell-matrix interactions induces apoptosis. J Cell Biol 124:619-626.

[24]Fujita E, Egashira J, Urase K, Kuida K, Momoi T (2001) Caspase-9 processing by caspase-3 via a feedback amplification loop in vivo. Cell Death Differ 8:335-344.

[25]Gartel AL, Radhakrishnan SK (2005) Lost in transcription: p21 repression, mechanisms, and consequences. Cancer Res 65:3980-3985.

[26]Gille H, Downward J (1999) Multiple ras effector pathways contribute to G(1) cell cycle progression. J Biol Chem 274:22033-22040.

[27]Griffin BD, Moynagh PN (2006) In vivo binding of NF-kappaB to the IkappaBbeta promoter is insufficient for transcriptional activation. Biochem J 400:115-125.

[28]Harbour JW, Dean DC (2000) The Rb/E2F pathway: expanding roles and emerging paradigms. Genes Dev 14:2393-2409.

[29]Hiyama H, Iavarone A, Reeves SA (1998) Regulation of the cdk inhibitor p21 gene during cell cycle progression is under the control of the transcription factor E2F. Oncogene 16:1513-1523.

[30]Hoffman B, Liebermann DA (2008) Apoptotic signaling by c-MYC. Oncogene 27:6462-6472.

[31]Holtmann H, Enninga J, Kalble S, Thiefes A, Dorrie A, Broemer M, Winzen R, Wilhelm A, Ninomiya-Tsuji J, Matsumoto K, Resch K, Kracht M (2001) The MAPK kinase kinase TAK1 plays a central role in coupling the interleukin-1 receptor to both transcriptional and RNA-targeted mechanisms of gene regulation. J Biol Chem 276:3508-3516.

[32]Igney FH, Krammer PH (2002) Death and anti-death: tumour resistance to apoptosis. Nat Rev Cancer 2:277-288.

[33]Jessen KR, Mirsky R (2008) Negative regulation of myelination: relevance for development, injury, and demyelinating disease. Glia 56:1552-1565.

[34]Jiang P, Du W, Wu M (2007) p53 and Bad: remote strangers become close friends. Cell Res 17:283-285.

[35]Jo DH, Lee K, Kim JH, Jun HO, Kim Y, Cho YL, Yu YS, Min JK, Kim JH (2017) L1 increases adhesion-mediated proliferation and chemoresistance of retinoblastoma. Oncotarget 8:15441-15452.

[36]Kant S, Swat W, Zhang S, Zhang ZY, Neel BG, Flavell RA, Davis RJ (2011) TNF-stimulated MAP kinase activation mediated by a Rho family GTPase signaling pathway. Genes Dev 25:2069-2078.

[37]Karin M, Lin A (2002) NF-kappaB at the crossroads of life and death. Nat Immunol 3:221-227.

[38]Katsuda K, Kataoka M, Uno F, Murakami T, Kondo T, Roth JA, Tanaka N, Fujiwara T (2002) Activation of caspase-3 and cleavage of Rb are associated with p16-mediated apoptosis in human non-small cell lung cancer cells. Oncogene 21:2108-2113.

[39]Khwaja A, Rodriguez-Viciana P, Wennström S, Warne PH, Downward J (1997) Matrix adhesion and Ras transformation both activate a phosphoinositide 3-OH kinase and protein kinase B/Akt cellular survival pathway. Embo j 16:2783-2793.

[40]Kiss Bimbova K, Bacova M, Kisucka A, Galik J, Zavacky P, Lukacova N (2022) Activation of Three Major Signaling Pathways After Endurance Training and Spinal Cord Injury. Mol Neurobiol 59:950-967.

[41]Kitaura H, Shinshi M, Uchikoshi Y, Ono T, Iguchi-Ariga SM, Ariga H (2000) Reciprocal regulation via protein-protein interaction between c-Myc and p21(cip1/waf1/sdi1) in DNA replication and transcription. J Biol Chem 275:10477-10483.

[42]Levine AJ (1997) p53, the cellular gatekeeper for growth and division. Cell 88:323-331.

[43]Li S, Zhu X, Liu B, Wang G, Ao P (2015) Endogenous molecular network reveals two mechanisms of heterogeneity within gastric cancer. Oncotarget 6:13607-13627.

[44]Lopez-Bergami P, Huang C, Goydos JS, Yip D, Bar-Eli M, Herlyn M, Smalley KS, Mahale A, Eroshkin A, Aaronson S, Ronai Z (2007) Rewired ERK-JNK signaling pathways in melanoma. Cancer Cell 11:447-460.

[45]Madrid LV, Wang CY, Guttridge DC, Schottelius AJ, Baldwin AS, Jr., Mayo MW (2000) Akt suppresses apoptosis by stimulating the transactivation potential of the RelA/p65 subunit of NF-kappaB. Mol Cell Biol 20:1626-1638.

[46]Marsden VS, Ekert PG, Van Delft M, Vaux DL, Adams JM, Strasser A (2004) Bcl-2-regulated apoptosis and cytochrome c release can occur independently of both caspase-2 and caspase-9. J Cell Biol 165:775-780.

[47]McKenna S, García-Gutiérrez L, Matallanas D, Fey D (2021) BAX and SMAC regulate bistable properties of the apoptotic caspase system. Sci Rep 11:3272.

[48]Murray AW, Hunt T (1993) The cell cycle: an introduction: Oxford University Press New York.

[49]Obaya AJ, Mateyak MK, Sedivy JM (1999) Mysterious liaisons: the relationship between c-Myc and the cell cycle. Oncogene 18:2934-2941.

[50]Oh SJ, Cho H, Kim S, Noh KH, Song KH, Lee HJ, Woo SR, Kim S, Choi CH, Chung JY, Hewitt SM, Kim JH, Baek S, Lee KM, Yee C, Park HC, Kim TW (2018) Targeting Cyclin D-CDK4/6 Sensitizes Immune-Refractory Cancer by Blocking the SCP3-NANOG Axis. Cancer Res 78:2638-2653.

[51]Pahl HL (1999) Activators and target genes of Rel/NF-kappaB transcription factors. Oncogene 18:6853-6866.

[52]Parkinson DB, Bhaskaran A, Droggiti A, Dickinson S, D'Antonio M, Mirsky R, Jessen KR (2004) Krox-20 inhibits Jun-NH2-terminal kinase/c-Jun to control Schwann cell proliferation and death. J Cell Biol 164:385-394.

[53]Parkinson DB, Bhaskaran A, Arthur-Farraj P, Noon LA, Woodhoo A, Lloyd AC, Feltri ML, Wrabetz L, Behrens A, Mirsky R, Jessen KR (2008) c-Jun is a negative regulator of myelination. J Cell Biol 181:625-637.

[54]Paul WE (2012) Fundamental immunology: Lippincott Williams & Wilkins.

[55]Pylayeva-Gupta Y, Grabocka E, Bar-Sagi D (2011) RAS oncogenes: weaving a tumorigenic web. Nat Rev Cancer 11:761-774.

[56]Radhakrishnan SK, Feliciano CS, Najmabadi F, Haegebarth A, Kandel ES, Tyner AL, Gartel AL (2004) Constitutive expression of E2F-1 leads to p21-dependent cell cycle arrest in S phase of the cell cycle. Oncogene 23:4173-4176.

[57]Rajasingh J, Bord E, Luedemann C, Asai J, Hamada H, Thorne T, Qin G, Goukassian D, Zhu Y, Losordo DW, Kishore R (2006) IL-10-induced TNF-alpha mRNA destabilization is mediated via IL-10 suppression of p38 MAP kinase activation and inhibition of HuR expression. Faseb j 20:2112-2114.

[58]Reiprich S, Kriesch J, Schreiner S, Wegner M (2010) Activation of Krox20 gene expression by Sox10 in myelinating Schwann cells. J Neurochem 112:744-754.

[59]Romashkova JA, Makarov SS (1999) NF-kappaB is a target of AKT in anti-apoptotic PDGF signalling. Nature 401:86-90.

[60]Ruiz-León Y, Pascual A (2001) Brain-derived neurotrophic factor stimulates beta-amyloid gene promoter activity by a Ras-dependent/AP-1-independent mechanism in SH-SY5Y neuroblastoma cells. J Neurochem 79:278-285.

[61]Rutault K, Hazzalin CA, Mahadevan LC (2001) Combinations of ERK and p38 MAPK inhibitors ablate tumor necrosis factor-alpha (TNF-alpha ) mRNA induction. Evidence for selective destabilization of TNF-alpha transcripts. J Biol Chem 276:6666-6674.

[62]Salvesen GS, Duckett CS (2002) IAP proteins: blocking the road to death's door. Nat Rev Mol Cell Biol 3:401-410.

[63]Saraiva M, O'Garra A (2010) The regulation of IL-10 production by immune cells. Nat Rev Immunol 10:170-181.

[64]Seoane J, Le HV, Massagué J (2002) Myc suppression of the p21(Cip1) Cdk inhibitor influences the outcome of the p53 response to DNA damage. Nature 419:729-734.

[65]She QB, Solit DB, Ye Q, O'Reilly KE, Lobo J, Rosen N (2005) The BAD protein integrates survival signaling by EGFR/MAPK and PI3K/Akt kinase pathways in PTEN-deficient tumor cells. Cancer Cell 8:287-297.

[66]Sherr CJ, Roberts JM (1999) CDK inhibitors: positive and negative regulators of G1-phase progression. Genes Dev 13:1501-1512.

[67]Shiozaki EN, Chai J, Rigotti DJ, Riedl SJ, Li P, Srinivasula SM, Alnemri ES, Fairman R, Shi Y (2003) Mechanism of XIAP-mediated inhibition of caspase-9. Mol Cell 11:519-527.

[68]Song W, Volosin M, Cragnolini AB, Hempstead BL, Friedman WJ (2010) ProNGF induces PTEN via p75NTR to suppress Trk-mediated survival signaling in brain neurons. J Neurosci 30:15608-15615.

[69]Stambolic V, MacPherson D, Sas D, Lin Y, Snow B, Jang Y, Benchimol S, Mak TW (2001) Regulation of PTEN transcription by p53. Mol Cell 8:317-325.

[70]Sun S-C, Ganchi PA, Ballard DW, Greene WC (1993) NF-κB controls expression of inhibitor IκBα: evidence for an inducible autoregulatory pathway. Science 259:1912-1915.

[71]Tricaud N, Park HT (2017) Wallerian demyelination: chronicle of a cellular cataclysm. Cell Mol Life Sci 74:4049-4057.

[72]Vasudevan KM, Gurumurthy S, Rangnekar VM (2004) Suppression of PTEN expression by NF-kappa B prevents apoptosis. Mol Cell Biol 24:1007-1021.

[73]Vogelstein B, Lane D, Levine AJ (2000) Surfing the p53 network. Nature 408:307-310.

[74]Waite KA, Eng C (2002) Protean PTEN: form and function. Am J Hum Genet 70:829-844.

[75]Weber A, Wasiliew P, Kracht M (2010) Interleukin-1 (IL-1) pathway. Sci Signal 3:cm1.

[76]Weinberg RA (1995) The retinoblastoma protein and cell cycle control. cell 81:323-330.

[77]Windheim M, Stafford M, Peggie M, Cohen P (2008) Interleukin-1 (IL-1) induces the Lys63-linked polyubiquitination of IL-1 receptor-associated kinase 1 to facilitate NEMO binding and the activation of IkappaBalpha kinase. Mol Cell Biol 28:1783-1791.

[78]Woodhoo A, Alonso MB, Droggiti A, Turmaine M, D'Antonio M, Parkinson DB, Wilton DK, Al-Shawi R, Simons P, Shen J, Guillemot F, Radtke F, Meijer D, Feltri ML, Wrabetz L, Mirsky R, Jessen KR (2009) Notch controls embryonic Schwann cell differentiation, postnatal myelination and adult plasticity. Nat Neurosci 12:839-847.

[79]Xu X, Song L, Li Y, Guo J, Huang S, Du S, Li W, Cao R, Cui S (2023) Neurotrophin-3 promotes peripheral nerve regeneration by maintaining a repair state of Schwann cells after chronic denervation via the TrkC/ERK/c-Jun pathway. J Transl Med 21:733.

[80]Yoshimatsu T, Kawaguchi D, Oishi K, Takeda K, Akira S, Masuyama N, Gotoh Y (2006) Non-cell-autonomous action of STAT3 in maintenance of neural precursor cells in the mouse neocortex. Development 133:2553-2563.

[81]Zandi E, Karin M (1999) Bridging the gap: composition, regulation, and physiological function of the IkappaB kinase complex. Mol Cell Biol 19:4547-4551.

[82]Zhang X, Wrzeszczynska MH, Horvath CM, Darnell JE, Jr. (1999) Interacting regions in Stat3 and c-Jun that participate in cooperative transcriptional activation. Mol Cell Biol 19:7138-7146.

[83]Zhang Z, Lapolla SM, Annis MG, Truscott M, Roberts GJ, Miao Y, Shao Y, Tan C, Peng J, Johnson AE, Zhang XC, Andrews DW, Lin J (2004) Bcl-2 homodimerization involves two distinct binding surfaces, a topographic arrangement that provides an effective mechanism for Bcl-2 to capture activated Bax. J Biol Chem 279:43920-43928.

[84]Zhu J, Blenis J, Yuan J (2008) Activation of PI3K/Akt and MAPK pathways regulates Myc-mediated transcription by phosphorylating and promoting the degradation of Mad1. Proc Natl Acad Sci U S A 105:6584-6589.

B． **Boolean equations of the endogenous network in Table S2**

Sox10(t+1) = Krox20(t) AND (Not c-Jun(t))

Krox20(t+1) = Sox10(t) AND (Not (c-Jun(t) OR NCAM(t)))

c-Jun(t+1) = (Stat3(t) OR ERK(t) OR JNK(t)) AND (Not (Krox20(t) OR Sox10(t)))

Stat3(t+1) = c-Jun(t)

NCAM(t+1) = L1CAM(t) AND (Not Krox20(t))

L1CAM(t+1) = (c-Jun(t) OR NCAM(t)) AND (Not Krox20(t))

BDNF(t+1) = c-Jun(t) AND (Not PTEN(t))

Notch(t+1) = (NF-kB(t) OR Stat3(t)) AND (Not Krox20(t))

CyclinD-Cdk4,6(t+1) = (Akt(t) OR Myc(t) OR Ras(t) OR c-Jun(t)) AND (Not (P21(t) OR Krox20(t) OR P53(t)))

P21(t+1) = (E2F(t) OR ERK(t) OR P53(t)) AND (Not (Akt(t) OR Myc(t)))

P53(t+1) = (Myc(t) OR PTEN(t) OR NF-kB(t)) AND (Not Akt(t))

Rb(t+1) = L1CAM(t) AND (Not (CyclinD-Cdk4,6(t) OR CASP3(t)))

Myc(t+1) = (NF-kB(t) OR E2F(t) OR Akt(t)) AND (Not (P21(t) OR P53(t)))

E2F(t+1) = (E2F(t) OR Myc(t)) AND (Not (Rb(t) OR P21(t)))

BAD(t+1) = P53(t) AND (Not (Akt(t) OR ERK(t)))

XIAP(t+1) = (NF-kB(t) OR Akt(t)) AND (Not (BAX(t) OR P53(t)))

CASP9(t+1) = (BAX(t) OR CASP3(t)) AND (Not (Akt(t) OR Bcl2(t) OR XIAP(t)))

CASP3(t+1) = CASP9(t) AND (Not XIAP(t))

Bcl2(t+1) = (NF-kB(t) OR ERK(t) OR Stat3(t)) AND (Not (Myc(t) OR BAD(t)))

BAX(t+1) = (BAD(t) OR P53(t) OR Myc(t)) AND (Not Bcl2(t))

TNF-α(t+1) = (NF-kB(t) OR ERK(t) OR JNK(t)) AND (Not IL-10(t))

IL-1(t+1) = NF-kB(t) OR JNK(t)

IL-10(t+1) = TNF-α(t) AND (Not IL-10(t))

NF-kB(t+1) = (JNK(t) OR BDNF(t) OR TNF-α(t)) AND (Not ikB(t))

ikB(t+1) = (NF-kB(t) OR ERK(t) OR JNK(t)) AND (Not (TNF-α(t) OR IL-1(t) OR Akt(t)))

Akt(t+1) = (Ras(t) OR BDNF(t)) AND (Not PTEN(t))

PTEN(t+1) = P53(t) AND (Not NF-kB(t))

Ras(t+1) = (BDNF(t) OR Notch(t)) AND (Not P53(t))

ERK(t+1) = (Stat3(t) OR Ras(t)) AND (Not PTEN(t))

JNK(t+1) = (IL-1(t) OR TNF-α(t)) AND (Not PTEN(t))

C． **Ordinary differential equation (ODEs) for endogenous network from Table S2.**

$$\frac{dx_{1}}{dt}=\frac{10 {x_{2}}^{3}}{\left( 10 {x_{2}}^{3}+1 \right) \left( 10 {x_{3}}^{3}+1 \right)}-x_{1}$$

$$\frac{dx_{2}}{dt}=\frac{10 {x_{1}}^{3}}{\left( 10 {x_{1}}^{3}+1 \right) \left( 10 {x_{3}}^{3}+10 {x_{5}}^{3}+1 \right)}-x_{2}$$

$$\frac{dx_{3}}{dt}=\frac{10 {x_{4}}^{3}+10 {x_{29}}^{3}+10 {x_{30}}^{3}}{\left( 10 {x_{4}}^{3}+10 {x_{29}}^{3}+10 {x_{30}}^{3}+1 \right) \left( 10 {x_{1}}^{3}+10 {x_{2}}^{3}+1 \right)}-x_{3}$$

$$\frac{dx_{4}}{dt}=\frac{10 {x_{3}}^{3}}{10 {x_{3}}^{3}+1}-x_{4}$$

$$\frac{dx_{5}}{dt}=\frac{10 {x_{6}}^{3}}{\left( 10 {x_{6}}^{3}+1 \right) \left( 10 {x_{2}}^{3}+1 \right)}-x_{5}$$

$$\frac{dx_{6}}{dt}=\frac{10 {x_{3}}^{3}+10 {x_{5}}^{3}}{\left( 10 {x_{2}}^{3}+1 \right) \left( 10 {x_{3}}^{3}+10 {x_{5}}^{3}+1 \right)}-x_{6}$$

$$\frac{dx_{7}}{dt}=\frac{10 {x_{3}}^{3}}{\left( 10 {x_{3}}^{3}+1 \right) \left( 10 {x_{27}}^{3}+1 \right)}-x_{7}$$

$$\frac{dx_{8}}{dt}=\frac{10{x_{4}}^{3}+10 {x_{24}}^{3}}{\left( 10 {x_{4}}^{3}+10 {x_{24}}^{3}+1 \right)\left( 10 {x_{2}}^{3}+1 \right)}-x_{8}$$

$$\frac{dx_{9}}{dt}=\frac{10{x_{3}}^{3}+10 {x_{13}}^{3}+10 {x_{26}}^{3}+10 {x_{28}}^{3}}{\left( 10{x_{3}}^{3}+10 {x_{13}}^{3}+10 {x_{26}}^{3}+10 {x_{28}}^{3}+1 \right)\left( 10 {x_{2}}^{3}+10 {x_{10}}^{3}+10 {x_{11}}^{3}+1 \right)}-x_{9}$$

$$\frac{dx_{10}}{dt}=\frac{10{x_{11}}^{3}+10 {x_{14}}^{3}+10 {x_{29}}^{3}}{\left( 10{x_{11}}^{3}+10 {x_{14}}^{3}+10 {x_{29}}^{3}+1 \right) \left( 10 {x_{13}}^{3}+10 {x_{26}}^{3}+1 \right)}-x_{10}$$

$$\frac{dx_{11}}{dt}=\frac{10 {x_{13}}^{3}+10 {x_{24}}^{3}+10 {x_{27}}^{3}}{\left( 10 {x_{13}}^{3}+10 {x_{24}}^{3}+10 {x_{27}}^{3}+1 \right) \left( 10 {x_{26}}^{3}+1 \right)}-x_{11}$$

$$\frac{dx_{12}}{dt}=\frac{10 {x_{6}}^{3}}{\left( 10{x_{6}}^{3}+1 \right)\left( 10 {x_{9}}^{3}+10 {x_{18}}^{3}+1 \right)}-x_{12}$$

$$\frac{dx_{13}}{dt}=\frac{10 {x_{14}}^{3}+10 {x_{24}}^{3}+10 {x_{26}}^{3}}{\left( 10 {x_{14}}^{3}+10 {x_{24}}^{3}+10 {x_{26}}^{3}+1 \right)\left( 10 {x_{11}}^{3}+10 {x_{10}}^{3}+1 \right)}-x_{13}$$

$$\frac{dx_{14}}{dt}=\frac{10 {x_{13}}^{3}+10 {x_{14}}^{3}}{\left( 10 {x_{13}}^{3}+10 {x_{14}}^{3}+1 \right) \left( 10 {x_{10}}^{3}+10 {x_{12}}^{3}+1 \right)}-x_{14}$$

$$\frac{dx_{15}}{dt}=\frac{10 {x_{11}}^{3}}{\left( 10 {x_{11}}^{3}+1 \right) \left( 10 {x_{26}}^{3}+10 {x_{29}}^{3}+1 \right)}-x_{15}$$

$$\frac{dx_{16}}{dt}=\frac{10{x_{24}}^{3}+10 {x_{26}}^{3}}{\left( 10{x_{24}}^{3}+10 {x_{26}}^{3}+1 \right) \left( 10 {x_{11}}^{3}+10 {x_{20}}^{3}+1 \right)}-x_{16}$$

$$\frac{dx_{17}}{dt}=\frac{10 {x_{18}}^{3}+10 {x_{20}}^{3}}{\left( 10 {x_{18}}^{3}+10 {x_{20}}^{3}+1 \right) \left( 10 {x_{16}}^{3}+10 {x_{19}}^{3}+10{x_{26}}^{3}+1 \right)}-x_{17}$$

$$\frac{dx_{18}}{dt}=\frac{10 {x_{17}}^{3}}{\left( 10 {x_{17}}^{3}+1 \right) \left( 10 {x_{16}}^{3}+1 \right)}-x_{18}$$

$$\frac{dx_{19}}{dt}=\frac{10 {x_{4}}^{3}+10 {x_{24}}^{3}+10 {x_{29}}^{3}}{\left( 10 {x_{4}}^{3}+10 {x_{24}}^{3}+10 {x_{29}}^{3}+1 \right) \left( 10 {x_{13}}^{3}+10 {x_{15}}^{3}+1 \right)}-x_{19}$$

$$\frac{dx_{20}}{dt}=\frac{10 {x_{11}}^{3}+10 {x_{13}}^{3}+10 {x_{15}}^{3}}{\left( 10 {x_{11}}^{3}+10 {x_{13}}^{3}+10 {x_{15}}^{3}+1 \right) \left( 10 {x_{19}}^{3}+1 \right)}-x_{20}$$

$$\frac{dx_{21}}{dt}=\frac{10 {x_{24}}^{3}+10 {x_{29}}^{3}+10 {x_{30}}^{3}}{\left( 10 {x_{24}}^{3}+10 {x_{29}}^{3}+10 {x_{30}}^{3}+1 \right) \left( 10 {x_{23}}^{3}+1 \right)}-x_{21}$$

$$\frac{dx_{22}}{dt}=\frac{10 {x_{24}}^{3}+10 {x_{30}}^{3}}{10 {x_{24}}^{3}+10 {x_{30}}^{3}+1}-x_{22}$$

$$\frac{dx_{23}}{dt}=\frac{10 {x_{21}}^{3}}{\left( 10 {x_{21}}^{3}+1 \right) \left( 10 {x_{23}}^{3}+1 \right)}-x_{23}$$

$$\frac{dx_{24}}{dt}=\frac{10 {x_{7}}^{3}+10 {x_{21}}^{3}+10 {x_{30}}^{3}}{\left( 10 {x_{7}}^{3}+10 {x_{21}}^{3}+10 {x_{30}}^{3}+1 \right) \left( 10 {x_{25}}^{3}+1 \right)}-x_{24}$$

$$\frac{dx_{25}}{dt}=\frac{10{x_{24}}^{3}+10 {x_{29}}^{3}+10 {x_{30}}^{3}}{\left( 10{x_{24}}^{3}+10 {x_{29}}^{3}+10 {x_{30}}^{3}+1 \right)\left( 10 {x_{21}}^{3}+10 {x_{22}}^{3}+10 {x_{26}}^{3}+1 \right)}-x_{25}$$

$$\frac{dx_{26}}{dt}=\frac{10{x_{7}}^{3}+10 {x_{28}}^{3}}{\left( 10{x_{7}}^{3}+10 {x_{28}}^{3}+1 \right)\left( 10 {x_{27}}^{3}+1 \right)}-x_{26}$$

$$\frac{dx_{27}}{dt}=\frac{10{x_{11}}^{3}}{\left( 10{x_{11}}^{3}+1 \right) \left( 10 {x_{24}}^{3}+1 \right)}-x_{27}$$

$$\frac{dx_{28}}{dt}=\frac{10 {x_{7}}^{3}+10 {x_{8}}^{3}}{\left( 10 {x_{7}}^{3}+10 {x_{8}}^{3}+1 \right) \left( 10 {x_{11}}^{3}+1 \right)}-x_{11}$$

$$\frac{dx_{29}}{dt}=\frac{10 {x_{4}}^{3}+10 {x_{28}}^{3}}{\left( 10 {x_{4}}^{3}+10 {x_{28}}^{3}+1 \right)\left( 10 {x_{27}}^{3}+1 \right)}-x_{12}$$

$$\frac{dx_{30}}{dt}=\frac{10 {x_{21}}^{3}+10 {x_{22}}^{3}}{\left( 10 {x_{21}}^{3}+10 {x_{22}}^{3}+1 \right)\left( 10 {x_{27}}^{3}+1 \right)}-x_{13}$$

D． **Differential equation calculation results**

Table S3 summarizes the 12 steady states and 47 transition states identified from one million random initial vector simulations. In this representation, 0 and 1 denote low and high expression levels, respectively, and each column corresponds to a steady or transition state.

**Table S3.** 12 stable states derived by differential equation method and gene expression levels (rounded to 2 decimal places)

| **Stable point** | **S1** | **S2** | **S3** | **S4** | **S5** | **S6** | **S7** | **S8** | **S9** | **S10** | **S11** | **S12** |
| --- | --- | --- | --- | --- | --- | --- | --- | --- | --- | --- | --- | --- |
| Sox10 | 0.00 | 0.86 | 0.00 | 0.87 | 0.00 | 0.87 | 0.87 | 0.00 | 0.00 | 0.00 | 0.00 | 0.00 |
| Krox20 | 0.00 | 0.86 | 0.00 | 0.87 | 0.00 | 0.87 | 0.87 | 0.00 | 0.00 | 0.00 | 0.00 | 0.00 |
| c-Jun | 0.87 | 0.06 | 0.96 | 0.00 | 0.00 | 0.00 | 0.00 | 0.00 | 0.00 | 0.00 | 0.00 | 0.00 |
| Stat3 | 0.87 | 0.00 | 0.90 | 0.00 | 0.00 | 0.00 | 0.00 | 0.00 | 0.00 | 0.00 | 0.00 | 0.00 |
| NCAM | 0.89 | 0.00 | 0.89 | 0.00 | 0.00 | 0.00 | 0.00 | 0.00 | 0.87 | 0.87 | 0.87 | 0.00 |
| L1CAM | 0.93 | 0.00 | 0.94 | 0.00 | 0.00 | 0.00 | 0.00 | 0.00 | 0.87 | 0.87 | 0.87 | 0.00 |
| BDNF | 0.12 | 0.00 | 0.90 | 0.00 | 0.00 | 0.00 | 0.00 | 0.00 | 0.00 | 0.00 | 0.00 | 0.00 |
| Notch | 0.87 | 0.12 | 0.94 | 0.00 | 0.00 | 0.00 | 0.00 | 0.00 | 0.00 | 0.00 | 0.00 | 0.00 |
| CyclinD-Cdk4,6 | 0.06 | 0.00 | 0.96 | 0.00 | 0.00 | 0.00 | 0.00 | 0.00 | 0.00 | 0.00 | 0.00 | 0.00 |
| P21 | 0.87 | 0.87 | 0.05 | 0.87 | 0.87 | 0.00 | 0.00 | 0.00 | 0.87 | 0.00 | 0.00 | 0.00 |
| P53 | 0.87 | 0.88 | 0.10 | 0.87 | 0.87 | 0.00 | 0.00 | 0.00 | 0.87 | 0.00 | 0.00 | 0.00 |
| Rb | 0.11 | 0.00 | 0.09 | 0.00 | 0.00 | 0.00 | 0.00 | 0.00 | 0.11 | 0.87 | 0.12 | 0.00 |
| Myc | 0.00 | 0.06 | 0.95 | 0.00 | 0.00 | 0.00 | 0.00 | 0.00 | 0.00 | 0.00 | 0.00 | 0.00 |
| E2F | 0.00 | 0.00 | 0.94 | 0.00 | 0.00 | 0.00 | 0.00 | 0.00 | 0.00 | 0.00 | 0.00 | 0.00 |
| BAD | 0.85 | 0.87 | 0.00 | 0.87 | 0.87 | 0.00 | 0.00 | 0.00 | 0.87 | 0.00 | 0.00 | 0.00 |
| XIAP | 0.00 | 0.06 | 0.12 | 0.00 | 0.00 | 0.00 | 0.00 | 0.00 | 0.00 | 0.00 | 0.00 | 0.00 |
| CASP9 | 0.92 | 0.92 | 0.09 | 0.94 | 0.94 | 0.00 | 0.87 | 0.87 | 0.94 | 0.00 | 0.87 | 0.00 |
| CASP3 | 0.89 | 0.88 | 0.01 | 0.89 | 0.89 | 0.00 | 0.87 | 0.87 | 0.89 | 0.00 | 0.87 | 0.00 |
| Bcl2 | 0.12 | 0.12 | 0.10 | 0.00 | 0.00 | 0.00 | 0.00 | 0.00 | 0.00 | 0.00 | 0.00 | 0.00 |
| BAX | 0.91 | 0.92 | 0.89 | 0.93 | 0.93 | 0.00 | 0.00 | 0.00 | 0.93 | 0.00 | 0.00 | 0.00 |
| TNF-$\alpha$ | 0.02 | 0.57 | 0.58 | 0.00 | 0.00 | 0.00 | 0.00 | 0.00 | 0.00 | 0.00 | 0.00 | 0.00 |
| IL-1 | 0.00 | 0.94 | 0.94 | 0.00 | 0.00 | 0.00 | 0.00 | 0.00 | 0.00 | 0.00 | 0.00 | 0.00 |
| IL-10 | 0.00 | 0.40 | 0.40 | 0.00 | 0.00 | 0.00 | 0.00 | 0.00 | 0.00 | 0.00 | 0.00 | 0.00 |
| NF-$\kappa$B | 0.02 | 0.90 | 0.94 | 0.00 | 0.00 | 0.00 | 0.00 | 0.00 | 0.00 | 0.00 | 0.00 | 0.00 |
| ikB | 0.02 | 0.08 | 0.05 | 0.00 | 0.00 | 0.00 | 0.00 | 0.00 | 0.00 | 0.00 | 0.00 | 0.00 |
| Akt | 0.00 | 0.00 | 0.94 | 0.00 | 0.00 | 0.00 | 0.00 | 0.00 | 0.00 | 0.00 | 0.00 | 0.00 |
| PTEN | 0.87 | 0.11 | 0.00 | 0.87 | 0.87 | 0.00 | 0.00 | 0.00 | 0.87 | 0.00 | 0.00 | 0.00 |
| Ras | 0.12 | 0.00 | 0.93 | 0.00 | 0.00 | 0.00 | 0.00 | 0.00 | 0.00 | 0.00 | 0.00 | 0.00 |
| ERK | 0.12 | 0.00 | 0.94 | 0.00 | 0.00 | 0.00 | 0.00 | 0.00 | 0.00 | 0.00 | 0.00 | 0.00 |
| JNK | 0.00 | 0.90 | 0.91 | 0.00 | 0.00 | 0.00 | 0.00 | 0.00 | 0.00 | 0.00 | 0.00 | 0.00 |

Table S4 shows the 47 transition states derived from differential equation calculations, which link distinct steady states. Consistent with the above description, 0 denotes low expression of the corresponding factor, while 1 denotes high expression.

**Table S4.** 47 transition states derived via the differential equation method; each column corresponds to one transition state, with values rounded to two decimal places.

| **Transition point** | **T1** | **T2** | **T3** | **T4** | **T5** | **T6** | **T7** | **T8** | **T9** | **T10** | **T11** | **T12** |
| --- | --- | --- | --- | --- | --- | --- | --- | --- | --- | --- | --- | --- |
| Sox10 | 0.41 | 0.41 | 0.00 | 0.52 | 0.41 | 0.41 | 0.00 | 0.00 | 0.00 | 0.87 | 0.87 | 0.41 |
| Krox20 | 0.41 | 0.41 | 0.00 | 0.52 | 0.41 | 0.41 | 0.00 | 0.00 | 0.00 | 0.87 | 0.87 | 0.41 |
| c-Jun | 0.00 | 0.07 | 0.87 | 0.23 | 0.07 | 0.07 | 0.37 | 0.18 | 0.17 | 0.01 | 0.01 | 0.07 |
| Stat3 | 0.00 | 0.00 | 0.87 | 0.11 | 0.00 | 0.00 | 0.33 | 0.05 | 0.05 | 0.00 | 0.00 | 0.00 |
| NCAM | 0.00 | 0.00 | 0.89 | 0.00 | 0.00 | 0.00 | 0.87 | 0.00 | 0.40 | 0.00 | 0.00 | 0.00 |
| L1CAM | 0.00 | 0.00 | 0.93 | 0.05 | 0.00 | 0.00 | 0.88 | 0.05 | 0.40 | 0.00 | 0.00 | 0.00 |
| BDNF | 0.00 | 0.00 | 0.28 | 0.11 | 0.00 | 0.00 | 0.21 | 0.03 | 0.03 | 0.00 | 0.00 | 0.00 |
| Notch | 0.00 | 0.19 | 0.87 | 0.37 | 0.08 | 0.08 | 0.41 | 0.32 | 0.25 | 0.02 | 0.04 | 0.14 |
| CyclinD-Cdk4,6 | 0.00 | 0.00 | 0.09 | 0.01 | 0.02 | 0.02 | 0.11 | 0.01 | 0.02 | 0.00 | 0.00 | 0.00 |
| P21 | 0.87 | 0.66 | 0.78 | 0.87 | 0.04 | 0.04 | 0.52 | 0.66 | 0.50 | 0.04 | 0.66 | 0.50 |
| P53 | 0.87 | 0.58 | 0.70 | 0.88 | 0.16 | 0.16 | 0.47 | 0.58 | 0.46 | 0.16 | 0.58 | 0.46 |
| Rb | 0.00 | 0.00 | 0.13 | 0.00 | 0.00 | 0.00 | 0.13 | 0.00 | 0.05 | 0.00 | 0.00 | 0.00 |
| Myc | 0.00 | 0.06 | 0.03 | 0.06 | 0.14 | 0.14 | 0.07 | 0.06 | 0.08 | 0.14 | 0.06 | 0.08 |
| E2F | 0.00 | 0.00 | 0.00 | 0.00 | 0.03 | 0.03 | 0.00 | 0.00 | 0.00 | 0.03 | 0.00 | 0.00 |
| BAD | 0.87 | 0.66 | 0.63 | 0.87 | 0.04 | 0.04 | 0.47 | 0.66 | 0.50 | 0.04 | 0.66 | 0.50 |
| XIAP | 0.00 | 0.04 | 0.03 | 0.06 | 0.14 | 0.14 | 0.06 | 0.04 | 0.05 | 0.14 | 0.04 | 0.05 |
| CASP9 | 0.94 | 0.92 | 0.78 | 0.92 | 0.80 | 0.00 | 0.80 | 0.92 | 0.90 | 0.80 | 0.92 | 0.90 |
| CASP3 | 0.89 | 0.89 | 0.83 | 0.88 | 0.82 | 0.00 | 0.84 | 0.89 | 0.88 | 0.82 | 0.89 | 0.88 |
| Bcl2 | 0.00 | 0.08 | 0.25 | 0.12 | 0.14 | 0.14 | 0.21 | 0.08 | 0.11 | 0.14 | 0.08 | 0.11 |
| BAX | 0.93 | 0.82 | 0.74 | 0.92 | 0.06 | 0.06 | 0.62 | 0.82 | 0.68 | 0.06 | 0.82 | 0.68 |
| TNF-$\alpha$ | 0.00 | 0.35 | 0.32 | 0.57 | 0.26 | 0.26 | 0.31 | 0.35 | 0.32 | 0.26 | 0.35 | 0.32 |
| IL-1 | 0.00 | 0.41 | 0.24 | 0.94 | 0.27 | 0.27 | 0.31 | 0.41 | 0.35 | 0.27 | 0.41 | 0.35 |
| IL-10 | 0.00 | 0.26 | 0.22 | 0.40 | 0.15 | 0.15 | 0.21 | 0.26 | 0.22 | 0.15 | 0.26 | 0.22 |
| NF-$\kappa$B | 0.00 | 0.36 | 0.31 | 0.90 | 0.26 | 0.26 | 0.32 | 0.36 | 0.32 | 0.26 | 0.36 | 0.32 |
| ikB | 0.00 | 0.19 | 0.24 | 0.08 | 0.20 | 0.20 | 0.21 | 0.19 | 0.20 | 0.20 | 0.19 | 0.20 |
| Akt | 0.00 | 0.00 | 0.07 | 0.01 | 0.00 | 0.00 | 0.10 | 0.00 | 0.00 | 0.00 | 0.00 | 0.00 |
| PTEN | 0.87 | 0.45 | 0.59 | 0.11 | 0.04 | 0.04 | 0.39 | 0.45 | 0.38 | 0.04 | 0.45 | 0.38 |
| Ras | 0.00 | 0.02 | 0.20 | 0.04 | 0.01 | 0.01 | 0.21 | 0.09 | 0.06 | 0.00 | 0.00 | 0.01 |
| ERK | 0.00 | 0.00 | 0.28 | 0.01 | 0.00 | 0.00 | 0.20 | 0.00 | 0.00 | 0.00 | 0.00 | 0.00 |
| JNK | 0.00 | 0.28 | 0.10 | 0.90 | 0.27 | 0.27 | 0.24 | 0.28 | 0.28 | 0.27 | 0.28 | 0.28 |

| **Transition point** | **T13** | **T14** | **T15** | **T16** | **T17** | **T18** | **T19** | **T20** | **T21** | **T22** | **T23** | **T24** |
| --- | --- | --- | --- | --- | --- | --- | --- | --- | --- | --- | --- | --- |
| Sox10 | 0.00 | 0.00 | 0.87 | 0.41 | 0.87 | 0.00 | 0.00 | 0.00 | 0.00 | 0.00 | 0.87 | 0.00 |
| Krox20 | 0.00 | 0.00 | 0.87 | 0.41 | 0.87 | 0.00 | 0.00 | 0.00 | 0.00 | 0.00 | 0.87 | 0.00 |
| c-Jun | 0.18 | 0.18 | 0.01 | 0.07 | 0.01 | 0.00 | 0.17 | 0.37 | 0.39 | 0.17 | 0.00 | 0.17 |
| Stat3 | 0.05 | 0.05 | 0.00 | 0.00 | 0.00 | 0.00 | 0.05 | 0.34 | 0.38 | 0.05 | 0.00 | 0.05 |
| NCAM | 0.40 | 0.87 | 0.00 | 0.00 | 0.00 | 0.87 | 0.87 | 0.87 | 0.87 | 0.00 | 0.00 | 0.00 |
| L1CAM | 0.40 | 0.87 | 0.00 | 0.00 | 0.00 | 0.87 | 0.87 | 0.88 | 0.88 | 0.05 | 0.00 | 0.05 |
| BDNF | 0.03 | 0.03 | 0.00 | 0.00 | 0.00 | 0.00 | 0.03 | 0.17 | 0.22 | 0.03 | 0.00 | 0.05 |
| Notch | 0.32 | 0.32 | 0.02 | 0.08 | 0.03 | 0.00 | 0.25 | 0.46 | 0.35 | 0.25 | 0.00 | 0.14 |
| CyclinD-Cdk4,6 | 0.01 | 0.01 | 0.00 | 0.02 | 0.00 | 0.00 | 0.02 | 0.06 | 0.16 | 0.02 | 0.00 | 0.07 |
| P21 | 0.66 | 0.66 | 0.04 | 0.04 | 0.50 | 0.41 | 0.50 | 0.67 | 0.45 | 0.50 | 0.41 | 0.04 |
| P53 | 0.58 | 0.58 | 0.16 | 0.16 | 0.46 | 0.41 | 0.46 | 0.58 | 0.42 | 0.46 | 0.41 | 0.16 |
| Rb | 0.05 | 0.11 | 0.00 | 0.00 | 0.00 | 0.11 | 0.11 | 0.11 | 0.15 | 0.00 | 0.00 | 0.00 |
| Myc | 0.06 | 0.06 | 0.14 | 0.14 | 0.08 | 0.00 | 0.08 | 0.05 | 0.01 | 0.08 | 0.00 | 0.14 |
| E2F | 0.00 | 0.00 | 0.03 | 0.03 | 0.00 | 0.00 | 0.00 | 0.00 | 0.00 | 0.00 | 0.00 | 0.03 |
| BAD | 0.66 | 0.66 | 0.04 | 0.04 | 0.50 | 0.41 | 0.50 | 0.64 | 0.37 | 0.50 | 0.41 | 0.04 |
| XIAP | 0.04 | 0.04 | 0.14 | 0.14 | 0.05 | 0.00 | 0.05 | 0.04 | 0.01 | 0.05 | 0.00 | 0.14 |
| CASP9 | 0.92 | 0.92 | 0.00 | 0.43 | 0.90 | 0.90 | 0.90 | 0.90 | 0.72 | 0.90 | 0.90 | 0.00 |
| CASP3 | 0.89 | 0.89 | 0.00 | 0.44 | 0.88 | 0.88 | 0.88 | 0.88 | 0.79 | 0.88 | 0.88 | 0.00 |
| Bcl2 | 0.08 | 0.08 | 0.14 | 0.14 | 0.11 | 0.00 | 0.11 | 0.13 | 0.26 | 0.11 | 0.00 | 0.14 |
| BAX | 0.82 | 0.82 | 0.06 | 0.06 | 0.68 | 0.58 | 0.68 | 0.80 | 0.47 | 0.68 | 0.58 | 0.06 |
| TNF-$\alpha$ | 0.35 | 0.35 | 0.26 | 0.26 | 0.32 | 0.00 | 0.32 | 0.35 | 0.11 | 0.32 | 0.00 | 0.26 |
| IL-1 | 0.41 | 0.41 | 0.27 | 0.27 | 0.35 | 0.00 | 0.35 | 0.39 | 0.01 | 0.35 | 0.00 | 0.27 |
| IL-10 | 0.26 | 0.26 | 0.15 | 0.15 | 0.22 | 0.00 | 0.22 | 0.25 | 0.01 | 0.22 | 0.00 | 0.15 |
| NF-$\kappa$B | 0.36 | 0.36 | 0.26 | 0.26 | 0.32 | 0.00 | 0.32 | 0.36 | 0.11 | 0.32 | 0.00 | 0.26 |
| ikB | 0.19 | 0.19 | 0.20 | 0.20 | 0.20 | 0.00 | 0.20 | 0.20 | 0.11 | 0.20 | 0.00 | 0.20 |
| Akt | 0.00 | 0.00 | 0.00 | 0.00 | 0.00 | 0.00 | 0.00 | 0.05 | 0.09 | 0.00 | 0.00 | 0.00 |
| PTEN | 0.45 | 0.45 | 0.04 | 0.04 | 0.38 | 0.41 | 0.38 | 0.45 | 0.41 | 0.38 | 0.41 | 0.04 |
| Ras | 0.09 | 0.09 | 0.00 | 0.01 | 0.00 | 0.00 | 0.06 | 0.17 | 0.21 | 0.06 | 0.00 | 0.03 |
| ERK | 0.00 | 0.00 | 0.00 | 0.00 | 0.00 | 0.00 | 0.00 | 0.15 | 0.22 | 0.00 | 0.00 | 0.00 |
| JNK | 0.28 | 0.28 | 0.27 | 0.27 | 0.28 | 0.00 | 0.28 | 0.26 | 0.01 | 0.28 | 0.00 | 0.27 |

| **Transition point** | **T25** | **T26** | **T27** | **T28** | **T29** | **T30** | **T31** | **T32** | **T33** | **T34** | **T35** | **T36** |
| --- | --- | --- | --- | --- | --- | --- | --- | --- | --- | --- | --- | --- |
| Sox10 | 0.00 | 0.00 | 0.00 | 0.00 | 0.00 | 0.00 | 0.00 | 0.41 | 0.41 | 0.00 | 0.00 | 0.00 |
| Krox20 | 0.00 | 0.00 | 0.00 | 0.00 | 0.00 | 0.00 | 0.00 | 0.41 | 0.41 | 0.00 | 0.00 | 0.00 |
| c-Jun | 0.17 | 0.17 | 0.41 | 0.17 | 0.17 | 0.00 | 0.00 | 0.00 | 0.00 | 0.17 | 0.00 | 0.17 |
| Stat3 | 0.05 | 0.05 | 0.41 | 0.05 | 0.05 | 0.00 | 0.00 | 0.00 | 0.00 | 0.05 | 0.00 | 0.05 |
| NCAM | 0.87 | 0.00 | 0.87 | 0.40 | 0.87 | 0.41 | 0.00 | 0.00 | 0.00 | 0.40 | 0.41 | 0.40 |
| L1CAM | 0.87 | 0.05 | 0.88 | 0.40 | 0.87 | 0.41 | 0.00 | 0.00 | 0.00 | 0.40 | 0.41 | 0.40 |
| BDNF | 0.05 | 0.05 | 0.05 | 0.05 | 0.05 | 0.00 | 0.00 | 0.00 | 0.00 | 0.05 | 0.00 | 0.05 |
| Notch | 0.14 | 0.14 | 0.41 | 0.14 | 0.14 | 0.00 | 0.00 | 0.00 | 0.00 | 0.14 | 0.00 | 0.14 |
| CyclinD-Cdk4,6 | 0.07 | 0.07 | 0.03 | 0.07 | 0.07 | 0.00 | 0.00 | 0.00 | 0.00 | 0.07 | 0.00 | 0.07 |
| P21 | 0.04 | 0.04 | 0.87 | 0.04 | 0.04 | 0.41 | 0.41 | 0.00 | 0.41 | 0.04 | 0.87 | 0.04 |
| P53 | 0.16 | 0.16 | 0.87 | 0.16 | 0.16 | 0.41 | 0.41 | 0.00 | 0.41 | 0.16 | 0.87 | 0.16 |
| Rb | 0.86 | 0.00 | 0.11 | 0.06 | 0.13 | 0.05 | 0.00 | 0.00 | 0.00 | 0.40 | 0.05 | 0.22 |
| Myc | 0.14 | 0.14 | 0.00 | 0.14 | 0.14 | 0.00 | 0.00 | 0.00 | 0.00 | 0.14 | 0.00 | 0.14 |
| E2F | 0.00 | 0.03 | 0.00 | 0.03 | 0.03 | 0.00 | 0.00 | 0.00 | 0.00 | 0.02 | 0.00 | 0.02 |
| BAD | 0.04 | 0.04 | 0.87 | 0.04 | 0.04 | 0.41 | 0.41 | 0.00 | 0.41 | 0.04 | 0.87 | 0.04 |
| XIAP | 0.14 | 0.14 | 0.00 | 0.14 | 0.14 | 0.00 | 0.00 | 0.00 | 0.00 | 0.14 | 0.00 | 0.14 |
| CASP9 | 0.00 | 0.80 | 0.94 | 0.80 | 0.80 | 0.90 | 0.90 | 0.00 | 0.90 | 0.00 | 0.94 | 0.43 |
| CASP3 | 0.00 | 0.82 | 0.89 | 0.82 | 0.82 | 0.88 | 0.88 | 0.00 | 0.88 | 0.00 | 0.89 | 0.44 |
| Bcl2 | 0.14 | 0.14 | 0.06 | 0.14 | 0.14 | 0.00 | 0.00 | 0.00 | 0.00 | 0.14 | 0.00 | 0.14 |
| BAX | 0.06 | 0.06 | 0.93 | 0.06 | 0.06 | 0.58 | 0.58 | 0.00 | 0.58 | 0.06 | 0.93 | 0.06 |
| TNF-$\alpha$ | 0.26 | 0.26 | 0.00 | 0.26 | 0.26 | 0.00 | 0.00 | 0.00 | 0.00 | 0.26 | 0.00 | 0.26 |
| IL-1 | 0.27 | 0.27 | 0.00 | 0.27 | 0.27 | 0.00 | 0.00 | 0.00 | 0.00 | 0.27 | 0.00 | 0.27 |
| IL-10 | 0.15 | 0.15 | 0.00 | 0.15 | 0.15 | 0.00 | 0.00 | 0.00 | 0.00 | 0.15 | 0.00 | 0.15 |
| NF-$\kappa$B | 0.26 | 0.26 | 0.00 | 0.26 | 0.26 | 0.00 | 0.00 | 0.00 | 0.00 | 0.26 | 0.00 | 0.26 |
| ikB | 0.20 | 0.20 | 0.00 | 0.20 | 0.20 | 0.00 | 0.00 | 0.00 | 0.00 | 0.20 | 0.00 | 0.20 |
| Akt | 0.00 | 0.00 | 0.00 | 0.00 | 0.00 | 0.00 | 0.00 | 0.00 | 0.00 | 0.00 | 0.00 | 0.00 |
| PTEN | 0.04 | 0.04 | 0.87 | 0.04 | 0.04 | 0.41 | 0.41 | 0.00 | 0.41 | 0.04 | 0.87 | 0.04 |
| Ras | 0.03 | 0.03 | 0.05 | 0.03 | 0.03 | 0.00 | 0.00 | 0.00 | 0.00 | 0.03 | 0.00 | 0.03 |
| ERK | 0.00 | 0.00 | 0.05 | 0.00 | 0.00 | 0.00 | 0.00 | 0.00 | 0.00 | 0.00 | 0.00 | 0.00 |
| JNK | 0.27 | 0.27 | 0.00 | 0.27 | 0.27 | 0.00 | 0.00 | 0.00 | 0.00 | 0.27 | 0.00 | 0.27 |

| **Transition point** | **T37** | **T38** | **T39** | **T40** | **T41** | **T42** | **T43** | **T44** | **T45** | **T46** | **T47** |
| --- | --- | --- | --- | --- | --- | --- | --- | --- | --- | --- | --- |
| Sox10 | 0.00 | 0.41 | 0.00 | 0.87 | 0.00 | 0.41 | 0.00 | 0.87 | 0.00 | 0.00 | 0.00 |
| Krox20 | 0.00 | 0.41 | 0.00 | 0.87 | 0.00 | 0.41 | 0.00 | 0.87 | 0.00 | 0.00 | 0.00 |
| c-Jun | 0.00 | 0.00 | 0.00 | 0.01 | 0.17 | 0.00 | 0.17 | 0.00 | 0.00 | 0.00 | 0.00 |
| Stat3 | 0.00 | 0.00 | 0.00 | 0.00 | 0.05 | 0.00 | 0.05 | 0.00 | 0.00 | 0.00 | 0.00 |
| NCAM | 0.41 | 0.00 | 0.41 | 0.00 | 0.87 | 0.00 | 0.00 | 0.00 | 0.41 | 0.87 | 0.00 |
| L1CAM | 0.41 | 0.00 | 0.41 | 0.00 | 0.87 | 0.00 | 0.05 | 0.00 | 0.41 | 0.87 | 0.00 |
| BDNF | 0.00 | 0.00 | 0.00 | 0.00 | 0.05 | 0.00 | 0.05 | 0.00 | 0.00 | 0.00 | 0.00 |
| Notch | 0.00 | 0.00 | 0.00 | 0.02 | 0.14 | 0.00 | 0.14 | 0.00 | 0.00 | 0.00 | 0.00 |
| CyclinD-Cdk4,6 | 0.00 | 0.00 | 0.00 | 0.00 | 0.07 | 0.00 | 0.07 | 0.00 | 0.00 | 0.00 | 0.00 |
| P21 | 0.00 | 0.00 | 0.00 | 0.04 | 0.04 | 0.00 | 0.04 | 0.00 | 0.00 | 0.00 | 0.00 |
| P53 | 0.00 | 0.00 | 0.00 | 0.16 | 0.16 | 0.00 | 0.16 | 0.00 | 0.00 | 0.00 | 0.00 |
| Rb | 0.05 | 0.00 | 0.41 | 0.00 | 0.47 | 0.00 | 0.00 | 0.00 | 0.24 | 0.51 | 0.00 |
| Myc | 0.00 | 0.00 | 0.00 | 0.14 | 0.14 | 0.00 | 0.14 | 0.00 | 0.00 | 0.00 | 0.00 |
| E2F | 0.00 | 0.00 | 0.00 | 0.03 | 0.01 | 0.00 | 0.03 | 0.00 | 0.00 | 0.00 | 0.00 |
| BAD | 0.00 | 0.00 | 0.00 | 0.04 | 0.04 | 0.00 | 0.04 | 0.00 | 0.00 | 0.00 | 0.00 |
| XIAP | 0.00 | 0.00 | 0.00 | 0.14 | 0.14 | 0.00 | 0.14 | 0.00 | 0.00 | 0.00 | 0.00 |
| CASP9 | 0.87 | 0.41 | 0.00 | 0.43 | 0.43 | 0.87 | 0.43 | 0.41 | 0.41 | 0.41 | 0.41 |
| CASP3 | 0.87 | 0.41 | 0.00 | 0.44 | 0.44 | 0.87 | 0.44 | 0.41 | 0.41 | 0.41 | 0.41 |
| Bcl2 | 0.00 | 0.00 | 0.00 | 0.14 | 0.14 | 0.00 | 0.14 | 0.00 | 0.00 | 0.00 | 0.00 |
| BAX | 0.00 | 0.00 | 0.00 | 0.06 | 0.06 | 0.00 | 0.06 | 0.00 | 0.00 | 0.00 | 0.00 |
| TNF-$\alpha$ | 0.00 | 0.00 | 0.00 | 0.26 | 0.26 | 0.00 | 0.26 | 0.00 | 0.00 | 0.00 | 0.00 |
| IL-1 | 0.00 | 0.00 | 0.00 | 0.27 | 0.27 | 0.00 | 0.27 | 0.00 | 0.00 | 0.00 | 0.00 |
| IL-10 | 0.00 | 0.00 | 0.00 | 0.15 | 0.15 | 0.00 | 0.15 | 0.00 | 0.00 | 0.00 | 0.00 |
| NF-$\kappa$B | 0.00 | 0.00 | 0.00 | 0.26 | 0.26 | 0.00 | 0.26 | 0.00 | 0.00 | 0.00 | 0.00 |
| ikB | 0.00 | 0.00 | 0.00 | 0.20 | 0.20 | 0.00 | 0.20 | 0.00 | 0.00 | 0.00 | 0.00 |
| Akt | 0.00 | 0.00 | 0.00 | 0.00 | 0.00 | 0.00 | 0.00 | 0.00 | 0.00 | 0.00 | 0.00 |
| PTEN | 0.00 | 0.00 | 0.00 | 0.04 | 0.04 | 0.00 | 0.04 | 0.00 | 0.00 | 0.00 | 0.00 |
| Ras | 0.00 | 0.00 | 0.00 | 0.00 | 0.03 | 0.00 | 0.03 | 0.00 | 0.00 | 0.00 | 0.00 |
| ERK | 0.00 | 0.00 | 0.00 | 0.00 | 0.00 | 0.00 | 0.00 | 0.00 | 0.00 | 0.00 | 0.00 |
| JNK | 0.00 | 0.00 | 0.00 | 0.27 | 0.27 | 0.00 | 0.27 | 0.00 | 0.00 | 0.00 | 0.0 |

**Table S5.** 37 attractors calculated by the Boolean algebra method; each column represents one attractor, with values rounded to two decimal places.

| **Boolean results** | **B1** | **B2** | **B3** | **B4** | **B5** | **B6** | **B7** | **B8** | **B9** | **B10** | **B11** | **B12** |
| --- | --- | --- | --- | --- | --- | --- | --- | --- | --- | --- | --- | --- |
| Sox10 | 0.50 | 0.50 | 0.00 | 0.00 | 0.00 | 0.00 | 0.00 | 1.00 | 0.00 | 1.00 | 0.00 | 0.50 |
| Krox20 | 0.50 | 0.50 | 0.00 | 0.00 | 0.00 | 0.00 | 0.00 | 1.00 | 0.00 | 1.00 | 0.00 | 0.50 |
| c-Jun | 0.00 | 0.00 | 1.00 | 1.00 | 1.00 | 1.00 | 0.50 | 0.00 | 0.50 | 0.00 | 0.00 | 0.00 |
| Stat3 | 0.00 | 0.00 | 1.00 | 1.00 | 1.00 | 1.00 | 0.50 | 0.00 | 0.50 | 0.00 | 0.00 | 0.00 |
| NCAM | 0.00 | 0.00 | 1.00 | 1.00 | 1.00 | 1.00 | 1.00 | 0.00 | 0.50 | 0.00 | 0.50 | 0.00 |
| L1CAM | 0.00 | 0.00 | 1.00 | 1.00 | 1.00 | 1.00 | 1.00 | 0.00 | 0.50 | 0.00 | 0.50 | 0.00 |
| BDNF | 0.00 | 0.00 | 1.00 | 0.00 | 0.67 | 0.67 | 0.00 | 0.00 | 0.00 | 0.00 | 0.00 | 0.00 |
| Notch | 0.50 | 0.33 | 1.00 | 1.00 | 1.00 | 1.00 | 0.50 | 0.00 | 0.50 | 0.00 | 0.00 | 0.00 |
| CyclinD-Cdk4,6 | 0.00 | 0.00 | 1.00 | 0.00 | 0.33 | 0.67 | 0.00 | 0.00 | 0.00 | 0.00 | 0.00 | 0.00 |
| P21 | 1.00 | 0.67 | 0.00 | 1.00 | 0.00 | 0.00 | 1.00 | 0.67 | 1.00 | 1.00 | 1.00 | 1.00 |
| P53 | 1.00 | 0.67 | 0.00 | 1.00 | 0.67 | 0.33 | 1.00 | 0.67 | 1.00 | 1.00 | 1.00 | 1.00 |
| Rb | 0.00 | 0.00 | 0.00 | 0.00 | 0.67 | 0.33 | 0.00 | 0.00 | 0.00 | 0.00 | 0.00 | 0.00 |
| Myc | 0.00 | 0.00 | 1.00 | 0.00 | 0.33 | 0.67 | 0.00 | 0.00 | 0.00 | 0.00 | 0.00 | 0.00 |
| E2F | 0.00 | 0.00 | 1.00 | 0.00 | 0.00 | 0.67 | 0.00 | 0.00 | 0.00 | 0.00 | 0.00 | 0.00 |
| BAD | 1.00 | 0.67 | 0.00 | 1.00 | 0.00 | 0.00 | 1.00 | 0.67 | 1.00 | 1.00 | 1.00 | 1.00 |
| XIAP | 0.00 | 0.33 | 0.00 | 0.00 | 0.00 | 0.33 | 0.00 | 0.33 | 0.00 | 0.00 | 0.00 | 0.00 |
| CASP9 | 1.00 | 0.00 | 0.00 | 1.00 | 0.00 | 0.00 | 1.00 | 0.00 | 1.00 | 1.00 | 1.00 | 1.00 |
| CASP3 | 1.00 | 0.00 | 0.00 | 1.00 | 0.00 | 0.00 | 1.00 | 0.00 | 1.00 | 1.00 | 1.00 | 1.00 |
| Bcl2 | 0.00 | 0.33 | 0.00 | 0.00 | 0.67 | 0.33 | 0.00 | 0.33 | 0.00 | 0.00 | 0.00 | 0.00 |
| BAX | 1.00 | 0.67 | 1.00 | 1.00 | 0.33 | 0.33 | 1.00 | 0.67 | 1.00 | 1.00 | 1.00 | 1.00 |
| TNF-$\alpha$ | 0.67 | 0.67 | 0.67 | 0.00 | 0.67 | 0.67 | 0.00 | 0.67 | 0.00 | 0.67 | 0.00 | 0.00 |
| IL-1 | 1.00 | 1.00 | 1.00 | 0.00 | 1.00 | 1.00 | 0.00 | 1.00 | 0.00 | 1.00 | 0.00 | 0.00 |
| IL-10 | 0.33 | 0.33 | 0.33 | 0.00 | 0.33 | 0.33 | 0.00 | 0.33 | 0.00 | 0.33 | 0.00 | 0.00 |
| NF-$\kappa$B | 1.00 | 0.67 | 1.00 | 0.00 | 0.67 | 0.67 | 0.00 | 0.67 | 0.00 | 1.00 | 0.00 | 0.00 |
| ikB | 0.00 | 0.00 | 0.00 | 0.00 | 0.00 | 0.00 | 0.00 | 0.00 | 0.00 | 0.00 | 0.00 | 0.00 |
| Akt | 0.00 | 0.00 | 1.00 | 0.00 | 0.33 | 0.67 | 0.00 | 0.00 | 0.00 | 0.00 | 0.00 | 0.00 |
| PTEN | 0.00 | 0.33 | 0.00 | 1.00 | 0.33 | 0.33 | 1.00 | 0.33 | 1.00 | 0.00 | 1.00 | 1.00 |
| Ras | 0.00 | 0.00 | 1.00 | 0.00 | 0.33 | 0.67 | 0.00 | 0.00 | 0.00 | 0.00 | 0.00 | 0.00 |
| ERK | 0.00 | 0.00 | 1.00 | 0.00 | 0.67 | 0.67 | 0.00 | 0.00 | 0.00 | 0.00 | 0.00 | 0.00 |
| JNK | 1.00 | 0.67 | 1.00 | 0.00 | 0.67 | 0.67 | 0.00 | 0.67 | 0.00 | 1.00 | 0.00 | 0.00 |

| **Boolean results** | **B13** | **B14** | **B15** | **B16** | **B17** | **B18** | **B19** | **B20** | **B21** | **B22** | **B23** | **B24** |
| --- | --- | --- | --- | --- | --- | --- | --- | --- | --- | --- | --- | --- |
| Sox10 | 0.50 | 0.00 | 1.00 | 1.00 | 0.50 | 0.00 | 0.00 | 0.50 | 1.00 | 0.00 | 0.00 | 0.00 |
| Krox20 | 0.50 | 0.00 | 1.00 | 1.00 | 0.50 | 0.00 | 0.00 | 0.50 | 1.00 | 0.00 | 0.00 | 0.00 |
| c-Jun | 0.00 | 0.00 | 0.00 | 0.00 | 0.00 | 0.00 | 0.00 | 0.00 | 0.00 | 0.00 | 0.00 | 0.00 |
| Stat3 | 0.00 | 0.00 | 0.00 | 0.00 | 0.00 | 0.00 | 0.00 | 0.00 | 0.00 | 0.00 | 0.00 | 0.00 |
| NCAM | 0.00 | 1.00 | 0.00 | 0.00 | 0.00 | 1.00 | 0.00 | 0.00 | 0.00 | 0.00 | 0.50 | 0.50 |
| L1CAM | 0.00 | 1.00 | 0.00 | 0.00 | 0.00 | 1.00 | 0.00 | 0.00 | 0.00 | 0.00 | 0.50 | 0.50 |
| BDNF | 0.00 | 0.00 | 0.00 | 0.00 | 0.00 | 0.00 | 0.00 | 0.00 | 0.00 | 0.00 | 0.00 | 0.00 |
| Notch | 0.00 | 0.00 | 0.00 | 0.00 | 0.00 | 0.00 | 0.00 | 0.00 | 0.00 | 0.00 | 0.00 | 0.00 |
| CyclinD-Cdk4,6 | 0.00 | 0.00 | 0.00 | 0.00 | 0.00 | 0.00 | 0.00 | 0.00 | 0.00 | 0.00 | 0.00 | 0.00 |
| P21 | 0.50 | 0.50 | 0.50 | 1.00 | 0.50 | 1.00 | 1.00 | 0.00 | 0.00 | 0.50 | 0.50 | 0.50 |
| P53 | 0.50 | 0.50 | 0.50 | 1.00 | 0.50 | 1.00 | 1.00 | 0.00 | 0.00 | 0.50 | 0.50 | 0.50 |
| Rb | 0.00 | 0.00 | 0.00 | 0.00 | 0.00 | 0.00 | 0.00 | 0.00 | 0.00 | 0.00 | 0.00 | 0.00 |
| Myc | 0.00 | 0.00 | 0.00 | 0.00 | 0.00 | 0.00 | 0.00 | 0.00 | 0.00 | 0.00 | 0.00 | 0.00 |
| E2F | 0.00 | 0.00 | 0.00 | 0.00 | 0.00 | 0.00 | 0.00 | 0.00 | 0.00 | 0.00 | 0.00 | 0.00 |
| BAD | 0.50 | 0.50 | 0.50 | 1.00 | 0.50 | 1.00 | 1.00 | 0.00 | 0.00 | 0.50 | 0.50 | 0.50 |
| XIAP | 0.00 | 0.00 | 0.00 | 0.00 | 0.00 | 0.00 | 0.00 | 0.00 | 0.00 | 0.00 | 0.00 | 0.00 |
| CASP9 | 1.00 | 1.00 | 1.00 | 1.00 | 1.00 | 1.00 | 1.00 | 0.00 | 0.00 | 1.00 | 1.00 | 1.00 |
| CASP3 | 1.00 | 1.00 | 1.00 | 1.00 | 1.00 | 1.00 | 1.00 | 0.00 | 0.00 | 1.00 | 1.00 | 1.00 |
| Bcl2 | 0.00 | 0.00 | 0.00 | 0.00 | 0.00 | 0.00 | 0.00 | 0.00 | 0.00 | 0.00 | 0.00 | 0.00 |
| BAX | 1.00 | 1.00 | 1.00 | 1.00 | 1.00 | 1.00 | 1.00 | 0.00 | 0.00 | 1.00 | 1.00 | 1.00 |
| TNF-$\alpha$ | 0.00 | 0.00 | 0.00 | 0.00 | 0.00 | 0.00 | 0.00 | 0.00 | 0.00 | 0.00 | 0.00 | 0.00 |
| IL-1 | 0.00 | 0.00 | 0.00 | 0.00 | 0.00 | 0.00 | 0.00 | 0.00 | 0.00 | 0.00 | 0.00 | 0.00 |
| IL-10 | 0.00 | 0.00 | 0.00 | 0.00 | 0.00 | 0.00 | 0.00 | 0.00 | 0.00 | 0.00 | 0.00 | 0.00 |
| NF-$\kappa$B | 0.00 | 0.00 | 0.00 | 0.00 | 0.00 | 0.00 | 0.00 | 0.00 | 0.00 | 0.00 | 0.00 | 0.00 |
| ikB | 0.00 | 0.00 | 0.00 | 0.00 | 0.00 | 0.00 | 0.00 | 0.00 | 0.00 | 0.00 | 0.00 | 0.00 |
| Akt | 0.00 | 0.00 | 0.00 | 0.00 | 0.00 | 0.00 | 0.00 | 0.00 | 0.00 | 0.00 | 0.00 | 0.00 |
| PTEN | 0.50 | 0.50 | 0.50 | 1.00 | 0.50 | 1.00 | 1.00 | 0.00 | 0.00 | 0.50 | 0.50 | 0.50 |
| Ras | 0.00 | 0.00 | 0.00 | 0.00 | 0.00 | 0.00 | 0.00 | 0.00 | 0.00 | 0.00 | 0.00 | 0.00 |
| ERK | 0.00 | 0.00 | 0.00 | 0.00 | 0.00 | 0.00 | 0.00 | 0.00 | 0.00 | 0.00 | 0.00 | 0.00 |
| JNK | 0.00 | 0.00 | 0.00 | 0.00 | 0.00 | 0.00 | 0.00 | 0.00 | 0.00 | 0.00 | 0.00 | 0.00 |

| **Boolean results** | **B25** | **B26** | **B27** | **B28** | **B29** | **B30** | **B31** | **B32** | **B33** | **B34** | **B35** | **B36** | **B37** |
| --- | --- | --- | --- | --- | --- | --- | --- | --- | --- | --- | --- | --- | --- |
| Sox10 | 0.50 | 0.50 | 1.00 | 0.00 | 0.00 | 0.00 | 0.00 | 0.50 | 0.00 | 0.00 | 0.00 | 0.00 | 1.00 |
| Krox20 | 0.50 | 0.50 | 1.00 | 0.00 | 0.00 | 0.00 | 0.00 | 0.50 | 0.00 | 0.00 | 0.00 | 0.00 | 1.00 |
| c-Jun | 0.00 | 0.00 | 0.00 | 0.00 | 0.00 | 0.00 | 0.00 | 0.00 | 0.00 | 0.00 | 0.00 | 0.00 | 0.00 |
| Stat3 | 0.00 | 0.00 | 0.00 | 0.00 | 0.00 | 0.00 | 0.00 | 0.00 | 0.00 | 0.00 | 0.00 | 0.00 | 0.00 |
| NCAM | 0.00 | 0.00 | 0.00 | 1.00 | 0.50 | 1.00 | 0.50 | 0.00 | 0.50 | 0.00 | 0.50 | 0.00 | 0.00 |
| L1CAM | 0.00 | 0.00 | 0.00 | 1.00 | 0.50 | 1.00 | 0.50 | 0.00 | 0.50 | 0.00 | 0.50 | 0.00 | 0.00 |
| BDNF | 0.00 | 0.00 | 0.00 | 0.00 | 0.00 | 0.00 | 0.00 | 0.00 | 0.00 | 0.00 | 0.00 | 0.00 | 0.00 |
| Notch | 0.00 | 0.00 | 0.00 | 0.00 | 0.00 | 0.00 | 0.00 | 0.00 | 0.00 | 0.00 | 0.00 | 0.00 | 0.00 |
| CyclinD-Cdk4,6 | 0.00 | 0.00 | 0.00 | 0.00 | 0.00 | 0.00 | 0.00 | 0.00 | 0.00 | 0.00 | 0.00 | 0.00 | 0.00 |
| P21 | 0.00 | 0.00 | 0.00 | 0.00 | 0.00 | 0.00 | 0.00 | 0.00 | 0.00 | 0.00 | 0.00 | 0.00 | 0.00 |
| P53 | 0.00 | 0.00 | 0.00 | 0.00 | 0.00 | 0.00 | 0.00 | 0.00 | 0.00 | 0.00 | 0.00 | 0.00 | 0.00 |
| Rb | 0.00 | 0.00 | 0.00 | 0.50 | 0.50 | 1.00 | 0.00 | 0.00 | 0.50 | 0.00 | 0.00 | 0.00 | 0.00 |
| Myc | 0.00 | 0.00 | 0.00 | 0.00 | 0.00 | 0.00 | 0.00 | 0.00 | 0.00 | 0.00 | 0.00 | 0.00 | 0.00 |
| E2F | 0.00 | 0.00 | 0.00 | 0.00 | 0.00 | 0.00 | 0.00 | 0.00 | 0.00 | 0.00 | 0.00 | 0.00 | 0.00 |
| BAD | 0.00 | 0.00 | 0.00 | 0.00 | 0.00 | 0.00 | 0.00 | 0.00 | 0.00 | 0.00 | 0.00 | 0.00 | 0.00 |
| XIAP | 0.00 | 0.00 | 0.00 | 0.00 | 0.00 | 0.00 | 0.00 | 0.00 | 0.00 | 0.00 | 0.00 | 0.00 | 0.00 |
| CASP9 | 0.50 | 0.50 | 0.50 | 0.50 | 0.00 | 0.00 | 0.50 | 1.00 | 0.50 | 0.50 | 1.00 | 0.00 | 1.00 |
| CASP3 | 0.50 | 0.50 | 0.50 | 0.50 | 0.00 | 0.00 | 0.50 | 1.00 | 0.50 | 0.50 | 1.00 | 0.00 | 1.00 |
| Bcl2 | 0.00 | 0.00 | 0.00 | 0.00 | 0.00 | 0.00 | 0.00 | 0.00 | 0.00 | 0.00 | 0.00 | 0.00 | 0.00 |
| BAX | 0.00 | 0.00 | 0.00 | 0.00 | 0.00 | 0.00 | 0.00 | 0.00 | 0.00 | 0.00 | 0.00 | 0.00 | 0.00 |
| TNF-$\alpha$ | 0.00 | 0.00 | 0.00 | 0.00 | 0.00 | 0.00 | 0.00 | 0.00 | 0.00 | 0.00 | 0.00 | 0.00 | 0.00 |
| IL-1 | 0.00 | 0.00 | 0.00 | 0.00 | 0.00 | 0.00 | 0.00 | 0.00 | 0.00 | 0.00 | 0.00 | 0.00 | 0.00 |
| IL-10 | 0.00 | 0.00 | 0.00 | 0.00 | 0.00 | 0.00 | 0.00 | 0.00 | 0.00 | 0.00 | 0.00 | 0.00 | 0.00 |
| NF-$\kappa$B | 0.00 | 0.00 | 0.00 | 0.00 | 0.00 | 0.00 | 0.00 | 0.00 | 0.00 | 0.00 | 0.00 | 0.00 | 0.00 |
| ikB | 0.00 | 0.00 | 0.00 | 0.00 | 0.00 | 0.00 | 0.00 | 0.00 | 0.00 | 0.00 | 0.00 | 0.00 | 0.00 |
| Akt | 0.00 | 0.00 | 0.00 | 0.00 | 0.00 | 0.00 | 0.00 | 0.00 | 0.00 | 0.00 | 0.00 | 0.00 | 0.00 |
| PTEN | 0.00 | 0.00 | 0.00 | 0.00 | 0.00 | 0.00 | 0.00 | 0.00 | 0.00 | 0.00 | 0.00 | 0.00 | 0.00 |
| Ras | 0.00 | 0.00 | 0.00 | 0.00 | 0.00 | 0.00 | 0.00 | 0.00 | 0.00 | 0.00 | 0.00 | 0.00 | 0.00 |
| ERK | 0.00 | 0.00 | 0.00 | 0.00 | 0.00 | 0.00 | 0.00 | 0.00 | 0.00 | 0.00 | 0.00 | 0.00 | 0.00 |
| JNK | 0.00 | 0.00 | 0.00 | 0.00 | 0.00 | 0.00 | 0.00 | 0.00 | 0.00 | 0.00 | 0.00 | 0.00 | 0.00 |

E． **Literature on obtaining low-throughput data and GEO data processing methods**

**Low-throughput experimental literature**

Low-throughput experiments are unable to cover all genes included in this study. We therefore integrated extensive low-throughput data with the results derived from endogenous network calculations, as illustrated in Figure 3A. Each column corresponds to one experiment; NA denotes genes not included in the experimental assay, 0 denotes genes downregulated in the simulated injury group relative to the healthy control group, and 1 denotes activated genes. The relevant references are listed below [1-19].

**References for Supplementary Material**

[1]Atanasoski S, Shumas S, Dickson C, Scherer SS, Suter U (2001) Differential cyclin D1 requirements of proliferating Schwann cells during development and after injury. Mol Cell Neurosci 18:581-592.

[2]Atanasoski S, Boller D, De Ventura L, Koegel H, Boentert M, Young P, Werner S, Suter U (2006) Cell cycle inhibitors p21 and p16 are required for the regulation of Schwann cell proliferation. Glia 53:147-157.

[3]Belin S, Nawabi H, Wang C, Tang S, Latremoliere A, Warren P, Schorle H, Uncu C, Woolf CJ, He Z, Steen JA (2015) Injury-induced decline of intrinsic regenerative ability revealed by quantitative proteomics. Neuron 86:1000-1014.

[4]Benito C, Davis CM, Gomez-Sanchez JA, Turmaine M, Meijer D, Poli V, Mirsky R, Jessen KR (2017) STAT3 Controls the Long-Term Survival and Phenotype of Repair Schwann Cells during Nerve Regeneration. J Neurosci 37:4255-4269.

[5]Chen H, Xiang J, Wu J, He B, Lin T, Zhu Q, Liu X, Zheng C (2018) Expression patterns and role of PTEN in rat peripheral nerve development and injury. Neurosci Lett 676:78-84.

[6]Christie KJ, Krishnan A, Martinez JA, Purdy K, Singh B, Eaton S, Zochodne D (2014) Enhancing adult nerve regeneration through the knockdown of retinoblastoma protein. Nat Commun 5:3670.

[7]Dubový P, Klusáková I, Hradilová Svíženská I (2014) Inflammatory profiling of Schwann cells in contact with growing axons distal to nerve injury. Biomed Res Int 2014:691041.

[8]Dzreyan V, Eid M, Rodkin S, Pitinova M, Demyanenko S (2022) E2F1 Expression and Apoptosis Initiation in Crayfish and Rat Peripheral Neurons and Glial Cells after Axonal Injury. Int J Mol Sci 23.

[9]Fontana X, Hristova M, Da Costa C, Patodia S, Thei L, Makwana M, Spencer-Dene B, Latouche M, Mirsky R, Jessen KR, Klein R, Raivich G, Behrens A (2012) c-Jun in Schwann cells promotes axonal regeneration and motoneuron survival via paracrine signaling. J Cell Biol 198:127-141.

[10]Harrisingh MC, Perez-Nadales E, Parkinson DB, Malcolm DS, Mudge AW, Lloyd AC (2004) The Ras/Raf/ERK signalling pathway drives Schwann cell dedifferentiation. Embo j 23:3061-3071.

[11]Ma W, Bisby MA (1998) Increased activation of nuclear factor kappa B in rat lumbar dorsal root ganglion neurons following partial sciatic nerve injuries. Brain Res 797:243-254.

[12]Martini R, Schachner M (1988) Immunoelectron microscopic localization of neural cell adhesion molecules (L1, N-CAM, and myelin-associated glycoprotein) in regenerating adult mouse sciatic nerve. J Cell Biol 106:1735-1746.

[13]Parkinson DB, Bhaskaran A, Arthur-Farraj P, Noon LA, Woodhoo A, Lloyd AC, Feltri ML, Wrabetz L, Behrens A, Mirsky R, Jessen KR (2008) c-Jun is a negative regulator of myelination. J Cell Biol 181:625-637.

[14]Saito H, Kanje M, Dahlin LB (2009) Delayed nerve repair increases number of caspase 3 stained Schwann cells. Neurosci Lett 456:30-33.

[15]Smith D, Tweed C, Fernyhough P, Glazner GW (2009) Nuclear factor-kappaB activation in axons and Schwann cells in experimental sciatic nerve injury and its role in modulating axon regeneration: studies with etanercept. J Neuropathol Exp Neurol 68:691-700.

[16]Sun G, Li Z, Wang X, Tang W, Wei Y (2013) Modulation of MAPK and Akt signaling pathways in proximal segment of injured sciatic nerves. Neurosci Lett 534:205-210.

[17]Tacke R, Martini R (1990) Changes in expression of mRNA specific for cell adhesion molecules (L1 and NCAM) in the transected peripheral nerve of the adult rat. Neurosci Lett 120:227-230.

[18]Thornton MR, Mantovani C, Birchall MA, Terenghi G (2005) Quantification of N-CAM and N-cadherin expression in axotomized and crushed rat sciatic nerve. J Anat 206:69-78.

[19]Woodhoo A, Alonso MB, Droggiti A, Turmaine M, D'Antonio M, Parkinson DB, Wilton DK, Al-Shawi R, Simons P, Shen J, Guillemot F, Radtke F, Meijer D, Feltri ML, Wrabetz L, Mirsky R, Jessen KR (2009) Notch controls embryonic Schwann cell differentiation, postnatal myelination and adult plasticity. Nat Neurosci 12:839-847.

**High-throughput data processing methods**

We retrieved the transcriptome dataset GSE109075 from the GEO database. This dataset profiles transcriptome expression in the sciatic nerves of wild-type mice pre- and post-crush injury, encompassing four injured samples and three uninjured samples. Given that the mouse sciatic nerve FPKM data only provided gene identifiers, we initially performed gene annotation using the clusterProfiler package in R. As the FPKM data had already undergone preliminary processing, no further preprocessing steps were applied prior to downstream analysis.

We first extracted the gene set included in our endogenous network, then subjected the filtered data to z-score normalization to achieve a mean of 0 and variance of 1. Outliers exceeding two standard deviations were adjusted to the corresponding boundary values, and the data were subsequently scaled to the 0–1 interval via Min-Max normalization to facilitate validation against our computational predictions. Given the inherent noise and technical variability associated with high-throughput profiling, a discrepancy of ≤ 0.5 between our computational results and transcriptome-derived values was deemed acceptable. Each repair-type Schwann cell sample was compared with the corresponding steady state of repair-type Schwann cells determined in our analysis. The corresponding results are presented in Figure 3B.

F． **Single-cell Data Analysis Methods**

The single-cell RNA-seq dataset (GSE216665) was obtained from the GEO database and used to characterize Schwann cell states during peripheral nerve injury. Cells were grouped according to the original annotations into Naive, Day 3, Day 12, and Day 60 post-injury samples.

Raw count matrices from multiple samples were imported and merged into a unified Seurat object. Standard preprocessing was performed, including quality control based on sequencing depth, removal of low-quality and outlier cells, and normalization of gene expression data. Cells with extremely low library size or abnormal global expression levels were excluded to improve data robustness.

After normalization, highly variable genes were identified and used for dimensionality reduction. Principal component analysis (PCA) and Uniform Manifold Approximation and Projection (UMAP) were applied to visualize the global transcriptional landscape across developmental stages.

Schwann cells were identified based on canonical marker genes, and major cellular subtypes were defined using established gene signatures, including myelinating, Remak, and repair-associated Schwann cell states. Each cell was assigned to the subtype with the highest module score derived from corresponding marker gene sets.

To compare experimental single-cell states with theoretical model predictions, a gene set corresponding to the Schwann cell endogenous regulatory network constructed in this study was extracted. This gene set was derived from the core network underlying Schwann cell state transitions in our dynamical model. Only genes present in both the scRNA-seq dataset and the model-derived steady state solutions were retained for downstream analysis.

Principal component analysis was then performed on the single-cell expression matrix of the selected genes. Model-derived steady-state expression profiles (M-steady state, NM-steady state, and RSC-steady state) were projected into the same PCA space using the transformation learned from the single-cell data, enabling direct comparison between experimental and simulated states.

Finally, single-cell distributions and model steady states were jointly visualized in the reduced PCA space, allowing assessment of the correspondence between transcriptional cell states and theoretically predicted stable states across different post-injury time points.
